# A catalogue of verified and characterized arterial enhancers for key arterial identity genes

**DOI:** 10.1101/2024.04.30.591717

**Authors:** Svanhild Nornes, Susann Bruche, Niharika Adak, Ian McCracken, Sarah De Val

## Abstract

The establishment and growth of the arterial endothelium requires the coordinated expression of numerous genes. However, the transcriptional and signalling pathways regulating this process are still not fully established, and only a small number of enhancers for key arterial genes have been characterized. Here, we sought to generate a useful and accessible cohort of arterial enhancers with which to study arterial transcriptional regulation. We combined *in silico* analysis with transgenic zebrafish and mouse models to find and validate enhancers associated with eight key arterial identity genes (*Acvrl1*/*Alk1*, *Cxcr4, Cxcl12, Efnb2, Gja4/Cx37, Gja5/Cx40*, *Nrp1* and *Unc5b)*. This identified a cohort of enhancers able to independently direct robust transcription to arterial ECs within the vasculature. To elucidate the regulatory pathways upstream of arterial gene transcription, we determined the occurrence of common endothelial transcription factor binding motifs, and assessed direct binding of these factors across all arterial enhancers compared to similar assessments of non-arterial-specific enhancers. These results find that binding of SOXF and ETS factors is a shared event across arterial enhancers, but also commonly occurs at pan-endothelial enhancers. Conversely, RBPJ/Notch, MEF2 and FOX binding was over-represented but not ubiquitous at arterial enhancers. We found no shared or arterial-specific signature for canonical WNT-associated TCF/LEF transcription factors, canonical TGFβ/BMP-associated SMAD1/5 and SMAD2, laminar shear stress-associated KLF factors or venous-enriched NR2F2 factors. This cohort of well characterized and in vivo-verified enhancers can now provide a platform for future studies into the interaction of different transcriptional and signalling pathways with arterial gene expression.

## INTRODUCTION

The blood vessel system consists of a highly branched network of tubes lined by endothelial cells (ECs) and hierarchically organised into arteries, veins, and capillaries. The EC layer is the first part of the blood vessel to form, initially via differentiation from mesoderm (vasculogenesis) and later from existing ECs (angiogenesis)^1^. While the first arterial ECs arise during vasculogenesis^2^, single cell transcriptomics and fate mapping experiments spanning humans, mice and zebrafish indicate that most arterial ECs form via angiogenesis from venous and venous-like capillary ECs^3–11^. During this process, a subset of ECs reduce cell-cycling and venous gene transcription, induce arterial gene expression and migrate against flow to form into new arteries or extend existing ones. While significant alterations to gene transcription occur during this transition, the precise mix of hardwired signalling pathways and environmental stimuli regulating the differentiation of arterial ECs has been challenging to untangle and identify.

Many components of Notch signalling are selectively expressed in arterial ECs and are essential for arterial formation (e.g. the DLL4 ligand)^12^. However, the assumed model of Notch signalling directly activating arterial gene expression has recently been challenged by new evidence. In particular, retinal and coronary vessels lacking both endothelial MYC (a driver of metabolism and proliferation) and RBPJ (the nuclear effector of Notch signalling) can still express of arterial genes and form arterial structures^13^. This research led to a new paradigm in which Notch drives arterial EC differentiation by reducing metabolism and cell-cycle rather than by directly activating arterial genes^13^. However, cell-cycle changes alone do not necessarily alter arterial identity, and *Myc* loss in retinal EC does not affect arterial patterning^14^. Therefore, Notch-mediated cell-cycle exit likely works alongside other regulators which directly control arterial gene transcription, while the precise role of Notch in direct arterial gene expression remains unclear.

Although numerous other regulatory pathways have been implicated in arterial transcriptional regulation, their exact contribution has been challenging to establish and none appear essential for arterial EC identity^11^. Both canonical WNT and TGFβ/BMP signalling pathways have been implicated in arterialization yet ECs lacking β-catenin or SMAD4 still express arterial genes and form arterial structures^15,16^. Likewise, blood flow is required for full expression of arterial genes, yet arteries in both early embryonic and coronary vasculatures form prior to blood flow^3,6,17,18^. Our knowledge of the transcription factors activating arterial gene expression is also incomplete. ETS factors are required for arterial gene activity but are also essential for vein-specific and pan-endothelial gene expression, suggesting a more general requirement for endothelial identity^19^. The link between FOXC factors and arteries was partially predicated on binding to *Dll4* regulatory regions later found to lack arterial activity^16^. DACH1 potentiates arterial differentiation but is widely expressed and cannot alter EC identity^20^, while arterial-enriched MECOM is linked to repression of venous gene expression rather than activation of arterial identity genes^5^. The evidence linking SOXF transcription factors to arterial differentiation is more extensive, with loss of either SOX17 (the SOXF factor most specific to arterial ECs) or SOX7 resulting in significant arterial defects^21–23^. However, although SOX17 is considered arterial-specific by late fetogenesis, both SOX7 and SOX18 are more widely expressed, and all SOXF factors bind the same motifs^24^. Consequently, the manner in which SOXF factors contribute to the specific activation of arterial genes in still unknown. It is also unclear whether SOXF factors primarily act upstream of Notch signalling (and subsequent cell-cycle related control of arterial differentiation), or whether they more widely activate arterial gene expression. Direct SOXF binding is best characterized at Notch pathway enhancers *Dll4in3*, *Dll4-12* and *Notch1+16,* and the arterial defects seen after *Sox17* deletion were attributed to a requirement for SOX17 in activation of *Notch1* and *Dll4*^23,25,26^. However, SOXF motifs are also required for the activity of the arterial-specific *ECE1in1* enhancer and are associated with coronary arterial *Nestin* expression^27,28^.

In this paper, we combine an *in silico* enhancer search with verification in transgenic models to identify a cohort of arterial enhancers associated with eight key arterial identity genes. We then use sequence analysis and DNA-protein binding surveys to investigate the involvement of many endothelial- and arterial-associated transcription factors in arterial enhancer binding, and to compare this pattern with that seen at pan-endothelial and venous enhancers. Our results indicate that ETS and SOXF factors play a general role in endothelial gene transcription, support a role for FOX factors more selectively in arterial activation, and link both RBPJ and MEF2 factors to a limited number of arterial genes. This cohort of well characterized, in vivo-verified enhancers, can also now be used by others as platform for future studies into the interaction of different transcriptional and signalling pathways with specific arterial genes and with subtype-specific gene expression within the endothelium more generally. Additionally, our data provides a useful training set for attempts to more accurately classify endothelial enhancers genome-wide.

## METHODS

### Animals

All animal procedures were approved by a local ethical review committee at Oxford University and licensed by the UK Home Office. F0 mosaic transgenic zebrafish embryos were generated using Tol2 mediated integration^29^. The F1 stable transgenic lines were generated from an initial outcross of adult F0 carriers. An intercross of adult F1 lines produced F2 lines. To enable visualization of the entire vasculature the adult F1 transgenic lines were intercrossed with the *tg(kdrl:HRAS-mCherry)* zebrafish line. Embryos were maintained in E3 medium (5 mM NaCl; 0.17 mM KCl; 0.33 mM CaCl2; 0.33 mM MgSO4) at 28.5°C. Some of the embryos were incubated at 30°C to 32°C to modify the speed of embryonic development. To image, all embryos were dechorionated and anesthetized with 0.01% Tricaine methanesulfonate in E3 medium. For analysis of transgenic zebrafish, single embryos were transferred into a flat bottom 96-well plate, mounted in 0.1% TopVision low melting point agarose (Thermo Fisher Scientific, R0801) in E3 medium with tricaine methanesulfonate and 0.003% phenylthiourea (Merck, P7629) to inhibit pigmentation. GFP and mCherry reporter gene expression was screened with a Zeiss LSM 980 (Carl Zeiss) confocal microscope at 32-72 hpf. Whole zebrafish were imaged using the ‘tile scan’ command, combined with Z-stack collection, under the confocal microscope Zeiss LSM 980 at 488 nm excitation and 510 nm emission for EGFP, and at 561 nm excitation and 610 nm emission for mCherry. The eyes of the zebrafish were imaged similarly, but without tile scanning.

E14.5 F0 transgenic mouse embryos were generated, dissected and stained in X-gal by Cyagen Biosciences. Yolk sac was collected separately and used for genotyping. All embryos were imaged using a Leica M165C stereo microscope equipped with a ProGres CF Scan camera and CapturePro software (Jenoptik). For each enhancer, embryos were also sectioned for histological analysis to investigate X-gal staining patterns. For histological analysis, embryos were dehydrated through a series of ethanol washes, cleared by xylene and paraffin wax-embedded. 5 or 6-μm sections were prepared, de-waxed, and counterstained with nuclear fast red (Electron Microscopy Sciences).

### Cloning

All enhancer sequences were generated as custom-made, double-stranded linear DNA fragments (GeneArt® Strings™, Life Technologies). The sequences of all enhancers are provided in the Supplementary Methods section. DNA fragments containing the enhancer sequences were cloned into the pCR8 vector using the pCR8/GW/TOPO TA Cloning Kit (Thermo Fisher Scientific, K250020) following the manufacturer’s instructions. Once cloning was confirmed, the enhancer sequence was transferred from the pCR8/GW/enhancer entry vector to a suitable destination vector using Gateway LR Clonase II Enzyme mix (Thermo Fisher Scientific, 11791100) following the manufacturer’s instructions. For zebrafish transgenesis, the enhancer was cloned into the E1b-GFP-Tol2 vector (provided by N. Ahituv). For mouse transgenesis, the enhancer was cloned into the hsp68-LacZ-Gateway vector (provided by N. Ahituv).

### Transcription factor binding assays

With the exception of SOX7 and SOX17, all transcription factor binding data was previously published, and was assessed in IGV^30^ either through downloading from GEO or via ChIP Atlas ^31^. In every case, we first verified the correct data was accessed by reproducing images at loci used in the primary publication, and highly recommend this practice for others as errors inevitably occur during data deposition. ERG and NR2F2 binding data in HUVECs (GSM3673462 and GSM3673452) came from ^32^, ETS1 binding data in HUVECs after 12 hours of VEGFA stimulation (GSM2442778 SRX2452430) came from^33^. FOXO1 binding data in HUVECs (GSM3681485/6 SRX5548892) came from^34^ and FOXO1 binding in adult untreated mouse hearts (GSM4278011) came from^35^. RBPJ binding data in HUVECs after 12 hrs of VEGFA stimulation (GSM2947456 SRX3599311) came from ^36^. SMAD1/5 binding in HUVECs after BMP9 stimulation (GSM684747 SRX045541) was from^37^, SMAD2 in HUVECs after TGFβ stimulation (GSM3955796 SRX6476491) came from^38^. MEF2C binding data in HUVECs (GSM809016 SRX100256) came from^39^, MEF2A binding data in mouse adult hearts (GSM3518665 SRX5146757) came from ^40^.

For SOX7 and SOX17 CUT&RUN, HUVECs were cultured in EBM-2 basal medium (Lonza CC-3156) supplemented with EBM-2 SingleQuot supplement and growth factor kit (Lonza CC-4176). DNA binding assays were performed using the CUT&RUN Kit from Cell Signaling Technologies (CST 86652) following the manufacturer’s protocol with slight modifications. In brief, harvested cells were lightly crosslinked with 0.1% formaldehyde for 2min, rinsed with wash buffer supplemented with 1% Triton X-100 and 0.05% SDS, bound to Concanavalin A beads and incubated overnight with antibodies against IgG control (CST 66362), SOX7 (R&D Systems, AF2766) or SOX17 (R&D Systems, AF1924) in wash buffer containing 0.05% digitonin. DNA around binding sites was cleaved with pAG-MNase enzyme and the released DNA was reverse-crosslinked with proteinase K and 0.1% SDS overnight at 65°C before purification with a ChIP DNA Clean & Concentrator kit (Zymo Research D5205). DNA was converted into Illumina-compatible libraries with the NEBNext(R) Ultra(TM) II DNA Library Prep Kit (NEB E7645L) following the protocol described by ^41^ and using NEBNext Dual Index Multiplex Oligos (NEB E7600S). Libraries were sequenced on a NextSeq2000 (SOX17) or a NovaSeq (SOX7) using paired end reads. Data was processed using the nf-core/cutandrun v3.1 pipeline (10.5281/zenodo.5653535^42^) with the following adjustments to the default settings: --normalisation_mode CPM and -- trim_nextseq 20. The CUT&RUN hg38 blacklist^43^ was used during sequence alignment. Peak calling was performed with SEACR^44^ using stringent settings, and by HOMER^45^ using default settings for transcription factors (-style factor). Motif analysis was performed with HOMER using 200nt regions around peak centres for SOX7 and SOX17 or the full validated region of the arterial enhancers. Overlap of SOX7 and SOX17 peaks with published mCherry-SOX7 data^46^, HUVEC enhancer marks and TSS^32^ was executed using the vennRanges R package. Data has been deposited to ArrayExpress under the accession number E-MTAB-13999.

### Re-analysis of scRNA-seq data

Publicly available E12 and E17.5 scRNA-seq data from EC isolated from *BmxCreERT2;Rosa^tdTomato^* lineage traced murine hearts^47^ was obtained from GEO (GSE214942) prior to processing FASTQ files with the 10X Genomics CellRanger pipeline (V7.0.0) and filtering out low quality cells using Scater^48^. Data normalisation was performed using the MultiBatchNormalisation method prior to merging of *TdTomato* positive and negative datasets from individual timepoints. Detection of highly variable genes, principal component analysis, non-supervised clustering, and UMAP visualisation were conducted using the standard Seurat (V4) workflow^49^.

## RESULTS

Transcription factors primarily regulate endothelial gene transcription through binding to enhancers (*cis*-regulatory elements)^1^. Consequently, analysis of enhancer sequences can elucidate the precise combination of transcription factors, and cognate upstream signalling pathways, involved in different patterns of gene expression. One of the main challenges in understanding arterial regulation has been a paucity of characterized enhancers for key arterial genes. For example, of the 16 genes used to define mouse coronary arterial EC identity in single cell transcriptomics^20^, only four have *in vivo*-verified enhancers (*Dll4*, *Hey1*, *Notch1* and *Acvrl1*). Three of these are genes in the Notch pathway and are either self-regulated by Notch/RBPJ (*Dll4-12* and *Hey1-18*)^25,50^ or lack specificity during early coronary arterial specification (*Notch1+16*)^51^. The fourth enhancer, for *Acvrl1*, is of a size (9kb) that precludes analysis^52^. Beyond this, there are only four other *in vivo*-validated arterial enhancers described in the literature, for *Ece1, Flk1, Sema6d* and *Sox7*. Of these, only the *Ece1in1* and *Flk1in10* arterial enhancers, both genes not specific to arterial ECs, have been analysed at the level of transcription factor binding^28,53^. It is therefore clear that a better understanding of the regulatory pathways directing arterial differentiation requires the identification and characterization of a larger number of arterial enhancers directing the expression of key arterial identity genes. To identify a cohort of such enhancers, we looked in the loci of eight non-Notch genes: *Acvrl1*(ALK1) *Cxcr4, Cxcl12, Efnb2, Gja4*(CX37)*, Gja5* (CX40), *Nrp1* and *Unc5b.* These genes are all significantly enriched in arterial ECs, and are commonly used to define arterial EC populations in mouse and human scRNAseq analysis^4,5,20,54^.

To identify putative enhancers *in silico,* we used five published datasets detailing different enhancer-associated chromatin marks: (i) open chromatin as assessed by ATAC-seq in primary mouse adult aortic ECs (MAECs) from Engelbrecht et al^55^; (ii) open chromatin as assessed by ATAC-seq in mouse postnatal day 6 (P6) retina ECs (MRECs) from Yanagida et al^56^; (iii) enriched EP300 binding in Tie2Cre+ve cells from embryonic day (E)11.5 mouse embryos (from Zhou et al^53^); (iv) enriched H3K27Ac and/or H3K4Me1 in human umbilical vein ECs (HUVECs, data available on the UCSC genome browser^57^); and (v) open chromatin regions assessed by DNAseI hypersensitivity in HUVECs and dermal-derived neonatal and adult blood microvascular ECs (HMVEC-dBl-neo/ad) comparative to non-ECs (UCSC genome browser^57^). A retrospective analysis of 32 previously described mammalian *in vivo*-validated EC enhancers^1^, which included eight arterial enhancers, found that 31/32 were marked by at least one enhancer mark in both human and mouse samples (including 8/8 of arterial enhancers), see Table S1. We next analysed the loci of our target arterial genes to identify putative enhancers using these enhancer marks (Figure S1). For arterial genes robustly transcribed in both human and mouse EC datasets (determined by open chromatin/H3K4Me3 at the promoter region), we defined a putative enhancer as a region containing at least one enhancer mark in both mouse and human ECs. Because *Cxcr4*, *Cxcl12*, and *Gja5* were poorly transcribed in the human cell lines studied here, for these genes the putative enhancer definition was relaxed to include regions containing two enhancer marks in mouse ECs with no marks in human cells. Each putative enhancer was named according to their neighbouring arterial gene and distance from the transcriptional start site (TSS) in mice (e.g. the putative Efnb2-112 enhancer is 112kb upstream of the mouse Efnb2 TSS). In total, this analysis identified 39 putative enhancers for further testing (Table S2). Our initial analysis included assessment of seven regions previously identified as potential enhancers for *Efnb2*, *Nrp1* and *Cxcr4* but whose independent activity was never validated *in vivo*^58–61^ (marked in grey on Table S2). None of these met our putative enhancer threshold: two of these regions were associated with no enhancer marks, three had a single enhancer mark in HUVECs, one had non-specific enhancer marks in human cells only (Nrp1+76/NRP1A^62^), and one contained enhancer marks in human ECs only (Cxcr4-117/CXCR4+125^60^).

### Transgenesis in zebrafish confirmed vascular activity of a subset of these putative enhancers

It is well established that regions associated with enhancer marks do not necessarily act as enhancers by independently activating gene transcription. Out of the 39 putative enhancers identified here, only three had been previously investigated *in vivo*: Nrp1+28, which was able to drive robust pan-endothelial expression of the *lacZ* reporter gene in transgenic mouse embryos^63^; and Acvrl-5 and Acvrl1-1/p, which were silent in a similar mouse assay^52^. Additionally, Acvrl1+6 is contained within a nine kilobase region known to direct arterial-specific expression of *lacZ* in transgenic mice^52^. However, because of the size of this original piece, we treated the smaller Acvrl1+6 enhancer as untested.

To establish the ability of the 36 untested putative enhancers to drive arterial EC activity, we analysed the activity of each enhancer in F0 Tol2-mediated mosaic transgenic zebrafish embryos^29^. Similar zebrafish assays have been conducted with five of the eight previously identified arterial enhancers, and in each case these mammalian enhancer sequences were able to drive GFP expression in arterial ECs in zebrafish embryos (Fig. S2 and ^25,64–66)^. Here, the mouse sequence of each of the 36 putative arterial enhancers was cloned upstream of the E1b minimal promoter and GFP reporter gene and used to generate transgenic embryos, which were examined at 2 days post fertilization (dpf)^29^. This analysis identified 18 enhancers able to drive GFP expression in ECs of transgenic zebrafish, defined as EC GFP expression in more than 5% of injected embryos (Table 1 and Figure 1). This included at least one enhancer for each of our eight target genes. Of the remaining 18 putative enhancers, none were able to consistently drive detectable GFP expression in ECs (Table 1).

**Figure 1.**
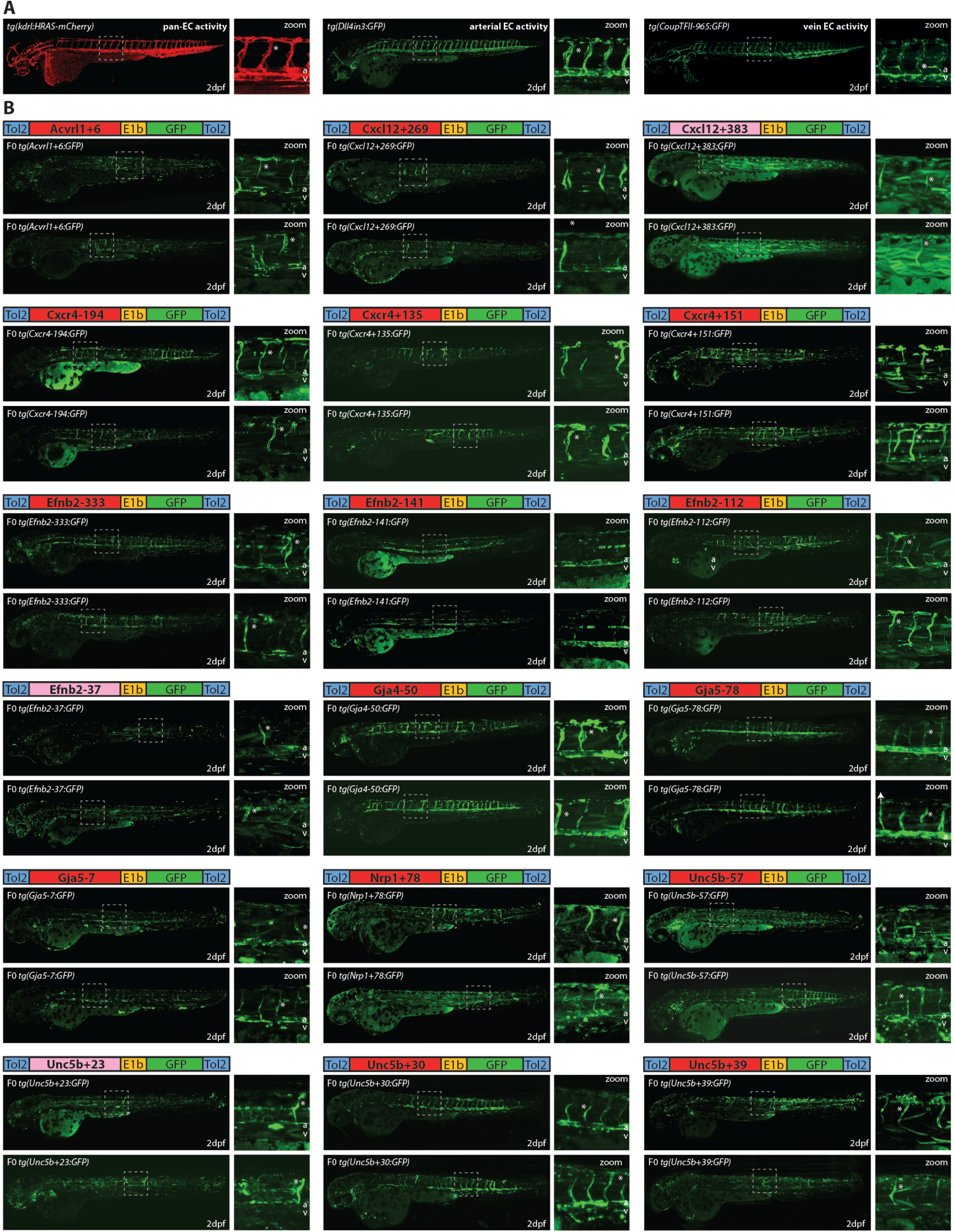
Analysis of putative enhancers in F0 mosaic Tol2 transgenic zebrafish identifies 18 enhancers able to drive GFP activity in arterial ECs. **A** Examples of the expression of known pan-EC (kdrl1:HRAS-mCherry)^87^, arterial (Dll4in3:GFP^25^) and vein (CoupTFII-956:GFP^15^) enhancers in 2dpf zebrafish. **B** Two representative F0 transgenic zebrafish expressing each of the 18 new arterial enhancers alongside a schematic of each transgene. Enhancers in red represent robust arterial expression, pink represents weaker enhancers with limited expression in a lower percentage of injected zebrafish, see also Table 1. Grey dashed box indicates region of zoom, a=dorsal aorta, v=cardinal vein, * indicates intersegmental vessels.

**Table 1.** Summary of putative enhancer activity in mosaic Tol2 transgenic zebrafish. “In vivo classification” indicates the results of this screen, “in vitro classification” indicates previous designation in ^32^, as defined by relative enhancer and promoter marks in HUVECs vs HUAECs. * indicates only limited expression in a very small number of ECs. Acvrl1-5 and Acvrl1-1/p were previously investigated by ^85^, Nrp1+28 by ^63^, Notch1+16 by ^26^, Dll4-12 and Dll4in3 by ^25^, Flk1in10 by ^64^, Sema6d-55 by ^53^, Sox7+14 by ^53,66^, Ece1in10 by ^28^ and Hey1-18 by ^50^. ** After this analysis but prior to this publication, Gja5-21 has been shown to direct expression in the zebrafish endocardium at 4 dpf ^86^.

| Enhancer | # Injected/# any EC GFP | In vivo classification | In vitro classification |
| --- | --- | --- | --- |
| Acvrl1-5 | Seki *et al* 2004 | inactive | pan-EC enhancer |
| Acvrl1-1/p | Seki *et al* 2004 | inactive | pan-EC TSS |
| **Acvrl1+6** | 127/49 | **arterial enhancer** | pan EC TSS |
| Acvrl1+16 | 205/0 | inactive | pan EC TSS |
| Acvrl1+19 | 95/0 | inactive | pan EC TSS |
| Cxcl12-184 | 64/0 | inactive | pan-EC enhancer |
| Cxcl12-2 | 32/0 | inactive | pan-EC enhancer |
| Cxcl12+239 | 70/1 | inactive | inactive |
| Cxcl12+265 | 42/1 | inactive | inactive |
| **Cxcl12+269** | 163/63 | **arterial enhancer** | inactive |
| Cxcl12+298 | 51/0 | inactive | pan-EC enhancer |
| Cxcl12+376 | 149/0 | inactive | inactive |
| **Cxcl12+383** | 152/37* | **weak enhancer\*** | inactive |
| Cxcl12+439 | 73/0 | inactive | pan-EC enhancer |
| Cxcl12+445 | 145/0 | inactive | pan EC TSS |
| Cxcr4-232 | 163/0 | inactive | inactive |
| **Cxcr4-194** | 46/25 | **arterial enhancer** | inactive |
| Cxcr4-130 | 95/0 | inactive | inactive |
| Cxcr4-117^ | 209/9* | inactive | inactive |
| *hCxcr4-117^/CXCR4-125* | 81/0 | inactive | inactive |
| Cxcr4-113 | 300/0 | inactive | pan-EC enhancer |
| Cxcr4-109 | 89/0 | inactive | pan-EC enhancer |
| Cxcr4+1 | 33/0 | inactive | inactive |
| **Cxcr4+135** | 69/56 | **arterial enhancer** | inactive |
| **Cxcr4+151** | 96/50 | **arterial enhancer** | arterial TSS |
| **Efnb2-333** | 152/93 | **arterial enhancer** | pan-EC enhancer |
| **Efnb2-141** | 74/36 | **arterial enhancer** | arterial TSS |
| **Efnb2-112** | 65/30 | **arterial enhancer** | arterial TSS |
| Efnb2+3 | 92/0 | inactive | pan EC TSS |
| **Efnb2+37** | 114/18* | **weak enhancer\*** | pan-EC enhancer |
| Efnb2+172 | 63/0 | inactive | pan-EC enhancer |
| Efnb2+209 | 158/0 | inactive | pan-EC enhancer |
| Gja4+24 | 52/0 | inactive | inactive |
| **Gja4+50** | 232/187 | **arterial enhancer** | pan-EC enhancer |
| *Gja4+57* | 192/0 | inactive | inactive |
| **Gja5-7** | 66/39 | **arterial enhancer** | arterial enhancer |
| *Gja5-21\*\** | 38/0 | inactive | pan-EC enhancer |
| Gja5-28 | 156/0 | inactive | inactive |
| **Gja5-78** | 76/62 | **arterial enhancer** | pan-EC enhancer |
| Gja5-93 | 253/0 | inactive | arterial enhancer |
| **Nrp1+28** | De Val *et al* 2008 | **pan-EC enhancer** | pan-EC enhancer |
| *Nrp1+76* | 54/0 | inactive | pan-EC enhancer |
| **Nrp1+78** | 191/34 | **arterial enhancer** | pan-EC enhancer |
| Nrp1+91 | 109/0 | inactive | pan-EC enhancer |
| Nrp1+129 | 110/0 | inactive | pan-EC enhancer |
| **Unc5b-57** | 50/21 | **arterial enhancer** | inactive |
| *Unc5b+14* | 61/0 | inactive | inactive |
| **Unc5b+23** | 82/16* | **weak enhancer\*** | arterial enhancer |
| **Unc5b+30** | 96/79 | **arterial enhancer** | arterial enhancer |
| **Unc5b+39** | 96/56 | **arterial enhancer** | arterial enhancer |
| *Unc5b+43* | 111/0 | inactive | arterial enhancer |
| **NOTCH1+16** | Chiang *et al* 2017 | **arterial enhancer** | pan-EC enhancer |
| **DLL4-12** | Sacilotto *et al* 2013 | **arterial enhancer** | pan-EC enhancer |
| **DLL4in3** | Sacilotto *et al* 2013 | **arterial enhancer** | pan-EC enhancer |
| **Flk1in10** | Becker *et al* 2016 | **arterial enhancer** | inactive |
| **Sema6d-55** | Zhou *et al* 2017 | **arterial enhancer** | pan-EC enhancer |
| **ECE1in1** | Robinson *et al* 2014 | **arterial enhancer** | pan-EC enhancer |
| **SOX7+14** | Zhou *et al* 2017 | **arterial enhancer** | pan-EC enhancer |
| **Hey1-18** | Watanabe *et al* 2020 | **arterial enhancer** | pan-EC enhancer |

In addition to the 38 putative enhancers identified through our screen, we tested an additional 11 regions that fell below our enhancer threshold. This included regions with enhancer marks in mice only or human only, regions with only one enhancer mark, and the two regions previously implicated as enhancers (Nrp1+76 and Cxcr4-117^60^). Of these 11 regions, only Cxcr4-117 was able to drive any detectable expression in ECs (Table 1). However, this was seen in only 9 of 209 injected embryos (<5%) and was limited to 1-2 ECs in each F0 fish (see Figure S3). A human version of this enhancer, CXCR4-125 (identified as an enhancer in^60^), was also tested but was not able to drive any detectable GFP expression (Table 1), and this enhancer region was therefore not further analysed.

We compared our results to a published genome-wide classification of EC regulatory elements, which used assessment of relative H3K27ac and acEP300 occupancy in freshly isolated HUVEC and HUAECs to classify regions as arterial-enriched, venous-enriched and common (arterial and venous enriched) regulatory elements^32^. Only 4/18 of our in vivo-validated enhancers were classified as arterial enhancers using this assay, with 5/18 classified as common enhancers, 4/18 characterized as TSS and 5/18 uncalled (Table 1). A similar analysis of the eight previously identified arterial enhancers found none were classified as arterial enhancers (7/8 were marked as common enhancers and 1/8 was uncalled) (Table 1). Conversely, 9/18 putative enhancers that were inactive in our transgenic assays were classified as enhancers in the *in vitro* assay (8 pan-EC, 1 arterial) (Table 1). The low correlation between predicted activity and behaviour in transgenic assays suggest that *in vitro* assessments using enhancer marks in cultured ECs alone may not strongly predict the ability of a putative enhancer region to independently direct gene expression nor the resultant specificity of this expression.

### Arterial expression of enhancers in transgenic models

Of the 18 enhancers identified here, 15 enhancers drove robust GFP expression that could be assessed for arterial-venous specificity. Analysis of the trunk vessels in F0 transgenic zebrafish at 2 days post fertilization (dpf) found strong evidence that all 15 of these robust enhancers drove preferential expression of GFP in arterial ECs (Figure 1). 14/15 of these enhancers showed GFP expression in ECs of the dorsal aorta, with the outlier (Cxcr4+135) driving GFP expression only to intersegmental vessels (ISVs) in a pattern indicative of intersegmental arteries (ISArt). In comparison, very little GFP expression was seen in the axial veins (e.g. cardinal and ventral vein). The remaining three enhancers were able to drive only weak GFP or were limited to a small number of ECs in all analysed zebrafish, such that expression pattern within the vasculature could not be reliably analysed and stable transgenic lines could not be generated. These were labelled weak enhancers and no further analysis was conducted.

Because Tol2-mediated mosaic transgenesis gives a variable snapshot of enhancer activity, we also generated stable zebrafish transgenic lines with 14 of our robust enhancers. Analysis of these stable enhancer:GFP zebrafish found very similar patterns of expression relative to the mosaic F0 results, and confirmed arterial activity of all enhancers (Figure 2 and 3). Where enhancer activity was robustly detected in the main axial vessels of the trunk (12/14), GFP expression was overwhelmingly enriched in the ECs of the dorsal aorta comparative to the cardinal and ventral veins. Of the two enhancers active primarily in the intersegmental vessels of the trunk, GFP expression was restricted to intersegmental arteries by 3 dpf, as determined by direct connections to the dorsal aorta (Figure 2 and 3A). A similar enrichment for transgene expression in intersegmental arteries versus veins was seen with the majority of the arterial enhancers active in the dorsal aorta, although the breadth of enhancer activity in the intersegmental vessels varied between enhancers (Figure 2 and 3A). Analysis of specificity within the intersegmental vessels is complicated by the fact that these vessels form by angiogenic sprouting from the dorsal aorta. In addition to any issue of GFP half-life, many of the arterial genes investigated here are also expressed during angiogenic sprouting (e.g. *Efnb2*, *Cxcr4*, *Dll4*, *Nrp1* and *Sox7).* Consequently, GFP expression seen during early intersegmental sprouting may reflect angiogenic activity, explaining the relatively widespread GFP expression of a number of these enhancers in intersegmental vessels at 2 dpf (Figure 2). As such, these elements are potentially both arterial and angiogenic enhancers, similar to that of the well-defined Dll4in3 enhancer^25,67^.

**Figure 2.**
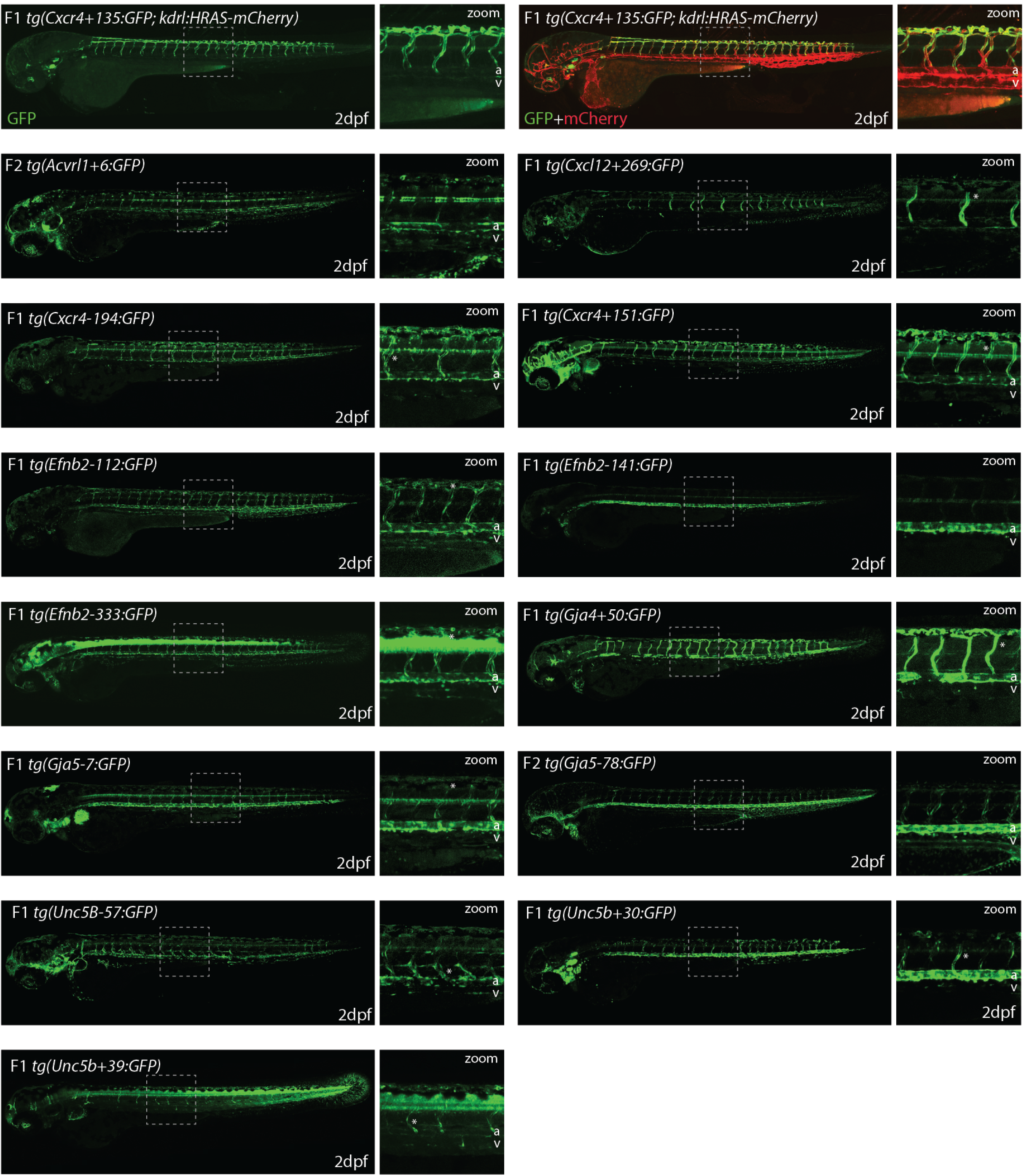
Stable transgenic zebrafish expressing 14 different putative enhancer:GFP transgenes demonstrate arterial-enriched expression patterns. Grey dashed box indicates region of zoom, a=dorsal aorta, v=cardinal vein, * indicates intersegmental vessels. *tg(Cxcr4-135:GFP)* was crossed with *tg(kdrl:HRAS-mCherry),* which expresses mCherry in all blood vascular ECs and is shown here on the top line as a guide to vessel structure at this timepoint. F1 indicates that embryos come from parents from the F1 generation, F2 indicates that embryos come from parents from the F2 generation.

**Figure 3.**
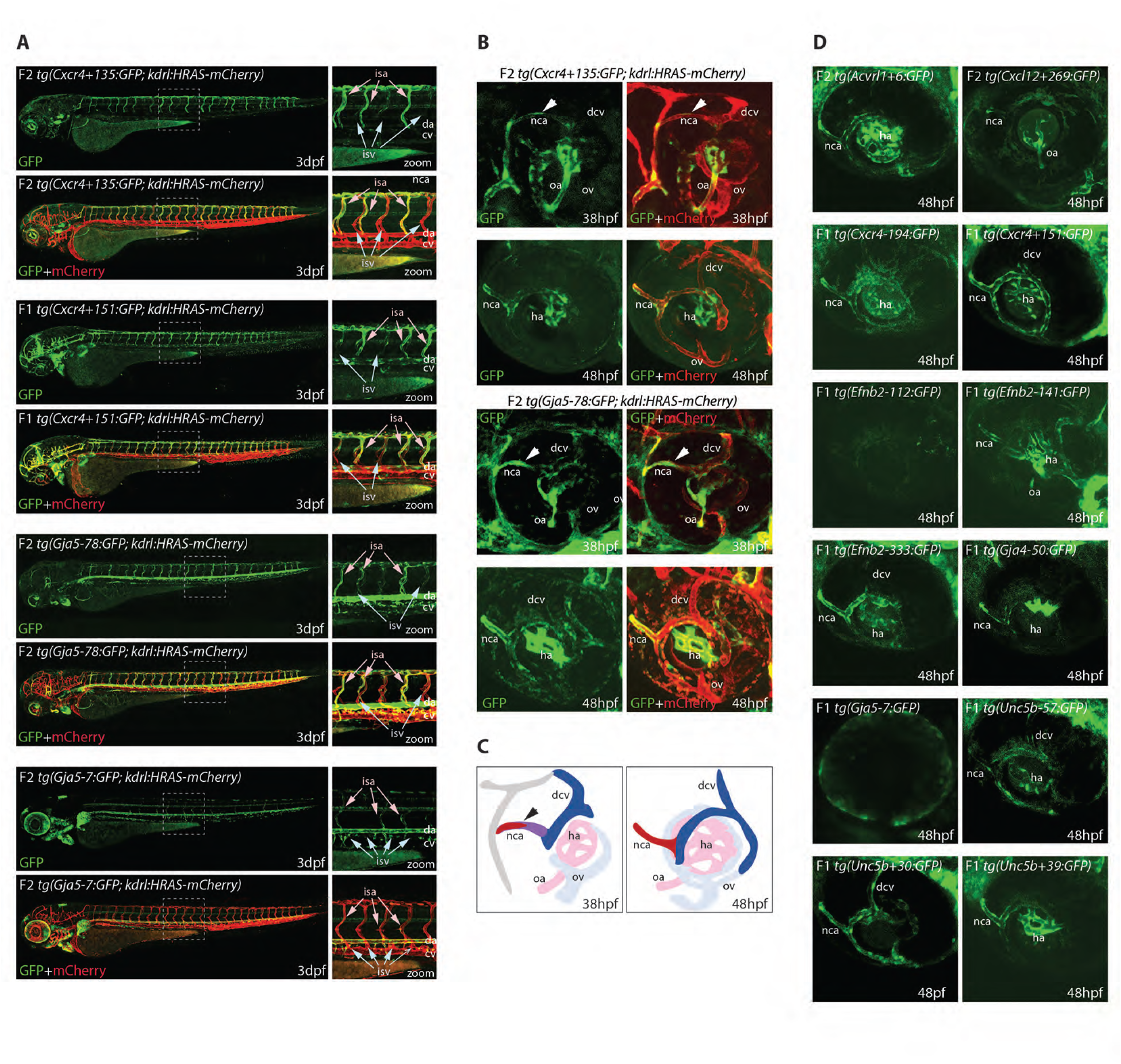
Analysis of arterial expression patterns of select arterial enhancers, some also transgenic for the pan-EC expressed kdrl1:HRAS-mCherry. **A** Enhancer:GFP expression patterns in intersegmental vessels at 3dpf relative to the pan-EC kdrl1:HRAS-mCherry indicates enhancer activity is restricted to vessels connecting to the dorsal aorta. **B-D** Analysis of enhancer activity in the ocular vasculature from 38-48 hpf, where the ECs that form the nasal ciliary artery (nca) sprout from the dorsal ciliary vein (dcv) (as shown in schematic **C**). Activity of arterial enhancers can be seen in the nasal ciliary artery soon after the initial sprout is formed at 38hpf (**B-C),** and 11/14 arterial enhancer lines investigated showed GFP expression in the nasal ciliary artery at 48 hpf **(B and D).** Grey dashed box indicates region of zoom, arrowhead points to the sprout forming the new nca, isa=intersegmental artery, isv=intersegmental veins, da= dorsal aorta, cv=cardinal vein, nca=nasal ciliary artery, dcv=dorsal ciliary vein, oa=optic artery, ov=optic vein, ha= hyaloid artery.

Arterial development in the zebrafish trunk at the timepoints investigated primarily occurs via vasculogenesis and arterial-to-arterial sprouting, as opposed to the vein/capillary origin of many mammalian arterial ECs^3,4,68^. In order to investigate enhancer activity during vein-to-arterial EC differentiation, we also looked at the zebrafish ocular vasculature, where venous ECs from the dorsal ciliary vein (DCV) sprout to become arterial ECs and form the nasal ciliary artery (NCA)^9,69,70^. Analysis of the eyes of our arterial enhancer:GFP lines found that 11/14 of our arterial enhancers were also able to drive GFP activity in the newly formed NCA (Figure 3B-C).

We next investigated whether these enhancers directed similar patterns of activity in transgenic mice, selecting five arterial enhancers active in zebrafish for further analysis. In each case, the enhancer was able to drive activity of the *lacZ* reporter gene in arterial ECs but not venous ECs in E14.5 F0 transgenic mouse embryos (Figure 4). We additionally tested one enhancer which was only weakly active in transgenic zebrafish (Efnb2-37). No endothelial activity was seen for this enhancer in transgenic mice (Figure S4). In combination, this transgenic zebrafish and mouse analysis indicates that we have successfully identified a cohort of enhancers directing gene expression to arterial ECs, accurately reflecting the expression of their cognate genes in mammals.

**Figure 4.**
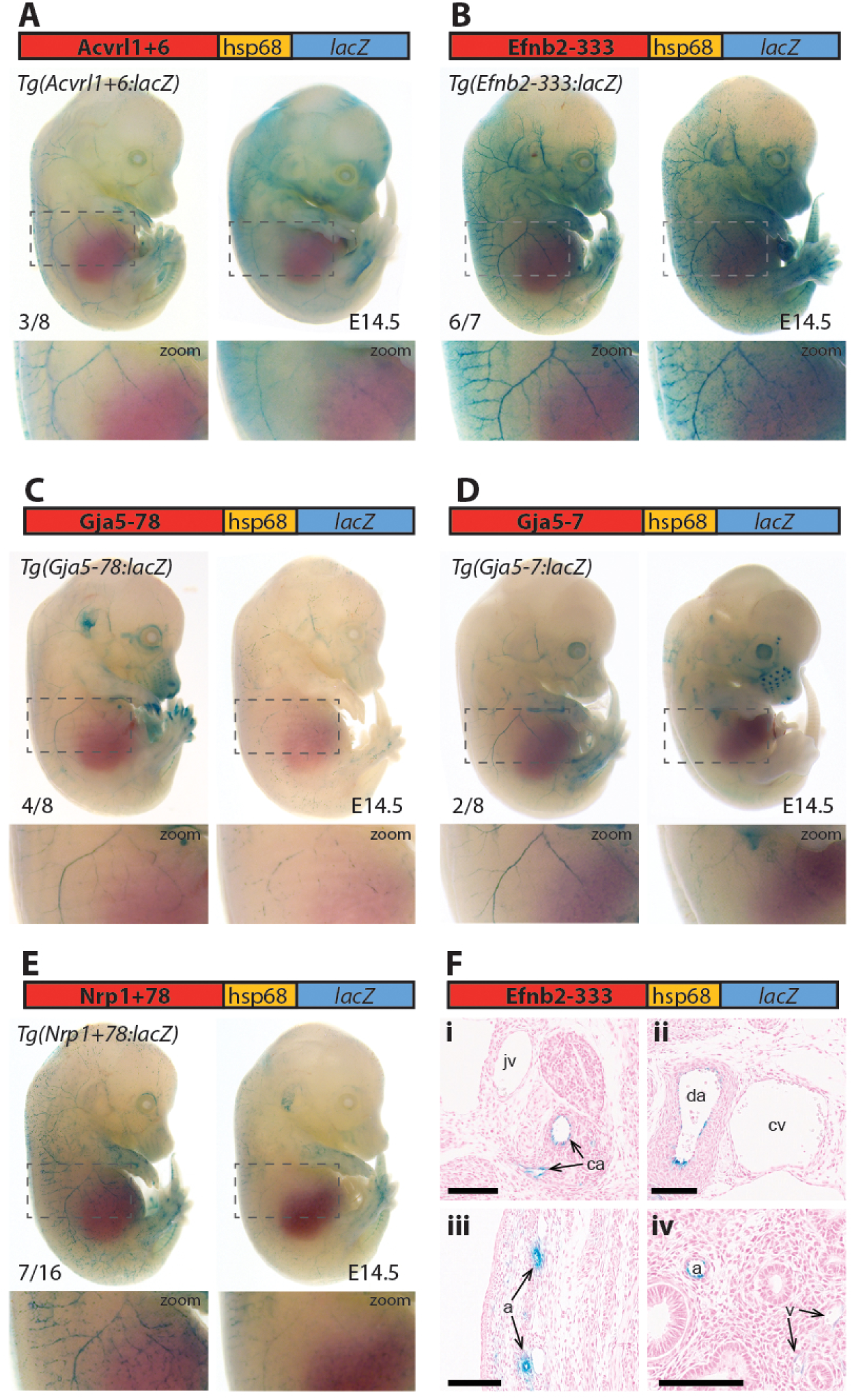
Five putative enhancers direct arterial expression of reporter genes in the vasculature of E14.5 transgenic mouse embryos. **A-E** two representative embryos expressing each tested putative enhancer alongside a schematic of the transgene. Numbers in bottom left indicate embryos with arterial lacZ/total transgenic embryos. Grey dashed boxes indicate region in zoom. **F** shows transverse sections through Efnb2-333:lacZ transgenic E14.5 transgenic embryo from region in head level to eye (**i**), region above heart (**ii**), body wall level with liver (**iii**) and lung (**iv**). ca=carotid artery, jv=jugular vein, da=dorsal aorta, cv=cardinal vein. Black line = 100um

### Assessment of transcription factor motifs and binding patterns at arterial enhancers

This work so far has identified a cohort of enhancers able to drive gene expression in arterial ECs, and to stay silent in venous ECs. We next investigated whether this common expression pattern was associated with the binding of particular transcription factors. The ability of a transcription factor to bind an enhancer or promoter sequence is commonly established by identifying one or more binding motif(s) within the region of interest, and/or by observing direct binding to the region of interest by chromatin immunoprecipitation (ChIP) or similar methodologies. Here, we combined both approaches. First, we performed HOMER analysis to identify overrepresented motifs within the core regions of all 15 arterial enhancers identified here (≈250-400bp, centred on enhancer marks and cross-species conservation) alongside all eight previously identified arterial enhancers (a total of 23). This HOMER motif analysis indicated the repeated presence of motifs for ETS (including EC-associated factors ETS1, ERG and ETV2), SOX (including EC-associated SOX17 and SOX7), FOX (including EC-associated FOXO1 and FOXO3), RBPJ (the transcriptional effector of NOTCH signalling) and MEF2 (including EC-associated MEF2A)( Figure S5). We next directly searched the sequences of all 23 core arterial enhancers to accurately determine the frequency of motifs for each of these transcription factors. In addition, we also looked for possible binding of other transcription factors previously implicated in arterial gene expression. The enhancer motif search used a combination of the JASPAR Transcription Factors Track Settings (TFTS) on the UCSC Genome Browser and hand annotation using previously defined consensus motifs (Figure S6). In parallel, we compared this analysis with a variety of published endothelial ChIP-seq and CUT&RUN datasets to determine where there was evidence of direct binding for each of these transcription factors at each enhancer (see Table 2, Figure 5-6 and Figures S6-9).

**Figure 5.**
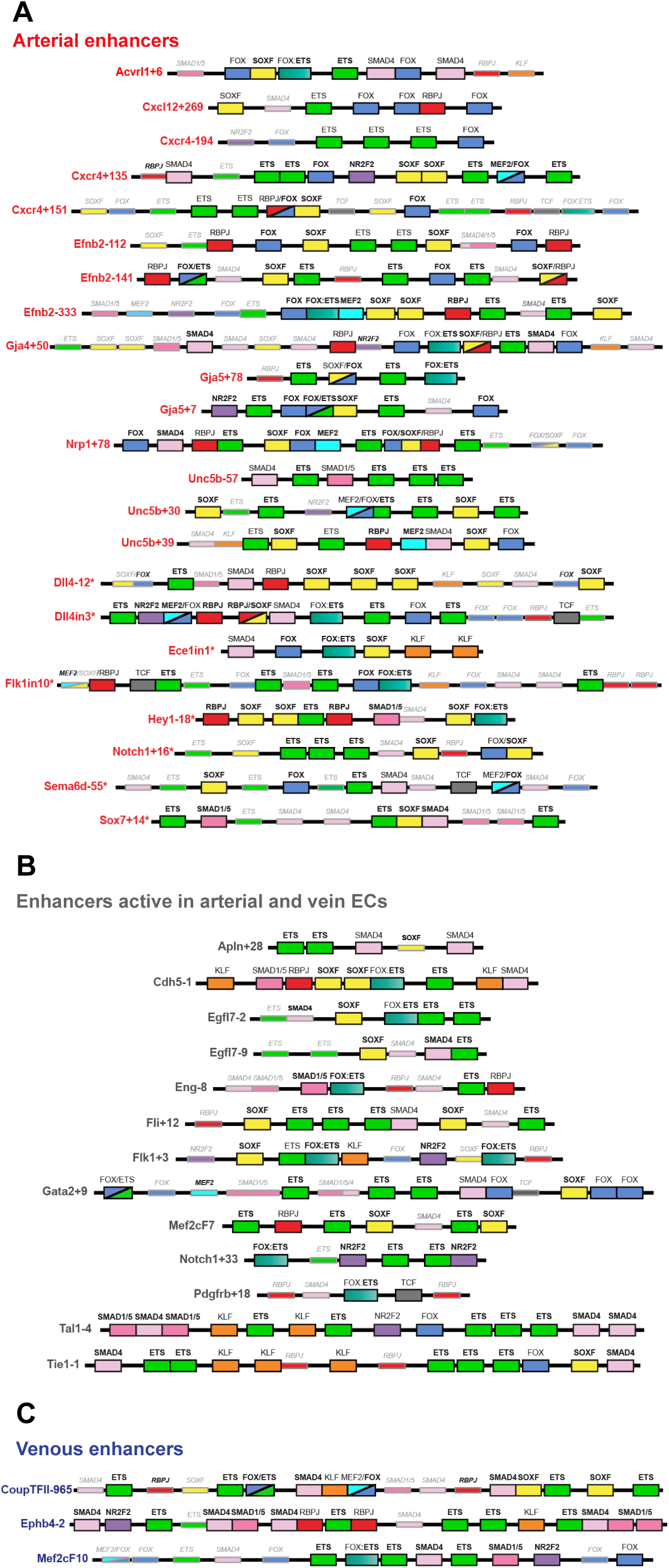
Schematics summarizing the transcription factor motifs found within each arterial (**A**), pan-EC (**B**) and venous (**C**) enhancer. Deep black-lined boxes indicate motifs for transcription factors conserved at the same depth as the surrounding enhancer sequence, shallow grey-lined boxes/text indicate motifs conserved between mouse and human enhancer sequence but not at the same depth as the surrounding sequence. **Bold** transcription factor names indicate places where ChIP-seq (or similar analysis) confirms binding. See Figure S6 for motif information, S8 for annotated sequences and Figures 6, S7 and 9 for IGV visualization of transcription factor binding patterns. Arterial enhancers listed with * are previously published (as detailed in ^1^), genome locations for each enhancer are provided in Table S3. Distances between motifs are representative but not scaled.

**Figure 6.**
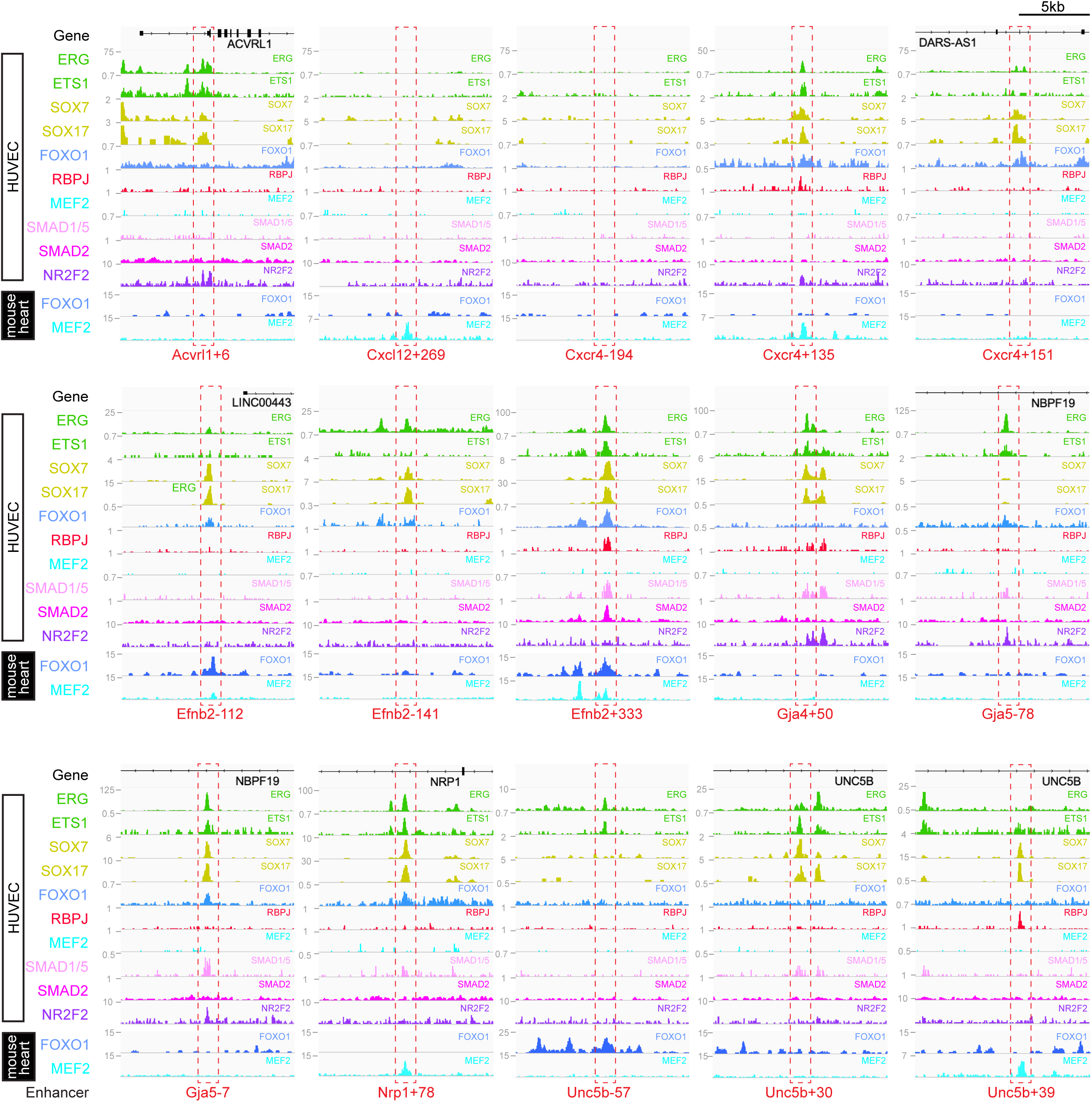
Genomic regions around each arterial enhancer identified in this paper (red dashed box) alongside tracks showing ChIP-seq/CUT&RUN signal for ERG ^32^, ETS1 ^33^, SOX7 and SOX17 (this paper), FOXO1 ^34^, RBPJ ^36^, MEF2C ^39^, SMAD1/5^37^, SMAD2 ^38^ and NR2F2 ^32^ in HUVECs, alongside FOXO1 ^32^ and MEF2A ^40^ in adult mouse hearts.

**Table 2.**
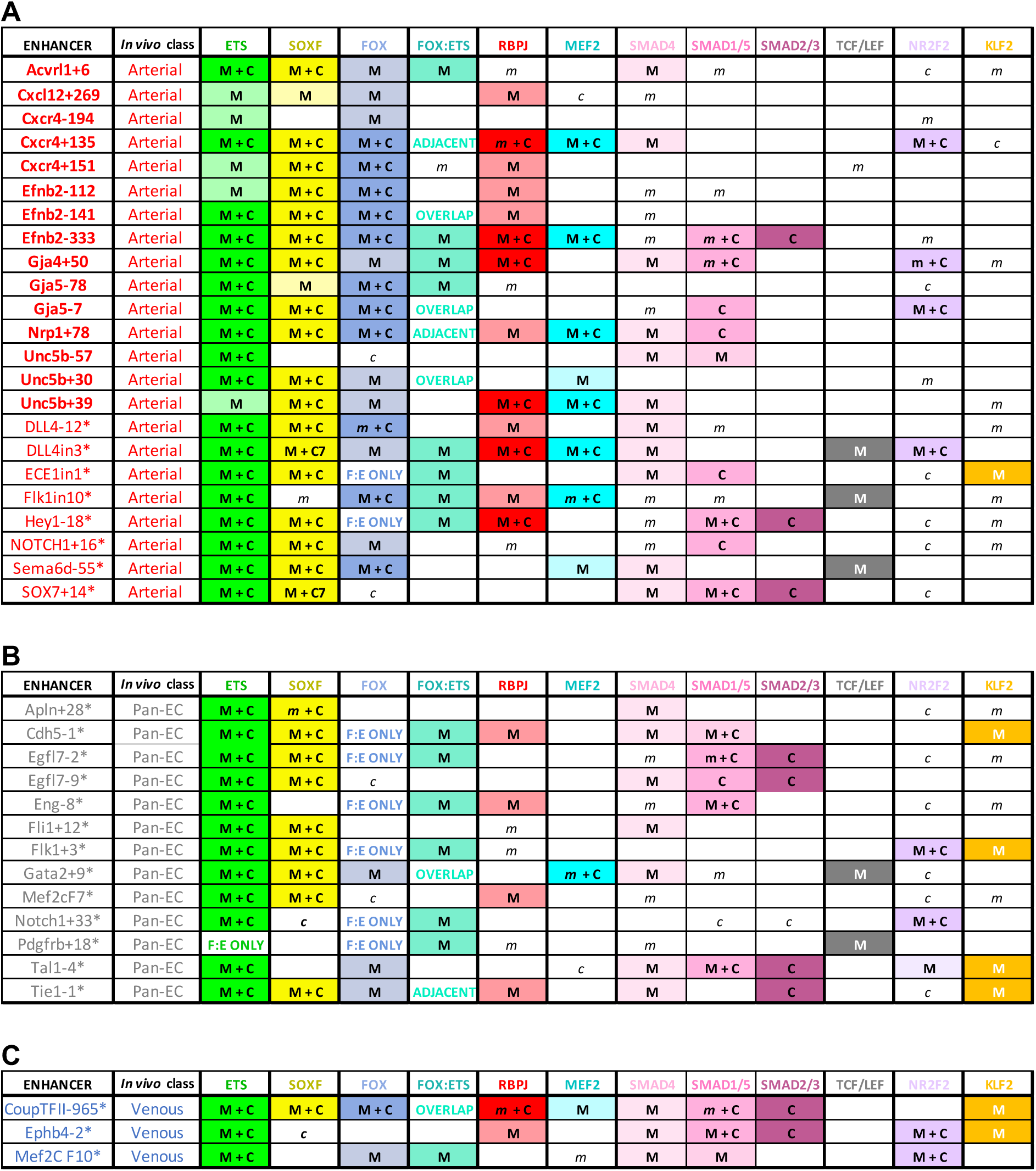
Summary of transcription factor motif and binding patterns at arterial (**A**), pan-EC (**B**) and venous (**C**) enhancers. All known (e.g. published) endothelial enhancers with adequately described expression patterns in transgenic mouse embryos were included in this analysis. Within these tables, an M indicates that an enhancer contains one of more motifs for the listed transcription factor that are conserved at the same depth as the surrounding enhancer sequence, while *m* indicates a motif that is conserved between the mouse and human enhancer sequence but is not conserved at the same depth as the surrounding enhancer sequence. See Figure S6 for motif information and S8 for annotated sequences. Within these tables, a C indicates a binding peak for the listed transcription factor (as assessed by ChIP-seq or similar) in an enhancer with a corresponding motif, while *c* indicates a binding peak for the listed transcription factor in an enhancer without a corresponding motif. See Figures 6, S7 and S9 for visualization of binding peaks. Enhancers listed in bold were identified in this paper, those with * are previously published (as detailed in ^1^), genome locations for each enhancer are provided in Table S3.

### ETS, SOXF and FOX binding is a common but not unique occurrence at arterial enhancers

Our motif analysis revealed near-ubiquitous binding sites for ETS, SOXF and FOX transcription factors at all arterial enhancers (Table 2A, Figure 5 and S8). For ETS, 23/23 arterial enhancers contained at least one conserved motif (all “deeply” conserved to the same depth as the surrounding enhancer, see S7). Confirming this motif identification, 18/23 of these enhancers also directly bound ERG and ETS1, two very common EC-associated ETS factors (Figure 6 and S8). For FOX, 21/23 arterial enhancers contained conserved FOX motifs (20 deeply conserved) with 13/23 directly binding FOXO1. For SOXF, 21/23 arterial enhancers contained conserved motifs (20 deeply conserved). Initial comparison of our enhancer cohort with publicly available information on SOX7 binding (from ^46^, using ChIP-seq in HUVECs over-expressing SOX7-mCherry) showed no overlap. However, only 6% of SOX7-mCherry peaks overlapped with EC-associated enhancers or TSS marks and only 4% correlated with ERG binding, an ETS family transcription factor strongly associated with EC gene transcription (Fig S10A, ERG and enhancer mark data from ^32^). Further, no SOX17 ChIP-seq in ECs has been published, despite SOX17 being most closely associated with arterial identity. We therefore performed a new assessment of SOX17 and SOX7 binding with antibodies against the endogenous proteins using CUT&RUN in HUVECs. In this new SOX17 analysis, 75% of binding peaks overlapped with EC enhancer/promoter marks^32^, 73% overlapped with ERG binding^32^, and HOMER analysis identified the SOX17 consensus motif as the most significantly enriched motif (Fig S10B-C). While our SOX7 binding analysis was slightly less specific (with 61% overlap with ERG-bound peaks and 62% overlap with enhancer/promoter marks), 86% of SOX17 peaks were bound by SOX7, and HOMER analysis again identified SOXF as a significantly enriched motif (Figure 9D-E). Assessment of our 23 arterial enhancer cohort using these new datasets found called SOXF peaks at 19/23 arterial enhancers (16 for SOX17 peaks, 14 for SOX7 peaks), in every case correlating with the presence of deeply conserved SOXF motifs (Table 2 and Figure 6 and S7).

Looked at in isolation, this analysis would strongly suggest that ETS, FOX and SOXF transcription factors work together to direct arterial-specific gene expression. However, although our enhancers are all restricted or strongly enriched in arterial ECs, this analysis cannot by itself distinguish between factors that specifically direct arterial expression, and those required for endothelial gene expression more generally. Consequently, to determine if these common ETS, SOXF and FOX binding patterns were unique to arterial enhancers, we expanded our analysis to 16 *in vivo*-validated endothelial enhancers that were not specifically active in arterial ECs (Table 2B-C, Figure 5 and S9). As assessed in transgenic mouse embryos at mid-late gestation, 13 of these enhancers drove relatively equal expression in arterial and venous ECs (pan-EC enhancers), while the activity of the other three was vein-enriched and artery-excluded (vein enhancers)^1^. Unsurprisingly, given their known role in general endothelial gene expression and identity, binding of ETS factors was ubiquitous at pan-EC enhancers and venous enhancers in addition to arterial enhancers. However, despite the association of SOXF factors with arterial identity, 10/13 pan-EC enhancers also directly bound SOX7/SOX17 (10/13 bound by SOX17, 9/13 by SOX7) and 8/13 contained deeply conserved SOXF motifs. The binding patterns at venous enhancers were harder to interpret due to a limited number of well-validated enhancers in this category. While the Ephb4-2 or Mef2cF10 vein enhancers contained no SOXF motifs and showed little evidence of robust SOXF binding, the CoupTFII-965 enhancer contained deeply conserved SOXF motifs and strongly bound both SOX7 and SOX17. Consequently, while the enhancers of some venous genes may lack sensitivity to SOXF factors, our data suggests that SOXF binding is not a unique feature of arterial-restricted gene expression. These observations align with previous studies showing roles for SOXF factors in vasculogenesis and angiogenesis^21,22,71^, and with the expression of SOXF factors throughout the vascular plexus during arterial-venous differentiation, as seen in the embryonic heart (Figure S11 and ^27,72^) and postnatal retina^73^. This also agrees with the severe vascular phenotypes seen after compound loss of SOXF factors, which include EC hyperplasia, loss of angiogenic markers and inhibited arteriovenous differentiation^22,73^.

Our findings were somewhat less clear for FOX transcription factors. 9/13 pan-EC enhancers contained some kind of FOX motif (compared to 21/23 arterial enhancers). However, the only FOX motif within 6 of these pan-EC enhancers was a composite part of a FOX:ETS motif (a vasculogenic-associated element whose FOX component is often fairly degenerative). This leaves only 3/13 pan-EC enhancers containing independent FOX motifs, compared to 19/23 arterial enhancers, suggesting that FOX binding may be enriched among our arterial enhancer cohort. While this aligns with previous observations linking FOXC1 and FOXC2 with arterial differentiation expression^74^, neither are highly expressed in developing coronary arteries nor commonly identified as arterial-enriched in single cell transcriptomics in developing mouse embryos (Figure S11 and ^4,75^). In addition to FOXC1/2, FOXO1 is also expressed widely throughout the endothelium and directly bound many of our arterial enhancers, but neither deletion or constitutive activation of FOXO1 in postnatal mouse retina prevented arterial differentiation^14^.

### MEF2 binding occurs at a subset of arterial enhancers

MEF2 factor binding was overrepresented in arterial enhancers compared to pan-EC enhancers, although it was seen at only a minority of enhancers. In total, 8/23 arterial enhancers contained conserved MEF2 motifs (7 deeply conserved), compared to only 1/13 pan-EC enhancers. This arterial-skewed pattern was repeated in assays of MEF2 factor binding, with direct MEF2 binding found at 7/23 arterial enhancers and only 2/13 pan-EC enhancers. Given the known role of MEF2 factors in angiogenic sprouting and the close link between angiogenesis and arterial differentiation ^67,76^, it is possible that MEF2 factors are regulating gene expression in response to angiogenic rather than arterial cues at these enhancers. This has been already shown for the Dll4in3 enhancer, where loss of MEF2 binding ablated enhancer activity in angiogenic ECs but not in mature arterial ECs^67^. Supporting this hypothesis, MEF2-bound enhancers were associated with genes expressed in early “pre-arterial” EC and/or involved in angiogenesis (*Cxcr4, Efnb2, Nrp1, Unc5b, Flk1* and *Dll4*)^4,20^. However, while MEF2 binding was not seen at the arterial enhancers least active in intersegmental sprouts (e.g. Efnb2-141, Gja5-7, Gja5-78, Unc5b+30), there was no obvious correlation between arterial enhancers driving the strongest expression in the intersegmental vessels at 2 dpf and MEF2 binding status.

### RBPJ binding indicates a role for Notch in transcription of arterial genes

RBPJ is the transcriptional effector of the Notch pathway, complexing with the Notch intracellular domain (NICD) and the co-activator MAML in order to directly bind DNA and activate transcription^77^. Deeply conserved RBPJ motifs were found in 12/23 arterial enhancers and 4/13 pan-EC enhancers, while direct RBPJ binding was confirmed at 6/23 arterial enhancers only (Table 2). RBPJ motifs are relatively short and share close similarity to the ETS motifs (consensus TGGGAA vs HGGAAR), potentially explaining the discrepancy between motif and direct binding. Direct RBPJ binding to arterial enhancers has previously only been reported for genes in the Notch pathway, supporting the hypothesis that Notch does not directly induce arterial differentiation through gene activation but instead by reducing their MYC-dependent metabolic and cell-cycle activities^13^. However, here we found good evidence for RBPJ binding at enhancers for *Cxcr4*, *Efnb2*, *Gja4* and *Unc5b,* suggesting that Notch may directly influence at least some arterial identity genes. In most cases, these genes also contained additional enhancers not directly bound by RBPJ, potentially providing an explanation for the maintenance of some arterial gene expression in the absence of RBPJ/Notch^13^. Additionally, previous studies on the Dll4in3 enhancer found that loss of RBPJ (and Notch signalling) did not affect enhancer or gene expression unless SOXF factors were also perturbed^25^. Cooperation between SOXF and Notch may also partly explain how the widely expressed SOXF factors enact arterial-specific gene activation, although SOX factors bound arterial enhancers more commonly than RBPJ. An alternative explanation may be that these results instead reflect the known involvement of RBPJ/Notch in angiogenesis. However, as with MEF2 binding, no obvious correlation between RBPJ binding and enhancer expression pattern in zebrafish can be seen, nor did MEF2 and RBPJ always target the same enhancers.

### No other arterial or venous-related transcription factors are commonly present or excluded at our arterial enhancers

Lastly, we investigated potential roles for SMADs (transcription factors downstream of TGFβ/BMP signalling), TCF7/TCF7L1/TCFL2/LEF1 (transcription factors downstream of canonical WNT signalling) and KLF2/4 (downstream of laminar shear stress). Although the binding motifs for these factors were not overrepresented in our arterial cohort as assessed by HOMER analysis, these pathways have all previously been implicated in arterial gene expression. We also looked for evidence of NR2F2/COUP-TFII binding, a vein and lymphatic-specific transcription factor previously implicated in both activation of venous genes and repression of arterial/Notch genes. This analysis found little evidence supporting a broad role for any of these pathways in arterial gene expression specificity. NR2F2 motifs were seen in 7/23 arterial enhancers (but only deeply conserved in 3/23) and 3/13 pan-EC enhancers. Largely uncorrelated ChIP-seq peaks were seen at 10/23 arterial enhancers, and at 10/13 pan-EC enhancers. Deeply conserved KLF motifs were only seen in 1/23 arterial and 4/13 pan-EC enhancers. Deeply conserved TCF/LEF motifs were only found in 3/23 arterial enhancers and 2/13 pan-EC enhancers. SMAD1/5-SMAD4 factors downstream of BMP signalling have been previously associated with the expression of venous genes including Nr2f2 and Ephb4^15,61^, and all three vein enhancers contained multiple motifs for SMAD factor binding (with 2/3 also directly binding SMAD1/5 and SMAD2 in HUVECS after BMP9/TGFβ stimulation) (Figure 5C). However, here we found that SMAD binding also occurred at arterial enhancers, with 12/23 containing motifs, 5/23 directly binding SMAD1/5 and 3/23 binding both SMAD1/5 and SMAD2. This agrees with the lack of venous-specificity reported for phosphorylated SMAD1/5, which led to the supposition that addition factors needed to work alongside SMAD1/5 to regulate vein specification^15^. An arterial role for SMADs is also not without precedent, particularly downstream of TGFβ (e.g.^78–80^. While Tie2:Cre-mediated excision of SMAD4 (effectively knocking out all canonical BMP/TGFβ signalling) did not obviously affect arterial differentiation at E9.5, an earlier or later role cannot be dismissed, as Tie2:Cre becomes active after vasculogenic-driven arterial differentiation occurs, and vein-related lethality occurs in these embryos by E10.5 ^15^, prior to most vein/capillary-to-arterial EC differentiation.

## DISCUSION

Recent years have brought a new appreciation of the vein/capillary origin of most arterial ECs, and an increasing interest in arterialization as a therapeutic aim of regenerative medicine. However, the transcriptional pathways driving arterial differentiation are still incompletely understood. Many factors have contributed to this, including a focus on the Notch signalling pathway and a lack of characterized arterial enhancers for most key arterial genes. The latter has resulted in regulatory pathways being linked to arterial gene expression through proposed binding at promoter regions, although these elements are often poorly characterized or unsupported by functional data (e.g. binding motifs located kilobases away from TSS at regions without promoter or enhancer marks, transcription factor binding not verified by available ChIP-seq datasets). As well as the potential for incorrect assumptions, a reliance on poorly defined enhancer/promoter regions prevents further research building on these initial observations, for example by looking for associated motifs to identify combinatorial, synergistic and antagonistic factors or to link with newly discovered pathways or transcription factors. In this paper, we sought to generate a useful and accessible cohort of arterial enhancers with which to study arterial transcriptional regulation more effectively. Alongside *Dll4*, *Notch1* and *Hey1* (all genes with previously described enhancers included in our analysis), the eight arterial genes focused on here represent the majority of genes used to define arterial identity in single cell transcriptomics in mice and humans (e.g. ^4,5,20,75^). Further, our choice of targets included genes with essential and well-studied roles in arterial differentiation (e.g. *Efnb2*), implicated in arteriovenous malformations in humans (e.g. *Acvrl1*), associated with processes important for regeneration (*Cxcr4* and *Cxcl12*^81^) or commonly used as arterial markers in animal models (e.g. *Gja5*). Thus, our hope was to identify a cohort of arterial enhancers likely to be directly targeted by arterial lineage specification and differentiation factors, that represent key stages of arterial development of interest to a wide range of researchers, and which we can easily link to previous observations on arterial development in animal models of gene depletion.

Analysis of single-cell transcriptomic data has indicated that arterial ECs can be further subdivided into two groups, reflecting maturity but also potentially slightly different developmental trajectories^4,20^. The genes studied here cover both subgroups, with *Acvrl1*, *Cxcl12*, *Gja5* and *Nrp1* primarily restricted to the mature arterial EC subgroup, while *Cxcr4*, *Efnb2*, *Gja4* and *Unc5b* were also expressed in the less mature/arterial plexus/pre-arterial EC subgroup^4,20^. Although we saw no obvious differences in transcription factor motif and binding between the two sets overall, the genes expressed in both immature and mature sub-groups tended to have multiple enhancers with differential expression patterns: there are three *Efnb2* enhancers, of which Efnb2-141 is restricted to the mature dorsal aorta while Efnb2-333 and Efnb2-121 enhancers are more widely active; there are three *Unc5b* enhancers, of which Unc5b+39 is restricted to the intersegmental arteries while Unc5b-57 and Unc5b+30 are expressed more widely. It is therefore possible that the upstream signals involved at different stages of arterial differentiation may, to some extent, target separate enhancers. This would align with our previous analysis of *Dll4*, a gene expressed in both arterial sub-groups and regulated by one enhancer active in mature arterial ECs only (Dll4-12) and one that is expressed in both angiogenic arterial precursors and mature arteries (Dll4in3). Although this paper has focused on transcriptional signatures of arterial vs non-arterial-specific enhancers, future research into the transcriptional differences seen between these differentially expressed arterial enhancers may therefore bring further insights into arterial transcription factors, and the manner in which upstream pathways combine to enact subtle but essential changes in gene expression.

Alongside a deficit of characterized enhancers, our understanding of vascular transcription is also affected by the considerable redundancy shown by many endothelial transcription factors. In particular, this can complicate analysis of gene disruption in animal models. A good example of this problem is the SOXF factors. SOX7, SOX17 and SOX18 not only show distinct yet overlapping expression patterns, but their ability to functionally compensate for each other can vary on different mouse backgrounds. For example, the phenotype in mice lacking SOX18 varies from essentially normal to complete loss of lymphatic ECs, with lethality depending on the mouse strain and associated variation in the ability of SOX7 and SOX17 to compensate^82^. SOX17 is the SOXF factor most strongly expressed in arterial ECs, and arterial defects occur after its deletion^23^. This has resulted in the hypothesis that SOX17 selectively activates arterial genes, but this is not well supported by the results here. An alternative explanation could be that SOXF factors are required for endothelial gene expression more generally, potentially as master regulators. This aligns with their robust and primarily endothelial-specific expression (particularly after mid-gestation)^1^), and by the widespread presence of SOXF motifs and binding at endothelial enhancers of all varieties. In this second model, the loss of SOX17 may affect the arterial compartment more severely simply because it comprises the majority of arterial SOXF (so the total amount of SOXF factors is more significantly depleted in arteries than elsewhere when SOX17 is deleted), with similar explanations for the consequences of SOX7 depletion on vasculogenesis^21^ and SOX18 depletion on lymphangiogenesis^82,83^. Alternatively, Stewen et al^61^ have recently shown that combined depletion of SOXF factors in cultured ECs significantly reduced *Efnb2* expression while increasing *Ephb4* expression, and instead argue for a more specific role for SOXF factors in arteriovenous differentiation related to elevated SOXF levels in arterial ECs ^61^. While this alone cannot explain the widespread binding of SOXF factors to pan-EC genes, SOXF factors are also crucial during vasculogenesis/early angiogenesis and may instead be more generally driving a general angiogenic/arterial gene expression program. Supporting this, neither the Ephb4-2 or Mef2cF7 vein enhancers had SOXF motifs, while the role of SOXF in CoupTFII-965 regulation may be simply related to its expression in lymphatic ECs, although the paucity of defined venous/capillary enhancers currently limits our ability to make conclusions here. However, endothelial SOXF factors are clearly strongly expressed widely in the endothelium at timepoints where a much more limited number of ECs become committed to an arterial fate (perfectly illustrated in the coronary vasculature), suggesting that additional transcriptional regulators must be involved alongside SOXF to enable this exquisitely specific pattern of gene activation. While RBPJ, MEF2 and FOX factors represent obvious potential partners, none would fully explain the specificity of all arterial enhancers and they all have wider roles in the vasculature.

Complicating analysis of our arterial enhancer cohort is the possibility that all arterial enhancers are not necessarily directly activated by the same regulator(s). Instead, a transcriptional cascade may be started by the activation of just one or two early genes, which then create a more permissive environment (e.g. high concentration or post-translation modification of transcription factors) for later arterial gene expression downstream of more widely expressed transcription factors. This would align with the observed elevated levels of SOXF expression as ECs switch to an arterial fate ^61^, and may suggest that the pathways upstream of SOXF expression play the most important role in arterial gene expression. However, a far more systematic analysis of all three SOXF loci, and the enhancers within, is required to test this hypothesis.

The relatively simple and cost-effective approach of in silico identification and zebrafish transgenesis of arterial enhancers used here had a success rate around 50%. This could doubtless be further refined (e.g. by including assessments of ERG binding, obtaining enhancer marks from *in vivo* arterial cells), and made more efficient by limiting verification to F0 transgenic fish or utilizing a higher throughput assay (e.g.^84^). However, a potentially more pressing issue is how to better understand the exact transcriptional regulators of these enhancers, a challenge shared with the gene regulatory field more widely. Transcription factors do not always bind DNA at their consensus motifs, with optimal syntax (order, orientation and spacing of motifs) often able to compensate for poor binding sites. Additionally, the presence of multiple motifs within a single enhancer, and the ability of many transcription factors to both directly and indirectly bind enhancers, means that enhancer sequence mutational analysis can be very complicated (e.g. ^25^, whilst restricting this analysis to a single timepoint (usually required to make such an approach practical), can be an issue where angiogenic and arterial programmes overlap. While assessments of direct binding by ChIP-seq or similar approaches can bypass a requirement to understand the exact motifs at an enhancer, neither cultured HUVECs nor iPSC cells recapitulate conditions in vivo, particularly regarding availability of ligands, exposure to shear stress and other environmental stimuli. Here, for example, the low expression of some of our arterial genes in culture HUVECs and HUAECs has probably affected verification of motifs with ChIP-seq data. Consequently, while our analysis here has provided clarity as to some transcription factors involved (and not involved) in arterial gene expression, none of our observations fully explain the shared ability of these short sequences of DNA to direct arterial patterns of expression even when removed from native chromatin context and endogenous promoters. Some of these answers can be expected to come from increasing identification of new or unappreciated transcription factors specifically expressed or specifically modified in either arterial or non-arterial ECs (e.g. MECOM), better appreciation of the consensus motifs and binding patterns of proteins already known to be involved (e.g. DACH1), and improved proteomic techniques. Additionally, new iPSC models of endothelial differentiation offer the opportunity to more easily study the consequences of transcription factor perturbation during angiogenesis and arterial differentiation, and artificial intelligence, improved bioinformatic pathways and machine learning all offer new avenues for research. It is anticipated that the cohort of in vivo-verified arterial enhancers characterized here will provide a vital platform for these future studies.

## Supporting information

All Supplemental Tables, Figures and Methods

## ACKNOWLEDGEMENTS

We thank Nadav Ahituv for providing the GW vectors. This work was supported by the BHF (FS/1735/32929 and FS/SBSRF/22/31037 to S.D.V. and S.N.; FS/IPBSRF/23/27085 to I.R.M.), the Oxford BHF Centre of Research Excellence (RE/18/3/34214) and by Ludwig Cancer Research Ltd.

## Notes

### Competing Interest Statement

The authors have declared no competing interest.

