## Supplementary material for "A catalogue of verified and characterized arterial enhancers for key arterial identity genes": All Supplemental Tables, Figures and Methods

**Table S1.** Enhancer marks around 31 known *in vivo*-characterized endothelial enhancers (all described in[1]). Human data in yellow, mouse data in green, red text indicates arterial enhancers. “H DNAseI” indicates open chromatin regions as defined by DNAseI hypersensitivity in HUVECs, HMVEC-dBl-neo and HMVEC-dBl-ad comparative to non-ECs and relative to surrounding region (UCSC genome browser[2]); “H histone” indicates relatively enriched binding of H3K27Ac and/or H3K4Me1 in HUVECs (UCSC genome browser[2]); “M artery ATAC” indicates regions of relatively open chromatin assessed by ATAC-seq in primary mouse adult aortic ECs (MAECs) (Engelbrecht et al[3]); “M retina ATAC” indicates regions of relatively open chromatin assessed by ATAC-seq in mouse postnatal day 6 (P6) retina ECs (MRECs) (Yanagida et al[4]); “M E11 p300” indicates regions relatively enriched for EP300 binding in Tie2Cre+ve cells from embryonic day (E)11.5 mouse embryos (Zhou et al[5]).

#### Table S1.

| Enhancer | hg19 coordinates | H DNAseI | H histone | Mm9 coordinates | M artery ATAC | M retina ATAC | M E11 p300 |
|---|---|---|---|---|---|---|---|
| Apln+28 | chrX:128,756,756-128,757,160 | YES | YES | chrX:45,359,306-45,359,632 | NO | YES | YES |
| Dab2-240 | chr5:39,755,997-39,756,596 | YES* | YES | chr15:6,009,719-6,010,138 | NO | NO | YES |
| Dll4in3 | chr15:41,222,881-41,223,570 | YES | YES | chr2:119,152,838-119,153,684 | YES | YES | YES |
| Dll4-12 | chr15:41,210,706-41,211,825 | YES | YES | chr2:119,140,274-119,141,353 | YES | YES | YES |
| Ece1in1 | chr1:21,606,038-21,607,057 | YES | YES | chr4:137,475,719-137,476,738 | YES | YES | YES |
| Egfl7-9 | chr9:139,540,750-139,541,299 | YES | YES | chr2:26,427,513-26,427,707 | YES | YES | YES |
| Egfl7-2 | chr9:139,550,292-139,550,891 | YES | YES | chr2:26,434,087-26,434,301 | YES | YES | YES |
| Emcn-22 | chr4:101,460,885-101,461,224 | NO | YES | chr3:136,984,547-136,984,951 | YES | NO | NO |
| Eng-8 | chr9:130,624,538-130,624,804 | YES | YES | chr2:32,493,606-32,493,823 | YES | YES | YES |
| Eng+9 | chr9:130,607,199-130,607,657 | YES | YES | chr2:32,511,282-32,511,641 | YES | YES | YES |
| Ephb4-2 | chr7:100,426,337-100,427,259 | YES | YES | chr5:137,789,910-137,790,581 | NO | NO | YES |
| Fli1+12 | chr11:128,575,436-128,575,782 | YES | YES | chr9:32,337,295-32,337,538 | YES | YES | YES |
| Flk1+3 | chr4:55,987,345-55,987,920 | YES | YES | chr5:76,370,627-76,371,056 | NO | NO | YES |
| Flk1in10 | chr4:55,972,978-55,973,903 | YES | NO | chr5:76,357,891-76,358,715 | NO | YES | YES |
| Flt4+26 | chr5:180,050,291-180,050,684 | NO | YES | chr11:49,445,777-49,446,175 | YES | YES | YES |
| Foxp1+138 | chr3:71,493,515-71,493,886 | YES | YES | chr6:99,338,958-99,339,515 | NO | YES | YES |
| Gata2+9 | chr3:128,201,971-128,202,273 | YES | YES | chr6:88,153,077-88,153,386 | YES | YES | YES |
| Hey1-18 | chr8:80,695,610-80,697,109 | YES | YES | chr3:8,685,099-8,685,821 | YES | YES | YES |
| Hlx-3 | chr1:221,049,978-221,050,354 | YES | YES | chr1:186,558,918-186,559,303 | YES | YES | YES |
| Mef2F10 | chr5:88,110,980-88,111,253 | YES | YES | chr13:83,721,761-83,722,057 | NO | YES | YES |
| Mef2F7 | chr5:88,123,031-88,123,357 | YES | YES | chr13:83,711,180-83,711,509 | YES | YES | YES |
| Notch1+16 | chr9:139,424,543-139,424,953 | YES | YES | chr2:26,346,100-26,346,671 | YES | YES | YES |
| Notch1+33 | chr9:139,406,356-139,406,655 | YES | YES | chr2:26,330,559-26,330,785 | YES | YES | YES |
| CoupTFII-965 | chr15:95,908,708-95,909,240 | YES | YES | chr7:78,456,407-78,456,767 | YES | YES | YES |
| Nrp1+28 | chr10:33,590,960-33,591,499 | NO | YES | chr8:130,911,132-130,911,551 | YES | YES | YES |
| Nrp2+26 | chr2:206,573,202-206,573,523 | YES | YES | chr1:62,776,231-62,776,553 | YES | YES | YES |
| Pdgfrb+18 | chr5:149,516,883-149,517,356 | NO | YES | chr18:61,219,244-61,219,566 | NO | NO | NO |
| Epcr-5 | chr20:33,754,176-33,754,585 | YES | YES | chr2:155,568,588-155,569,127 | YES | NO | YES |
| Sema6d-55 | chr15:47,958,023-47,958,764 | YES | YES | chr2:124,380,522-124,381,285 | YES | YES | YES |
| Sox7+14 | chr8:10,573,085-10,574,291 | YES | YES | chr14:64,576,271-64,577,533 | YES | YES | YES |
| Tal+19 | chr1:47,677,539-47,677,958 | YES | YES | chr4:114,748,131-114,748,530 | YES | YES | YES |
| Tal1-4 | chr1:47,701,050-47,701,347 | NO | YES | chr4:114,725,243-114,725,552 | YES | YES | YES |

**Table S2.** Enhancer marks in different human and mouse ECs at potential enhancer regions within the loci of eight arterial genes. “Selected” indicates that the region meets our threshold as a putative enhancer, “exception” indicates region did not meet our threshold but was included in transgenic analysis as a control. Numbers indicate approximate distance from the TSS of the named arterial gene. * indicates that enhancer mark was widely seen beyond endothelial cells. Grey italic text refers to regions previously implicated in enhancer activity, with the /enhancer name ascribed in the original reference. Cxcr4-117/CXCR4-125 is from [6]; Cxcr4-1, Nrp1-1/NRP1A and Nrp1+76/NRP1B are from [7]; Efnb2+17/EFNB2A and Efnb2+25/EFNB2B are from [8]; and Efnb2+4/EFNB2R1 and Efnb2+28/EFNB2R4 are from [9].

### Table S2

| Enhancer | H DNAseI | H histone | M artery ATAC | M retina ATAC | M E11 p300 | Selected | Exception |
|---|---|---|---|---|---|---|---|
| **Acvrl1-5** | YES | YES | NO | NO | YES | **SELECTED** | |
| **Acvrl1-1/p** | YES | YES | NO | YES | YES | **SELECTED** | |
| **Acvrl1+6** | YES | YES | NO | YES | YES | **SELECTED** | |
| *Acvrl1+16* | NO | NO | YES | YES | YES | NO | MOUSE ONLY |
| **Acvrl1+19** | YES | YES | YES | NO | YES | **SELECTED** | |
| *Cxcl12-184* | YES | YES | NO | NO | NO | NO | HUMAN ONLY |
| **Cxcl12-2** | NO | NO | YES | YES | NO | **SELECTED** | |
| *Cxcl12+239* | NO | NO | NO | YES | NO | NO | 1 MARK ONLY |
| *Cxcl12+265* | NO | NO | NO | YES | NO | NO | 1 MARK ONLY |
| **Cxcl12+269** | NO | NO | YES | YES | NO | **SELECTED** | |
| **Cxcl12+298** | YES | YES | NO | YES | YES | **SELECTED** | |
| **Cxcl12+376** | YES | NO | YES | NO | NO | **SELECTED** | |
| **Cxcl12+383** | NO | YES | YES | YES | YES | **SELECTED** | |
| *Cxcl12+439* | YES | YES | NO | NO | NO | NO | HUMAN ONLY |
| **Cxcl12+445** | YES* | YES | YES | YES | NO | **SELECTED** | |
| **Cxcr4-232** | YES | NO | NO | YES | NO | **SELECTED** | |
| **Cxcr4-194** | NO | NO | YES | NO | YES | **SELECTED** | |
| **Cxcr4-130** | NO | NO | NO | YES | YES | **SELECTED** | |
| *Cxcr4-117/CXCR4-125* | YES | YES | NO | NO | NO | NO | LITERATURE |
| **Cxcr4-113** | YES | YES | YES | YES | YES | **SELECTED** | |
| **Cxcr4-109** | YES | YES | YES | YES | YES | **SELECTED** | |
| **Cxcr4+1** | NO | NO | YES | YES | NO | **SELECTED** | |
| **Cxcr4+135** | YES | YES | YES | YES | YES | **SELECTED** | |
| **Cxcr4+151** | YES | YES | NO | YES | YES | **SELECTED** | |
| **Efnb2-333** | YES | YES | YES | YES | YES | **SELECTED** | |
| **Efnb2-141** | YES | YES | YES | YES | YES | **SELECTED** | |
| **Efnb2-112** | YES | YES | NO | YES | YES | **SELECTED** | |
| **Efnb2+3** | YES* | YES | YES | NO | NO | **SELECTED** | |
| *Efnb2+4/EFNB2 R1* | YES | NO | NO | NO | NO | NO | |
| *Efnb2+17/EFNB2 A* | NO | YES | NO | NO | NO | NO | |
| *Efnb2+25/EFNB2 B* | NO | NO | NO | NO | NO | NO | |
| *Efnb2+28/EFNB2 R4* | NO | NO | NO | NO | NO | NO | |
| **Efnb2+37** | YES | YES | YES | YES | YES | **SELECTED** | |
| **Efnb2+172** | YES | YES | YES | NO | YES | **SELECTED** | |
| **Efnb2+209** | YES* | YES | NO | YES | YES | **SELECTED** | |
| **Gja+24** | NO | YES | YES | NO | YES | **SELECTED** | |
| **Gja4+50** | YES | NO | YES | YES | YES | **SELECTED** | |
| *Gja4+57* | NO | NO | NO | NO | YES | NO | 1 MARK ONLY |
| **Gja5-93** | NO | NO | YES | YES | YES | **SELECTED** | |
| **Gja5-78** | NO | YES | YES | YES | YES | **SELECTED** | |
| **Gja5-28** | YES | NO | YES | NO | NO | **SELECTED** | |
| *Gja5-21* | YES | YES | NO | NO | NO | NO | HUMAN ONLY |
| **Gja5-7** | YES | YES | YES | YES | NO | **SELECTED** | |
| *Nrp1-1/NRP1 A* | YES* | NO | NO | NO | NO | NO | |
| **Nrp1+28** | NO | YES | NO | YES | YES | **SELECTED** | |
| *Nrp1+76/ NRP1 B* | YES* | YES | NO | NO | NO | NO | LITERATURE |
| **Nrp1+78** | YES | YES | NO | YES | YES | **SELECTED** | |
| **Nrp1+91** | YES | YES | YES | YES | YES | **SELECTED** | |
| **Nrp1+129** | YES | YES | NO | YES | YES | **SELECTED** | |
| **Unc5b-57** | NO | YES | NO | NO | YES | **SELECTED** | |
| *Unc5b+14* | NO | NO | YES | YES | NO | NO | MOUSE ONLY |
| **Unc5b+23** | YES* | YES | YES | YES | YES | **SELECTED** | |
| **Unc5b+30** | YES | YES | NO | YES | YES | **SELECTED** | |
| **Unc5b+39** | YES | YES | NO | YES | YES | **SELECTED** | |
| *Unc5b+43* | YES | YES | NO | NO | NO | NO | HUMAN ONLY |

**Table S3.** Genome locations for all enhancer (and human orthologues) investigated in this paper. Values given for mm9 represent the core enhancer regions analysed for motif sequences and protein binding.

### Table S3

**ARTERIAL ENHANCERS**

| name | mm10 of CORE | mm9 | hg38 | hg19 |
|---|---|---|---|---|
| **Acvrl1+6A** | chr15:101,134,090-101,134,328 | chr15:100,964,449-100,964,836 | chr12:51,911,761-51,912,189 | chr12:52,305,545-52,305,973 |
| **Cxcl12+269** | chr6:117,437,703-117,437,974 | chr6:117,387,534-117,388,459 | chr10:44,043,453-44,044,439 | chr10:44,538,901-44,539,887 |
| **Cxcr4-194** | chr1:128,785,634-128,786,023 | chr1:130,681,776-130,682,867 | chr2:136,325,218-136,326,027 | chr2:137,082,788-137,083,597 |
| **Cxcr4+135** | chr1:128,457,037-128,457,301 | chr1:130,353,525-130,353,952 | chr2:136,014,017-136,014,436 | chr2:136,771,587-136,772,006 |
| **Cxcr4+151** | chr1:128,440,683-128,440,929 | chr1:130,337,167-130,337,580 | chr2:136,000,071-136,000,490 | chr2:136,757,641-136,758,060 |
| **Efnb2-112** | chr8:8,772,250-8,772,661 | chr8:8,772,171-8,772,912 | chr13:106,651,110-106,652,084 | chr13:107,303,458-107,304,432 |
| **Efnb2-141** | chr8:8,801,731-8,802,062 | chr8:8,801,433-8,802,174 | chr13:106,680,416-106,681,557 | chr13:107,332,764-107,333,905 |
| **Efnb2-333** | chr8:8,994,388-8,994,688 | chr8:8,994,342-8,994,890 | chr13:106,919,439-106,920,184 | chr13:107,571,787-107,572,532 |
| **Gja4+50** | chr4:127,263,807-127,264,170 | chr4:126,940,851-126,941,567 | chr1:34,842,968-34,843,574 | chr1:35,308,569-35,309,175 |
| **Gja5-78** | chr3:96,953,907-96,954,116 | chr3:96,757,582-96,758,245 | chr1:147,877,925-147,878,475 | chr1:147,349,718-147,350,707 |
| **Gja5-7** | chr3:97,025,457-97,025,703 | chr3:96,829,228-96,829,714 | chr1:147,781,009-147,781,500 | chr1:147,253,120-147,253,611 |
| **Nrp1+78** | chr8:128,437,372-128,437,681 | chr8:130,961,192-130,961,715 | chr10:33,249,630-33,250,192 | chr10:33,538,558-33,539,120 |
| **Unc5b-57** | chr10:60,888,540-60,888,832 | chr10:60,351,050-60,351,950 | chr10:71,132,227-71,133,212 | chr10:72,891,984-72,892,969 |
| **Unc5b+30** | chr10:60,807,701-60,807,963 | chr10:60,263,361-60,264,141 | chr10:71,254,099-71,255,108 | chr10:73,013,856-73,014,865 |
| **Unc5b+39** | chr10:60,800,879-60,801,144 | chr10:60,255,453-60,256,125 | chr10:71,263,111-71,264,120 | chr10:73,022,868-73,023,877 |
| DLL4-12* | chr2:119,314,902-119,315,163 | chr2:119,140,274-119,141,353 | chr15:40,918,508-40,919,627 | chr15:41,210,706-41,211,825 |
| DLL4in3* | chr2:119,327,292-119,327,550 | chr2:119,152,838-119,153,684 | chr15:40,930,683-40,931,372 | chr15:41,222,881-41,223,570 |
| ECE1* | chr4:137,920,171-137,920,428 | chr4:137,475,719-137,476,738 | chr1:21,279,545-21,280,564 | chr1:21,606,038-21,607,057 |
| Flk1in10* | chr5:75,962,034-75,962,270 | chr5:76,357,891-76,358,715 | chr4:55,106,811-55,107,736 | chr4:55,972,978-55,973,903 |
| Hey1-18* | chr3:8,685,313-8,685,498 | chr3:8,685,099-8,685,821 | chr8:79,783,375-79,784,874 | chr8:80,695,610-80,697,109 |
| NOTCH1+16* | chr2:26,490,736-26,491,004 | chr2:26,346,100-26,346,671 | chr9:136,530,091-136,530,501 | chr9:139,424,543-139,424,953 |
| Sema6d-55* | chr2:124,555,118-124,555,549 | chr2:124,380,522-124,381,285 | chr15:47,665,826-47,666,567 | chr15:47,958,023-47,958,764 |
| SOX7+14* | chr14:63,957,836-63,958,080 | chr14:64,576,271-64,577,533 | chr8:10,715,575-10,716,781 | chr8:10,573,085-10,574,291 |

**PAN-EC ENHANCERS**

| name | mm10 of CORE | mm9 | hg38 | hg19 |
|---|---|---|---|---|
| Apln+28 | chrX:48,006,129-48,006,455 | chrX:45,359,306-45,359,632 | chrX:129,622,779-129,623,183 | chrX:128,756,756-128,757,160 |
| Cdh5-1 | chr8:104,101,460-104,101,691 | chr8:106,625,360-106,625,591 | chr16:66,366,342-66,366,812 | chr16:66,400,303-66,400,656 |
| Egfl7-2 | chr2:26,578,567-26,578,781 | chr2:26,434,087-26,434,301 | chr9:136,655,840-136,656,439 | chr9:139,550,292-139,550,891 |
| Egfl7-9 | chr2:26,571,993-26,572,187 | chr2:26,427,513-26,427,707 | chr9:136,646,298-136,646,847 | chr9:139,540,750-139,541,299 |
| Eng-8 | chr2:32,638,086-32,638,303 | chr2:32,493,606-32,493,823 | chr9:127,862,259-127,862,525 | chr9:130,624,538-130,624,804 |
| Fli1+12 | chr9:32,529,710-32,529,953 | chr9:32,337,295-32,337,538 | chr11:128,705,389-128,705,988 | chr11:128,575,436-128,575,782 |
| Flk1+3 | chr5:75,974,602-75,975,031 | chr5:76,370,627-76,371,056 | chr4:55,121,178-55,121,753 | chr4:55,987,345-55,987,920 |
| Gata2+9 | chr6:88,203,083-88,203,392 | chr6:88,153,077-88,153,386 | chr3:128,483,128-128,483,430 | chr3:128,201,971-128,202,273 |
| Mef2cF7 | chr13:83,711,228-83,711,450 | chr13:83,572,070-83,572,292 | chr5:88,827,268-88,827,485 | chr5:88,123,031-88,123,357 |
| Notch1+33 | chr2:26,475,039-26,475,265 | chr2:26,330,559-26,330,785 | chr9:136,511,904-136,512,203 | chr9:139,406,356-139,406,655 |
| Pdgfrb+18 | chr18:61,059,590-61,059,912 | chr18:61,219,244-61,219,566 | chr5:150,137,320-150,137,793 | chr5:149,516,883-149,517,356 |
| Tal1-4 | chr4:115,052,638-115,052,947 | chr4:114,725,243-114,725,552 | chr1:47,235,413-47,235,675 | chr1:47,701,050-47,701,347 |
| Tie1-1 | chr4:118,489,769-118,490,150 | chr4:118,162,374-118,162,755 | chr1:43,300,547-43,301,064 | chr1:43,766,218-43,766,735 |

**VEIN ENHANCERS**

| name | mm10 of CORE | mm9 | hg38 | hg19 |
|---|---|---|---|---|
| CoupTFII-965 | chr7:71,311,521-71,311,881 | chr7:78,456,407-78,456,767 | chr15:95,365,479-95,366,011 | chr15:95,908,708-95,909,240 |
| Ephb4-2 | chr5:137,348,682-137,349,353 | chr5:137,789,910-137,790,581 | chr7:100,828,715-100,829,637 | chr7:100,426,337-100,427,259 |
| Mef2cF10 | chr13:83,582,603-83,582,899 | chr13:83,721,761-83,722,057 | chr5:88,815,163-88,815,436 | chr5:88,110,980-88,111,253 |

**Figure S1.** Enhancer marks around eight target arterial genes. Red and orange solid boxes denote regions identified as putative enhancers (red indicates activity in transgenic models, orange indicates no activity in transgenic models, see Table 1 and Figure 2), numbers represent approximate distance from TSS. Orange dashed boxes indicates regions below the putative enhancer threshold but included in transgenic assays as controls, grey boxes indicate regions below the putative enhancer threshold and not tested. * indicates that enhancer marks were not specific for ECs but rather found in many cell types. Enhancer marks in mouse cells: dark red "ATAC adult artery EC" denotes open chromatin as assessed by ATAC-seq in primary mouse adult aortic ECs from Engelbrecht et al[3]; bright red "ATAC P6 EC" denotes open chromatin as assessed by ATAC-seq in mouse postnatal day 6 (P6) retina ECs from Yanagida et al[4]; orange "EP300 E11 EC" denotes enriched EP300 binding in Tie2Cre+ve cells in embryonic day (E)11.5 mouse embryos from Zhou et al[5]. Enhancer marks in human cells: light blue peaks denote enriched H3K27Ac and H3K4Me1 in human umbilical vein ECs (HUVECs), data from the UCSC genome browser[2]; grey heat map denotes open chromatin regions assessed by DNAseI hypersensitivity in HUVECs (upper line) and dermal-derived neonatal and adult blood microvascular ECs (HMVEC-dBl-neo/ad, middle and bottom line) from the UCSC genome browser.

### Figure S1

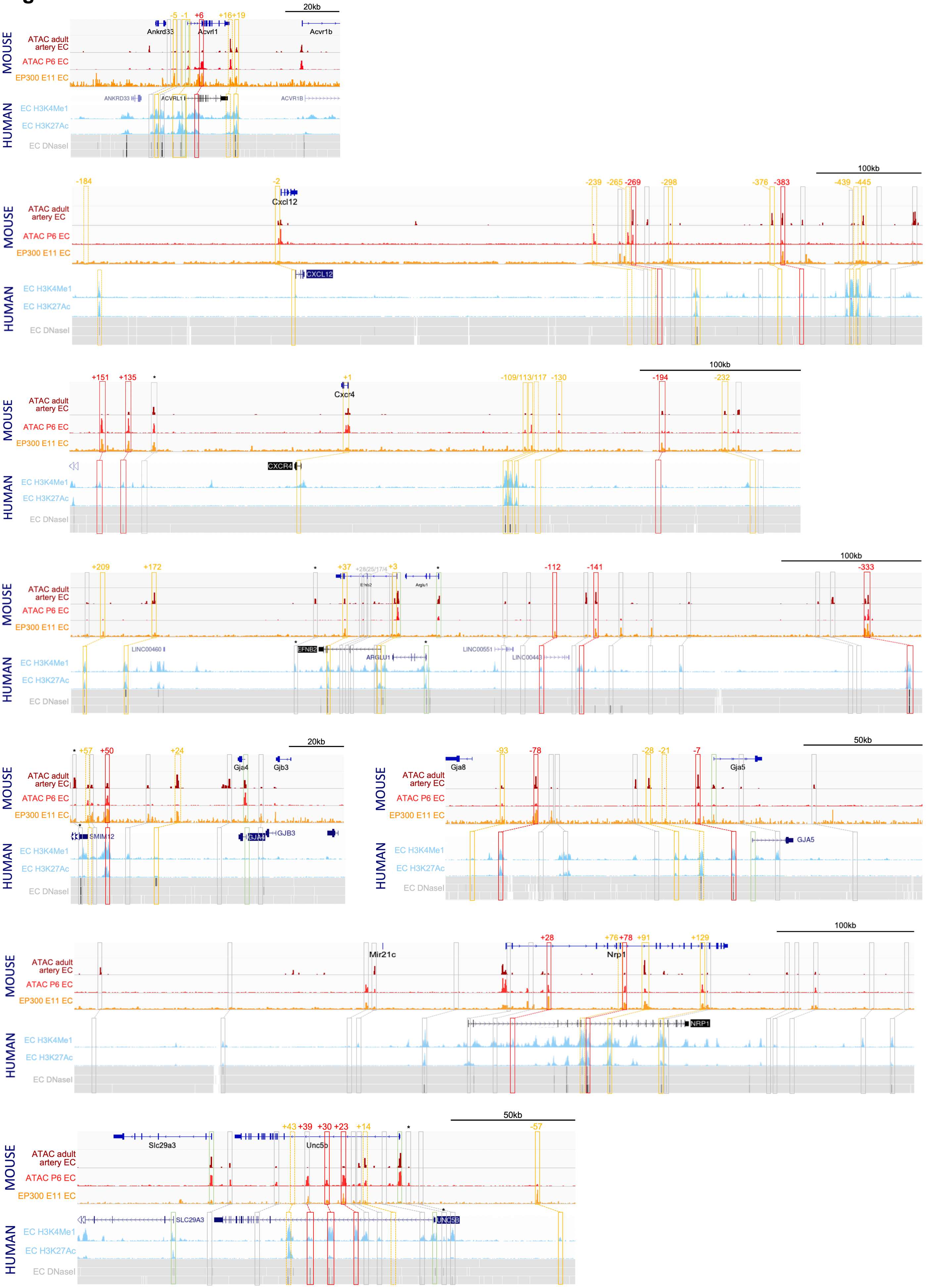

MOUSE
HUMAN
ATAC adult artery EC
ATAC P6 EC
EP300 E11 EC
EC H3K4Me1
EC H3K27Ac
EC DNaseI
20kb
Ankrd33
Acvrl1
Acvr1b
ANKRD33
ACVRL1
ACVR1B
-5 -1 +6 +16+19
100kb
Cxcl12
CXCL12
-184 -2 -239 -265 -269 -298 -376 -383 -439 -445
Cxcr4
CXCR4
+151 +135 +1 -109/113/117 -130 -194 -232
Efnb2
Arglu1
EFNB2
ARGLU1
LINC00460
LINC00551
LINC00443
+209 +172 +37 +28/25/17/4 +3 -112 -141 -333
Gja4
Gjb3
SMIM12
GJA4
GJB3
+57 +50 +24
50kb
Gja8
Gja5
GJA5
-93 -78 -28 -21 -7
Mir21c
Nrp1
NRP1
+28 +76 +78 +91 +129
Slc29a3
Unc5b
SLC29A3
UNC5B
+43 +39 +30 +23 +14 -57

**Figure S2**. The Dll4-12 enhancer directs arterial expression of reporter genes in the vasculature of transgenic zebrafish and mice. This enhancer was first reported in [10] but not tested in zebrafish.

**A**. The mouse Dll4-12:GFP transgene directs arterial endothelial cell expression in mosaic F0 (upper) and stable F1 (lower) transgenic zebrafish at two days post fertilization (dpf). Grey dashed box specifies region of zoom, a indicates dorsal aorta, v indicates cardinal vein, * indicates intersegmental vessels. Non-EC GFP expression was also seen in neural tube n.

**B**. The mouse Dll4-12:LacZ transgene directs arterial expression in a stable transgenic line. Representative whole-mount embryos from the Dll4-12:*lacZ* transgenic line show reporter gene expression (X-gal staining, blue) in the vasculature from embryonic day 9.5 (E9.5) to E15.5.

#### Figure S2

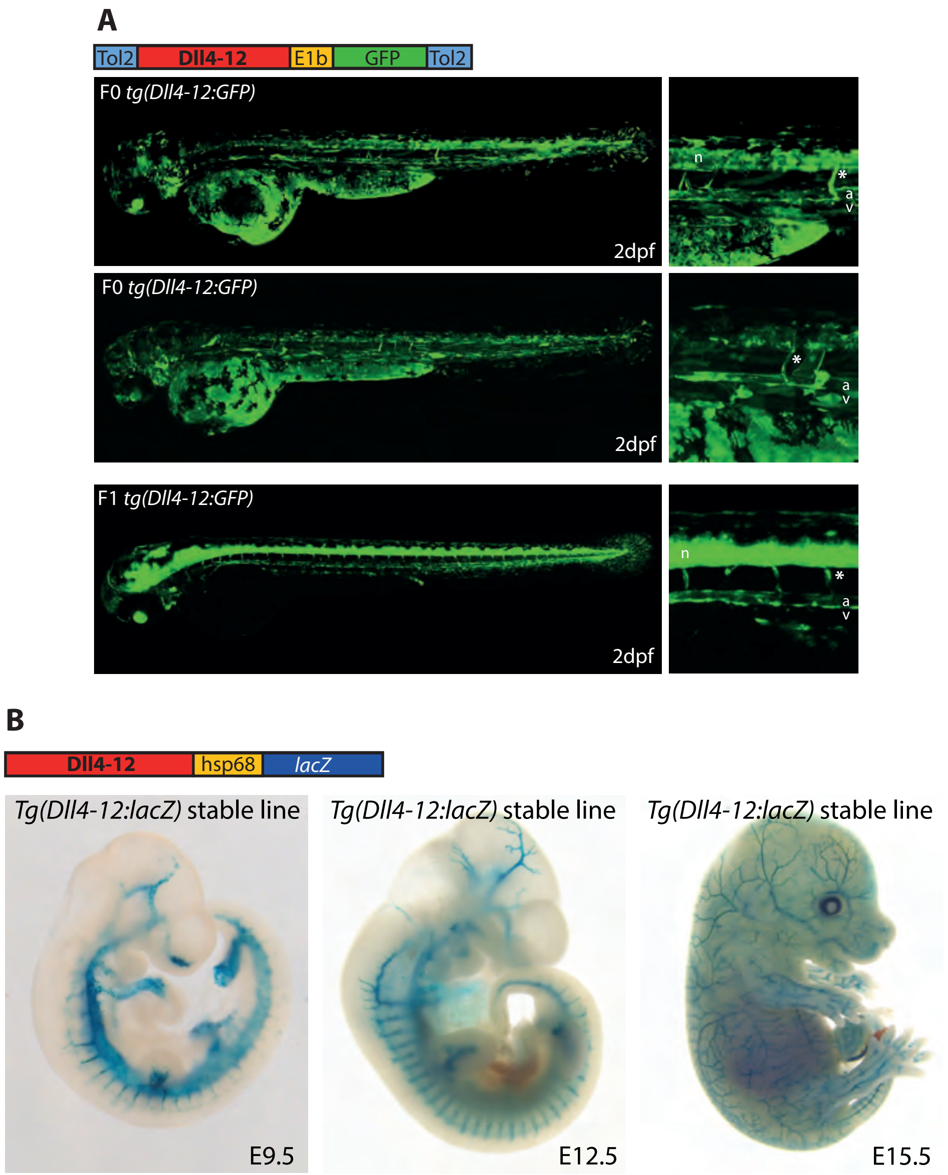
A
Tol2
Dll4-12
E1b
GFP
Tol2
F0 tg(Dll4-12:GFP)
2dpf
n
*
a
v
F0 tg(Dll4-12:GFP)
2dpf
*
a
v
F1 tg(Dll4-12:GFP)
2dpf
n
*
a
v
B
Dll4-12
hsp68
lacZ
Tg(Dll4-12:lacZ) stable line
E9.5
Tg(Dll4-12:lacZ) stable line
E12.5
Tg(Dll4-12:lacZ) stable line
E15.5

**Figure S3.** The Cxcr4-117:GFP transgene directs very limited reporter gene expression in transgenic zebrafish. The six transgenic zebrafish shown exhibited the greatest level of GFP expression seen in all injected zebrafish. Grey dashed box indicates region of zoom.

### Figure S3

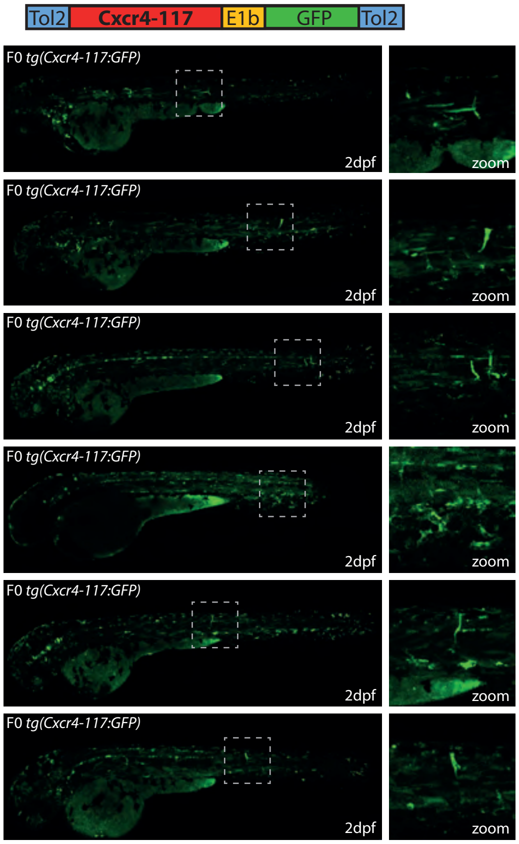

Tol2
Cxcr4-117
E1b
GFP
Tol2
F0 tg(Cxcr4-117:GFP)
2dpf
zoom
F0 tg(Cxcr4-117:GFP)
2dpf
zoom
F0 tg(Cxcr4-117:GFP)
2dpf
zoom
F0 tg(Cxcr4-117:GFP)
2dpf
zoom
F0 tg(Cxcr4-117:GFP)
2dpf
zoom
F0 tg(Cxcr4-117:GFP)
2dpf
zoom

**Figure S4.** All transgenic embryos expressing the Efnb2-37:lacZ transgene at E14.5 Grey dashed boxes indicate region in zoom. No expression is seen in vessels.

### Figure S4

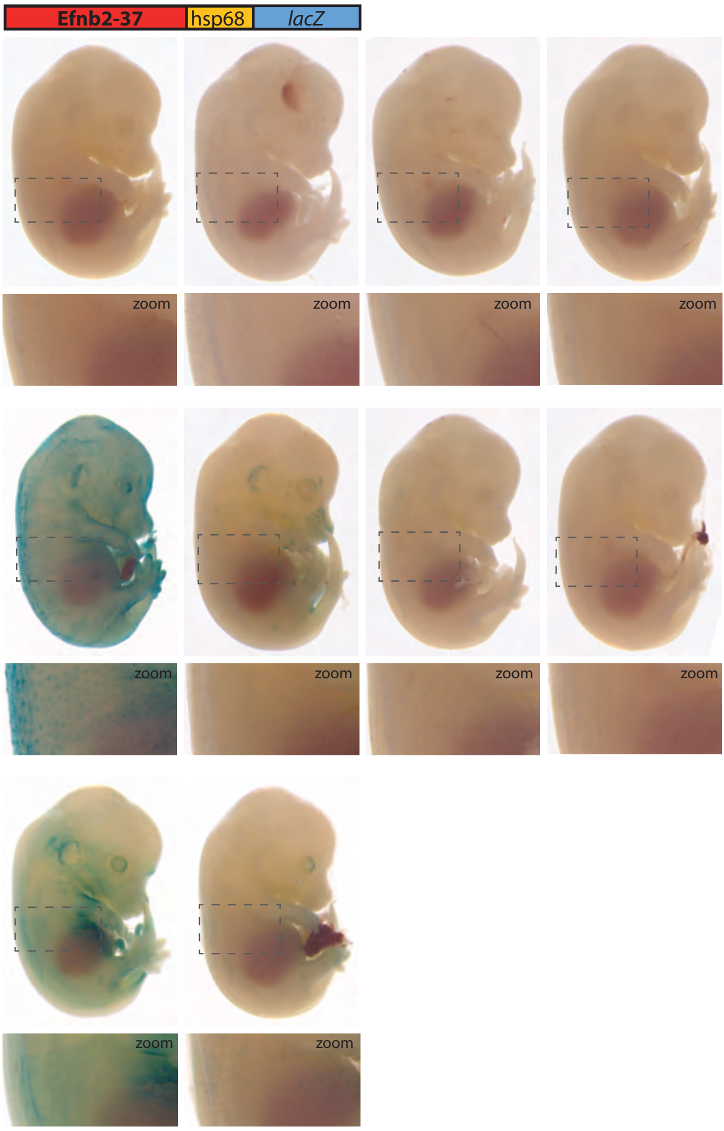

**Figure S5.** Homer Known Motif Enrichment Results on core enhancer sequences from all 23 arterial enhancers listed in Table 1. Exact sequences include in this analysis are listed in Figure S8, and the coordinates in mm10 recorded in Table S3.

### Figure S5

| Rank | Motif | Name | P-value | log P-pvalue |
|---|---|---|---|---|
| 1 | 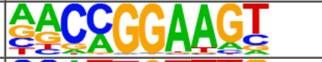 | ETV1(ETS)/GIST48-ETV1-ChIP-Seq(GSE22441)/Homer | 1e-10 | -2.344e+01 |
| 2 | 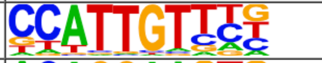 | Sox17(HMG)/Endoderm-Sox17-ChIP-Seq(GSE61475)/Homer | 1e-9 | -2.156e+01 |
| 3 | 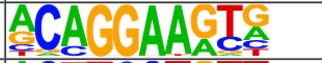 | ERG(ETS)/VCaP-ERG-ChIP-Seq(GSE14097)/Homer | 1e-8 | -2.063e+01 |
| 4 | 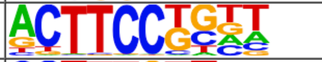 | Elf4(ETS)/BMDM-Elf4-ChIP-Seq(GSE88699)/Homer | 1e-8 | -2.057e+01 |
| 5 | 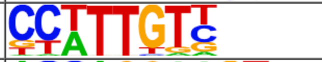 | Sox3(HMG)/NPC-Sox3-ChIP-Seq(GSE33059)/Homer | 1e-8 | -2.009e+01 |
| 6 | 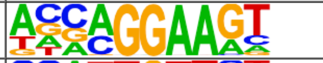 | EHF(ETS)/LoVo-EHF-ChIP-Seq(GSE49402)/Homer | 1e-8 | -1.986e+01 |
| 7 | 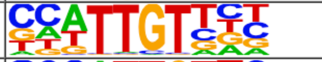 | Sox6(HMG)/Myotubes-Sox6-ChIP-Seq(GSE32627)/Homer | 1e-8 | -1.972e+01 |
| 8 | 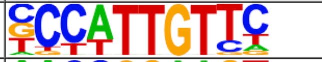 | Sox2(HMG)/mES-Sox2-ChIP-Seq(GSE11431)/Homer | 1e-7 | -1.768e+01 |
| 9 | 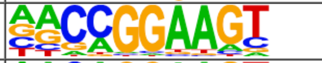 | GABPA(ETS)/Jurkat-GABPa-ChIP-Seq(GSE17954)/Homer | 1e-7 | -1.713e+01 |
| 10 | 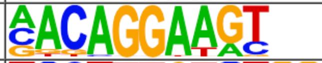 | Ets1-distal(ETS)/CD4+-PolII-ChIP-Seq(Barski_et_al.)/Homer | 1e-7 | -1.712e+01 |
| 11 | 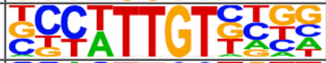 | Sox21(HMG)/ESC-SOX21-ChIP-Seq(GSE110505)/Homer | 1e-7 | -1.694e+01 |
| 12 | 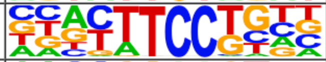 | Etv2(ETS)/ES-ER71-ChIP-Seq(GSE59402)/Homer | 1e-7 | -1.681e+01 |
| 13 | 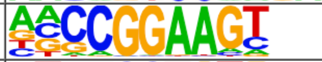 | ELF1(ETS)/Jurkat-ELF1-ChIP-Seq(SRA014231)/Homer | 1e-7 | -1.672e+01 |
| 14 | 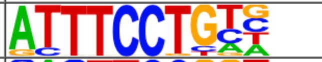 | EWS:ERG-fusion(ETS)/CADO_ES1-EWS:ERG-ChIP-Seq(SRA014231)/Homer | 1e-7 | -1.658e+01 |
| 15 | 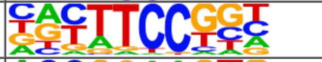 | Fli1(ETS)/CD8-FLI-ChIP-Seq(GSE20898)/Homer | 1e-7 | -1.656e+01 |
| 16 | 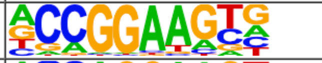 | ETV4(ETS)/HepG2-ETV4-ChIP-Seq(ENCODE)/Homer | 1e-7 | -1.621e+01 |
| 17 | 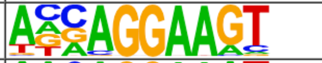 | ELF3(ETS)/PDAC-ELF3-ChIP-Seq(GSE64557)/Homer | 1e-6 | -1.570e+01 |
| 18 | 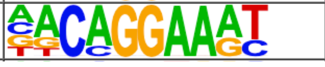 | EWS:FLI1-fusion(ETS)/SK_N_MC-EWS:FLI1-ChIP-Seq(SRA014231)/Homer | 1e-6 | -1.558e+01 |

| Rank | Motif | Name | P-value | log P-pvalue |
|---|---|---|---|---|
| 19 | 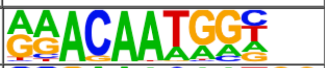 | Sox15(HMG)/CPA-Sox15-ChIP-Seq(GSE62909)/Homer | 1e-6 | -1.549e+01 |
| 20 | 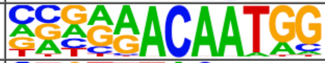 | Sox7(HMG)/ESC-Sox7-ChIP-Seq(GSE133899)/Homer | 1e-6 | -1.410e+01 |
| 21 | 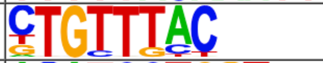 | Foxo1(Forkhead)/RAW-Foxo1-ChIP-Seq(Fan_et_al.)/Homer | 1e-6 | -1.408e+01 |
| 22 | 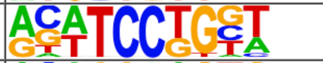 | SPDEF(ETS)/VCaP-SPDEF-ChIP-Seq(SRA014231)/Homer | 1e-6 | -1.406e+01 |
| 23 | 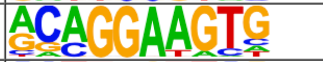 | ETS1(ETS)/Jurkat-ETS1-ChIP-Seq(GSE17954)/Homer | 1e-5 | -1.366e+01 |
| 24 | 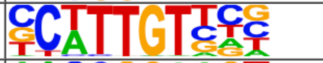 | Sox10(HMG)/SciaticNerve-Sox3-ChIP-Seq(GSE35132)/Homer | 1e-5 | -1.176e+01 |
| 25 | 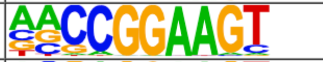 | ETS(ETS)/Promoter/Homer | 1e-5 | -1.157e+01 |
| 26 | 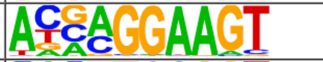 | ELF5(ETS)/T47D-ELF5-ChIP-Seq(GSE30407)/Homer | 1e-4 | -1.137e+01 |
| 27 | 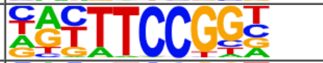 | Elk4(ETS)/Hela-Elk4-ChIP-Seq(GSE31477)/Homer | 1e-4 | -1.130e+01 |
| 28 | 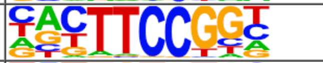 | Elk1(ETS)/Hela-Elk1-ChIP-Seq(GSE31477)/Homer | 1e-4 | -1.124e+01 |
| 29 | 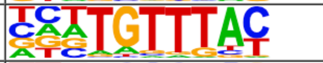 | FOXK1(Forkhead)/HEK293-FOXK1-ChIP-Seq(GSE51673)/Homer | 1e-4 | -1.031e+01 |
| 30 | 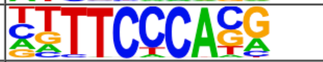 | Rbpj1(?)/Panc1-Rbpj1-ChIP-Seq(GSE47459)/Homer | 1e-4 | -9.257e+00 |
| 31 | 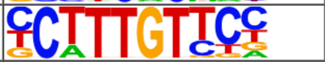 | Sox4(HMG)/proB-Sox4-ChIP-Seq(GSE50066)/Homer | 1e-3 | -7.520e+00 |
| 32 | 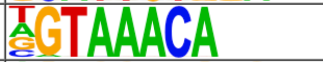 | Foxo3(Forkhead)/U2OS-Foxo3-ChIP-Seq(E-MTAB-2701)/Homer | 1e-3 | -7.292e+00 |
| 33 | 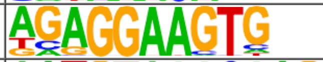 | PU.1(ETS)/ThioMac-PU.1-ChIP-Seq(GSE21512)/Homer | 1e-3 | -7.022e+00 |
| 34 | 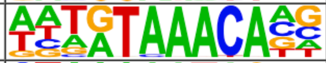 | FoxL2(Forkhead)/Ovary-FoxL2-ChIP-Seq(GSE60858)/Homer | 1e-3 | -7.001e+00 |
| 35 | 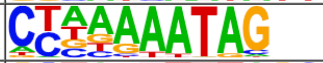 | Mef2a(MADS)/HL1-Mef2a.biotin-ChIP-Seq(GSE21529)/Homer | 1e-2 | -6.681e+00 |
| 36 | 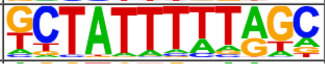 | Mef2d(MADS)/Retina-Mef2d-ChIP-Seq(GSE61391)/Homer | 1e-2 | -5.543e+00 |

**Figure S6. A** Sequence logos used to guide motif analysis alongside the source of each logo. Where data comes from ECs the EC type is also recorded**. B** Exact sequences assigned as motifs for each transcription factor.

#### Figure S5

| Rank | Motif | Name | P-value | log P-pvalue |
|---|---|---|---|---|
| 1 | 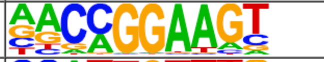 | ETV1(ETS)/GIST48-ETV1-ChIP-Seq(GSE22441)/Homer | 1e-10 | -2.344e+01 |
| 2 | 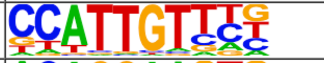 | Sox17(HMG)/Endoderm-Sox17-ChIP-Seq(GSE61475)/Homer | 1e-9 | -2.156e+01 |
| 3 | 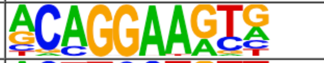 | ERG(ETS)/VCaP-ERG-ChIP-Seq(GSE14097)/Homer | 1e-8 | -2.063e+01 |
| 4 | 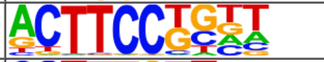 | Elf4(ETS)/BMDM-Elf4-ChIP-Seq(GSE88699)/Homer | 1e-8 | -2.057e+01 |
| 5 | 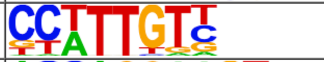 | Sox3(HMG)/NPC-Sox3-ChIP-Seq(GSE33059)/Homer | 1e-8 | -2.009e+01 |
| 6 | 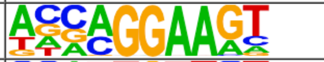 | EHF(ETS)/LoVo-EHF-ChIP-Seq(GSE49402)/Homer | 1e-8 | -1.986e+01 |
| 7 | 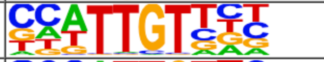 | Sox6(HMG)/Myotubes-Sox6-ChIP-Seq(GSE32627)/Homer | 1e-8 | -1.972e+01 |
| 8 | 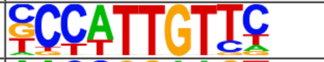 | Sox2(HMG)/mES-Sox2-ChIP-Seq(GSE11431)/Homer | 1e-7 | -1.768e+01 |
| 9 | 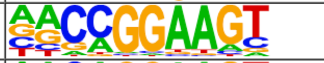 | GABPA(ETS)/Jurkat-GABPa-ChIP-Seq(GSE17954)/Homer | 1e-7 | -1.713e+01 |
| 10 | 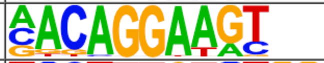 | Ets1-distal(ETS)/CD4+-PolII-ChIP-Seq(Barski_et_al.)/Homer | 1e-7 | -1.712e+01 |
| 11 | 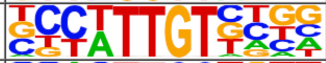 | Sox21(HMG)/ESC-SOX21-ChIP-Seq(GSE110505)/Homer | 1e-7 | -1.694e+01 |
| 12 | 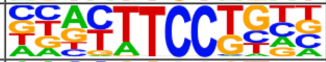 | Etv2(ETS)/ES-ER71-ChIP-Seq(GSE59402)/Homer | 1e-7 | -1.681e+01 |
| 13 | 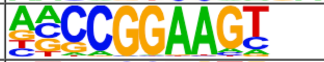 | ELF1(ETS)/Jurkat-ELF1-ChIP-Seq(SRA014231)/Homer | 1e-7 | -1.672e+01 |
| 14 | 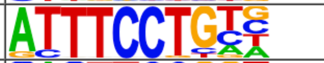 | EWS:ERG-fusion(ETS)/CADO_ES1-EWS:ERG-ChIP-Seq(SRA014231)/Homer | 1e-7 | -1.658e+01 |
| 15 | 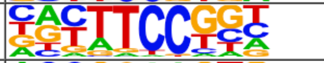 | Fli1(ETS)/CD8-FLI-ChIP-Seq(GSE20898)/Homer | 1e-7 | -1.656e+01 |
| 16 | 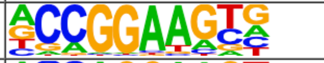 | ETV4(ETS)/HepG2-ETV4-ChIP-Seq(ENCODE)/Homer | 1e-7 | -1.621e+01 |
| 17 | 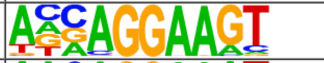 | ELF3(ETS)/PDAC-ELF3-ChIP-Seq(GSE64557)/Homer | 1e-6 | -1.570e+01 |
| 18 | 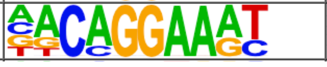 | EWS:FLI1-fusion(ETS)/SK_N_MC-EWS:FLI1-ChIP-Seq(SRA014231)/Homer | 1e-6 | -1.558e+01 |

| Rank | Motif | Name | P-value | log P-pvalue |
|---|---|---|---|---|
| 19 | 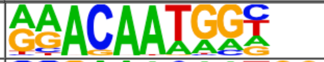 | Sox15(HMG)/CPA-Sox15-ChIP-Seq(GSE62909)/Homer | 1e-6 | -1.549e+01 |
| 20 | 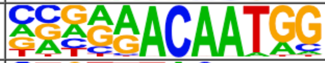 | Sox7(HMG)/ESC-Sox7-ChIP-Seq(GSE133899)/Homer | 1e-6 | -1.410e+01 |
| 21 | 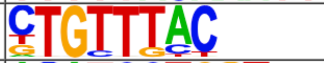 | Foxo1(Forkhead)/RAW-Foxo1-ChIP-Seq(Fan_et_al.)/Homer | 1e-6 | -1.408e+01 |
| 22 | 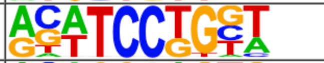 | SPDEF(ETS)/VCaP-SPDEF-ChIP-Seq(SRA014231)/Homer | 1e-6 | -1.406e+01 |
| 23 | 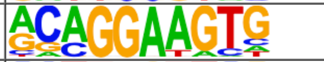 | ETS1(ETS)/Jurkat-ETS1-ChIP-Seq(GSE17954)/Homer | 1e-5 | -1.366e+01 |
| 24 | 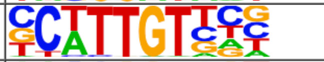 | Sox10(HMG)/SciaticNerve-Sox3-ChIP-Seq(GSE35132)/Homer | 1e-5 | -1.176e+01 |
| 25 | 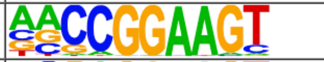 | ETS(ETS)/Promoter/Homer | 1e-5 | -1.157e+01 |
| 26 | 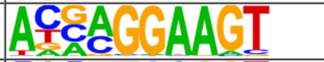 | ELF5(ETS)/T47D-ELF5-ChIP-Seq(GSE30407)/Homer | 1e-4 | -1.137e+01 |
| 27 | 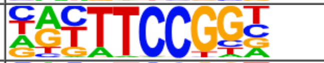 | Elk4(ETS)/Hela-Elk4-ChIP-Seq(GSE31477)/Homer | 1e-4 | -1.130e+01 |
| 28 | 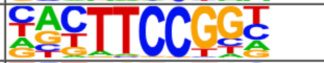 | Elk1(ETS)/Hela-Elk1-ChIP-Seq(GSE31477)/Homer | 1e-4 | -1.124e+01 |
| 29 | 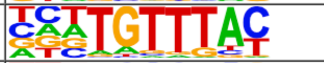 | FOXK1(Forkhead)/HEK293-FOXK1-ChIP-Seq(GSE51673)/Homer | 1e-4 | -1.031e+01 |
| 30 | 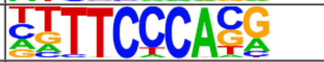 | Rbpj1(?)/Panc1-Rbpj1-ChIP-Seq(GSE47459)/Homer | 1e-4 | -9.257e+00 |
| 31 | 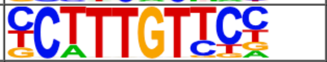 | Sox4(HMG)/proB-Sox4-ChIP-Seq(GSE50066)/Homer | 1e-3 | -7.520e+00 |
| 32 | 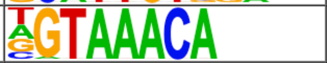 | Foxo3(Forkhead)/U2OS-Foxo3-ChIP-Seq(E-MTAB-2701)/Homer | 1e-3 | -7.292e+00 |
| 33 | 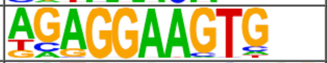 | PU.1(ETS)/ThioMac-PU.1-ChIP-Seq(GSE21512)/Homer | 1e-3 | -7.022e+00 |
| 34 | 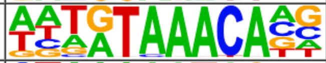 | FoxL2(Forkhead)/Ovary-FoxL2-ChIP-Seq(GSE60858)/Homer | 1e-3 | -7.001e+00 |
| 35 | 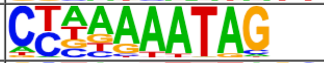 | Mef2a(MADS)/HL1-Mef2a.biotin-ChIP-Seq(GSE21529)/Homer | 1e-2 | -6.681e+00 |
| 36 | 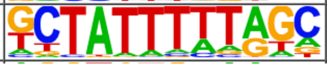 | Mef2d(MADS)/Retina-Mef2d-ChIP-Seq(GSE61391)/Homer | 1e-2 | -5.543e+00 |

**Figure S7.** Genomic regions around eight previously described arterial enhancers (red dashed box) alongside tracks showing ChIP-seq/CUT&RUN signal for ERG [11], ETS1 [12], SOX7 and SOX17 (this paper), FOXO1 [13], RBPJ [14], MEF2C [15], SMAD1/5 [16], SMAD2 [17] and NR2F2 [11] in HUVECs, alongside FOXO1 [11] and MEF2A [18] in adult mouse hearts.

### Figure S7

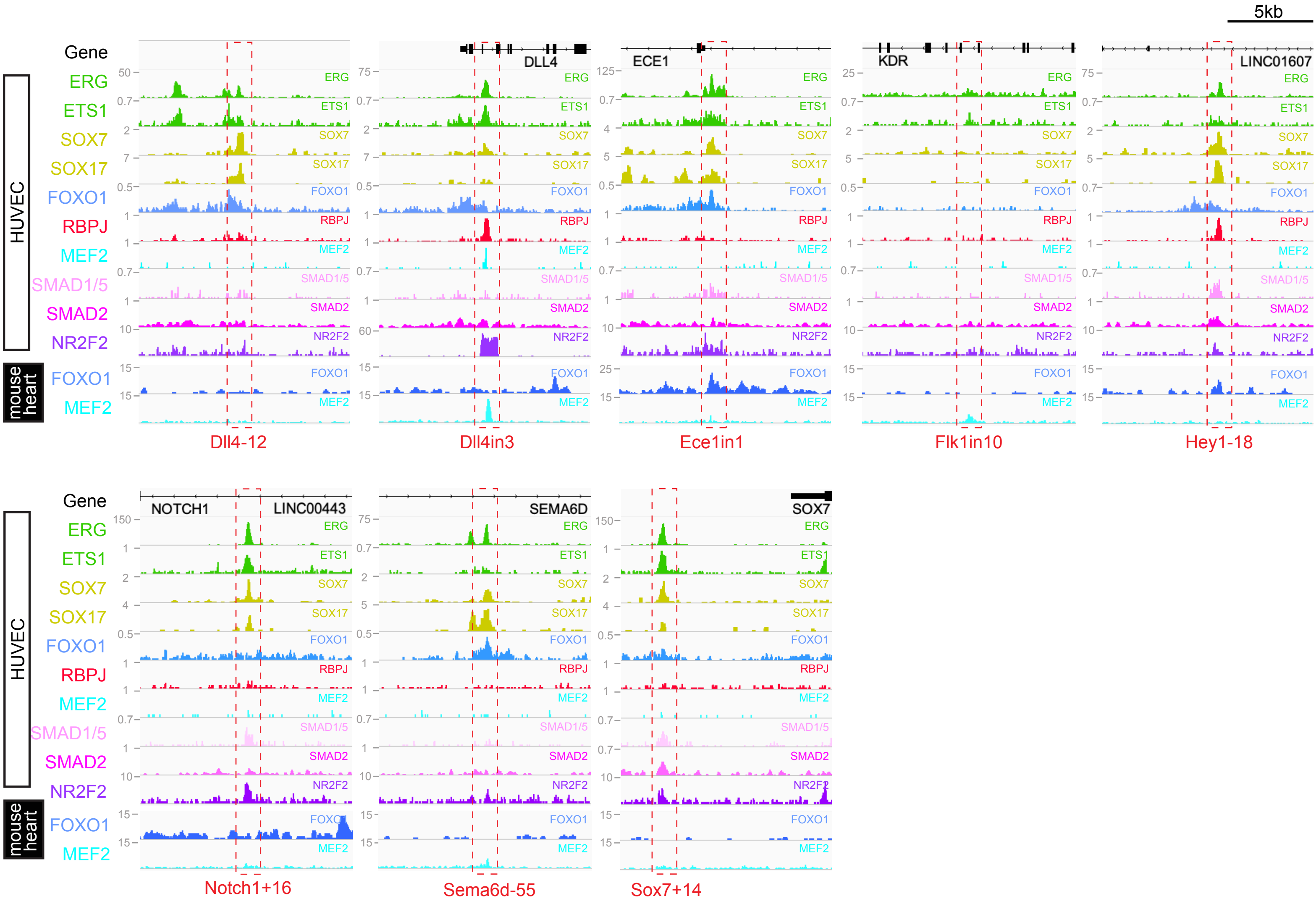
5kb
Gene
ERG
ETS1
SOX7
SOX17
FOXO1
RBPJ
MEF2
SMAD1/5
SMAD2
NR2F2
HUVEC
mouse heart
FOXO1
MEF2
DLL4
ECE1
KDR
LINC01607
Dll4-12
Dll4in3
Ece1in1
Flk1in10
Hey1-18
NOTCH1
LINC00443
SEMA6D
SOX7
Notch1+16
Sema6d-55
Sox7+14

**Figure S8**. Sequences of each core enhancer listed in Figure 5 alongside annotated transcription factor binding motifs. ETS motifs highlighted in green, SOXF motifs in yellow, FOX motifs in darker blue, FOX:ETS motifs in turquoise, RBPJ motifs in red, MEF2 in bright light blue, SMAD4 motifs in light pink, SMAD1/5 GC-rich motifs in darker pink, TCF/LEF (Wnt pathway) in grey, NR2F2 motifs in purple and KLF4 motifs in orange. The sequences assessed as transcription factor motifs are listed in Figure S6. **Bold underline** indicates that motif is conserved to the same depth as surrounding sequence (conservation depth indicated after title of each enhancer), **bold** indicates motif conserved between human and mouse, *italic* indicates motif just found in mouse sequence.

#### Figure S8

ETS SOXF FOX FOX:ETS RBPJ MEF2 SMAD4 SMAD1/5 TCF/LEF NR2F2 KLF

**Acvrl1+6 core**
Conserved to opossum
>CTGCCCTGAGCTCCCTGCTGTGCGTGACTTCAGCCTGTTCTATCCAGGCCTCAATCTAAACAATCTTGATTCCTGTTGCCGGCCTGGCGGGA
CCCTGAATGGCAGGAAGTAAGGACAAGAGCCTGTTTATGTTTGAAGCAGCCAGGCTGGGGGTGGGGAGTGGGGGCACTGGGAAACGGTGGGCA
GGGGTGGAGGCTGGAGCGATGGGCAAACGGCTGAGGACAAGAGATGAGCTATGA

**Cxcl12+269 core**
Conserved to opossum
>ATCACAGGCCCATTGTGTGCTGGGCGGGCTGCCAGGCGCCTGCTCCACACAGGGCTGTGCCACATCCCCACTTTCCTGGAATTCAAGGAGTA
AGAGTCACACCAGCTCCTGATTCACTCCCCAGTTATTTCCAGCTTCGTTTTTTGGGCTCCCCCTATAGCAGTGTTTTCTCAGGAGCAAACTCT
GGTGCAATGAAACTACAGAAAGTCCACATGCTGTACACGGGGTGTTTATCAAGAAAACAATTGTAGAAAAAGACTCTGCAACTTTGC

**Cxcr4-194 core**
Conservered to cow
>CAGCCCTTTAACCTTTTTACTAAATTTTCTACTGTCAACAAATAGCACAATGGTAGGAGGCTTGAAATATGTTTCCTTTTCCTGATCTTTAGT
AGAGCATTGCTGGGTTCAGTGAAAGACTTGAAGGAGGGGCAGGAAAAGACAAGTTCTATGTGCCAGTGGTGTCAAAGGCTATTTATAAGTTCA
TTGAATGTAGCTCTTCCAGCTGTGTGTGTGGGGGGGGGAGGTAAAAGGCTTTCACCCTGTACAACCCTTGAGGAAAGCCAGGTCTTAGTTTTG
TCTATACATGGACACAATAGTAAACACTTCTAAACCAAAAAGCACCTCTCAGGTTCCTTTGGCACACATTCTATGAGTTGGTGAAGGGAAGGA
ATCAGTAGAGTGTATGGA

**Cxcr4+135 core**
Conserved to chicken
>CCTCTCTGTTCCCACAGACTGCTATCCAAAGAGCGCCGCTCCAACTCGGAAGGGAAGGATTTGCCCCGGCTTGGAAGGAAGTAGAATAAATA
GAAACATAAGTGGCAAGGTCACCTTTAATATCATACGAACAGCTTCTCAACAAGCCTCACAATTGCTTTGTTTCTTCCTGCTCCTGTGCTATA
AACAGGCAGAGTGCAGGCAAACTCTTAGAACACACACAGCGAGCAGAGGAAGCTCCCAGAGTCCCGAGGAAGGGGGAGGC

**Cxcr4+151 core**
Conserved to chicken
>CACAAAGCTGTTGTGAAATCAAATACAAATCCTTCCTGAGTATTTCCAGCTTTCCTTCCCGGAGGAAGTTTTCAGTCAATGGGAAACAAAGC
CTTATTCAAAGGATTAAAGCCGGAGTCATTCCATTGTGTAAGGCTTCAATAACACGGCTTTCCTTCCACGGACAGAATTTTCAATTCACAAAT
CATCTCAAAAACAGGAAATTTTGGGGAAACACGTGCTTATGATAACAAACAAGGATCTATCC

**Efnb2-112 core**
Conserved to opossum
>ACTAAACCGTAACACCTACGCACACAGTGATTGAGTTATAGTCAGAAAGGCACCATAAATGACATTTCCTCCCACTAATTTCACATTGCCTG
TGCAGCTCTGGGGAGTAACATTACTCTTCAACAGAAAAATGTCCTATCCCACCCTGGCAACGCCTCACAGCACTTGTATTCCCGTCAGTTTTT
TTTTTTTATCAGCTTGACTTGCCCTGTTTCTTCAGGGGTAACTTGTTCTCTGATGCTAGTCTGCATTTTATGGAGTGTGGAGTCATAGCAGTT
TCCTTGTCCTGGTCCCATTAGCTTCCTGGTCCTATTGTTTGGAGTCTCCAGATCCTGATGTGTAATCCAACTGTTTCCCCACTATGTGGGAAG
CGGAAAGCTGCCGGCCTATGTCTTGAACAACCCAAGCTAGA

**Efnb2-141 core**
Conserved to tenrec
>GAAAGTCTCATGAGAACTTCTAGAAAGGAAACACACTCTGTCTGTCACCAACAGAGGGAATAATGTCACCGGCCCCGTAAGCACTGACTGGA
ACAATGAGAGCTTCCTGGTTCCAGGAGTCAGAACAAATGTGTGAAATGGGATGGAGGGGCATTGCCACCCGAAGCTCTCACAGGAAATTGCTC
AGTCAATGGAGGGCTCCGTGTCAATATTTGGGTAAGGAAGCATGTCTCAGTCGGATTCCAAGGTGCCCCACAAGCAAGAATTCCTAAGAGACA
TTATGCTGTACTTGGGTTCTGGGCTGCCCATTCTCATACCAATGTCATGTGTGT

**Efnb2-333 core**
Conserved to chicken
>AAAGAGGGGGCTCCAGGGCGGACATAAAATTAACTTCTCTTTCAGTGTGCTAACCTTTTGGTTTTAATTTATTCACTTGTCAATCTTATGAA
TGCCTGGTGAACTCAGGCTATTGATTTCCGTGGCCAATGCTATTTTTCCTGTTTATTATAATATATTGTCCCATTGTTACCAGCGCTTGCACT
CAGCTAGATTTCCCACACAGTGGACAGGAAACCGTTCCATGGCATCAGACTTCCTTAGATTGATTTTCCAGAGGCTGGGAGGGACCTGCCCAC
TCCTAGCTCTTTGTGCTGGGACA

**Gja4+50 core**
Conserved to opossum
>CTTACCCACCACTTTCCTTCCCACCCTGTGTGGTGCTGCCAGTTAGCACAATAGTGTATTCTGTGTCCAAGGGCCTCTGCTGGCGGCCCCCA
GGCTTGAGCAGCCCCCGTAGACAATGGTAGGGCCCGAAAGGCCTGATAGCCGGTGACAAATGTATGTGAATGACCAGCTCACGGGAAGTGGCC
TTCAATAGTCCGATAGCTTCCTGTATCCTGACATTGTGAAGGCCCACTTCCTGCTCTGTCTCCGAAAATACAGTCAGGCAACCCGGGGTGGGG
GTGGGGGCAGGGGCAGAGGCTGGGCTCCCCATTACCCCTTCCGGGCTCCTACAGTATTCTGAGGACAACACCAGTCTCACTTGCCT

**Gja5-78**
Conserved to chicken
>GGTCCAGGGTTCCCAAGTTCAATGGTCTTTTCTTTTGTTGCTGTAGAAAGATTGCTGAGGCATGAGCTACTTCCGTCCTCCCTTGGCATCTT
TGCACTGTGCCCCAAGAAACAATCATATCACATTCAGGAAGGCTGGGCCAGCAATGGCAGGGAAACAGGAACTGTCCCCACGGATGTCCTCCC
CAGGGGAGACAGCCAGGGACTGGGA

**Gja5-7 core**
Conserved to opossum

>AGGGCATTGTGCTCTGGAGCTGGCCAGCTTCGCCCTGCCCTGGCCCCAGCACCCCAACTTGTAAAGAAGGAATTGCGAGGTTATCTAAAAGG
AAGCCAAATTGTTTTCGGAAAACAATAGCCAAGTAGCTACAAGTCATTAGTTTCCTCCCAGGGGCCTAAAATCAAGCTTGTCTGGATAACTGA
AAGGGCTGTTAGCACTGTTTACTTAAAGACCTCTGGACAGAACCCATAAATTAATTCCTGTG

**Nrp1+78 core**
Conserved to opossum
>GCGGCTCCTTCTCTTGTTTTGAAGCTTCCTGCAGACAACAGATCTCACAGGAAGTGCCAGTGTGGACGTTCATATCCATGGCTAACGATGCA
TTGTTGTTTTTTTTTTTAAATGTTACTTGGTAATTTCCTAGAGAAAACAATGGGAACCGAAGGCTGTTCCTGTCTTTAAGGAAAGCCAGTTTC
CAATGTTGCTTTATTAACTTAACACATGCAAGTATTTGTTCTAGTTGTTGCTACAGCAACGTGTTTTGTGGGGGCATATTTCATGCAAAATGG
GATAAAGCTGAAAATCAAGACAAATGTCATCT

**Unc5b-57 core**
Conserved human-mouse only
>CTTCCGACCCTGCACTCAGTCTGACCCCGGAACACACCACCTTCAGCCACTTCCCACCACCGGGTCACTTCCTGGCCCACTCAGCAGAAGAG
CGCCAGTCCTTCTTCCACTCTTCCCACCCTGGGGGCACCTGCTTCCTGCTCTTCCTCTGCCTCTGTGCCCTAAGCTCAGGGTCCACTCGCAGC
TCAGGGGTCACTTCCTTGTGACCTCACCCCATCTGTTTGTCAGGAGACAAAGCAGGGGTTTCAGGATGGCTTCCTGAATCTGGTGCCTCAGGC
TTGGCTGAACTTGAT

**Unc5b+30 core**
Conserved to tenrec
>GGGGTCAACAATGCCAGGAAATAGCCAAAGGAAACCTCTGGGATGGGCCCCTGGAGTCTACTCAATGGCGTGGGCCCAGCTTGGGGAGATTT
TTCTCACGAGTCCTCAGACCCCCGGGTCCCCATTAAATGAGCTGACCTGAATGAAGGGAAGATTTATTTCCTGGCTTCGGAGTGGGCTCTGTT
CCTCGCCCCTCACTTTCCTCCTAGCCAGAGAACAATGAGGGTCCCCAGAGGTACTCTTCCTGTGGGAGCAGGTGCAGCCCC

**Unc5b+39 core**
Conserved to opossum
>TCTGAGCTTTAAGAGCCACCCGGAAGCAGGCACAAAGGGAATTGTCCTGAGCGCCAGGCAGTCCTCAGCTCCCAGCTGGGGAGGAAAAGCGA
GACCTTTTTGTCCAGGGCCAGGGCTGAGACAGTGGCCAGTGACGTGCCTGGGAAAAGCTGTAGTCCTCAGCTAAAATTAGCCACCCACCGAAC
CTCTGCATCAAGGGCAACCCGCCGCCACCGACACAGCTATTGTTTTGAACTTTGGATCCCAGGGGACTTGAGGAAGTTGGT

**ARTERIAL ENHANCERS IDENTIFIED IN OTHER PUBLICATIONS**

**Dll4-12 core**
Conserved to opossum
>CTCCCCTACAGACAGGGTGACGACATTGTTGTTCTTATACTACAGGGCCTTCCGCCTGAGGTTCCGGCTCCTAAATGGTGGGAATAAGGGTC
CCCAGGACAAAGCCGGCCTGGTTCCCGCAGTCACCTCAGAATGGACGGAATCCCCATTGTGTATGGTCGCCCCATGCCCGCCCTCACTTACCC
TCACCGGACATGCCAACAAACAGCTCATTGAGCCTGGGGAGGGGCCGGGGAGACAACAATGCCCCCCAGAAGGCAAT

**Dll4in3 core**
Conserved to chicken
>CCTGGACTCAGAGCACAATTGCGTTTCCTGCGGGTTATTTTTGGCGTGGGAACGCGGGGAGCACGGCGGTGAGAAAGGCCGAGGCTGCCAGC
GCCGCTGACGGGCCTCTTCCTGTATTTTACACCTTTTGCGAATTCCGCTCCTTTGGAAAGGGAATAATGGCTTTGGGATGTTGTTCTGACACA
GAGGAAAAGGATATTTCACCAGCACAACAATTCTCACTTTGAAAAGGAAAAAGAAAAACCATTACCTACGTCTA

**Ece1in1 core**
Conserved mouse-human only
>GGAGCCCACCAGTGAGTGAGACCTGGGTCTCAGGGGATGGTCAGGGAAGACAAGAGGCGAAATAATATCTCCATTCATGCAGGTCCTCTAGG
GGGGAAAAATGTGCCAGGCCCCTGTAATCTGGCTAAACAGGAAAGAAAGCACAATGAGGGGAAACGGAGGGTGTGGCAGGCCCTTGGGGGCAG
GGGACCAGAGGGGACAGGCTTGTACTCAGGACCCCAGGTCCTGGCTTTTCACACCCAAAGCCAGCACGGAGAT

**Flk1in10 core**
Conserved to chicken
>GGAGGGAAGCAGCTATTCTGGGAACAAGTCTCCATTCATAAATTGGATCGACAAGACAATTCAAGCTCACTTAGTTCAAAGGAAGGTAAGGA
AACTTGGAAGCCATTGGGGCTTCTTAAAAGTCACCTCTCTGGGACGGACCGACTGCGGGCTTTTGTTTAGGAAATGGCCAGCAGCAGAGGAAG
AAACTCGGGTTTGCTATTTCAAAAACAACAACAGGAAGTGGAATGCTTGGGGTGGTAGGTTGAAGGTGGTGGCTGTTGTTTTCCCTAAGGATG
TCTGCACTTGTGGTAGACCTTGATAAGCCTGGGCCTGAGACTCTCGAGGCCTGGGCTAGGTTTATCACTGCCTCGCATCCACCAGCACTTCCT
CTGAGAGATGGACATTCCCACAGATAAGGAGGAGCAGTGTGGTCCTC

**Hey1-18 core**
Conserved to chicken
>AGGGGAGGTGGAACTGTGGCGCGGCCGGGGTTCCCACGCTCCTATTGTTGGCGACTTTGTTTCCAATTCCCATCACGTGTGCTAATTAGTTG
GAGCTGCTGGCTGGCTGGAAGGCGTGTGGCTTGCCTGAAAGCAGAAAGATCGCTTCCTGTTGGGAGTCGGGCAGGCAGGACTCTAGCTCAGAA
G

**Notch1+16 core**
Conserved to opossum
>TGCAGGGCTGGCCAGGATTGAAGCCAGTGTCCAGCTTTCCAGCTCAAAGAACCACTCCCAGCACACTCAGCTCTTTTGTCTTCAAATCTTTC
CCCACAGTTGCATCCTTGGGTGGTTTAAAATACGCTGAAAAAGGAAGCTCAGGAAAAGCGCAGGCTTCCTGGCCTGGCTCAGGGAAGCCCTTT
GTGTGGGAGACAGTTGAAGTTGGAACCTGCCCTCTAGGGAAGCCACCTTTTCCCACTATAGGGAAAACAATACTCCCCTCCAGG

**Sema6d-55 core**
Conserved to cow
>GTCAGCCAGCTGTTAGAGGAAATTTACAAGACACCTTTTCCTTAAATCTACAAAAATTGCTTTTGTCAACAAATTCCTGTTCAGACTTCCTG
CTCCAGCAGTGTTTGCTTCCTTGCTGCTGTGAGAGGAAACAAAAAATAGAACGGCCTTCCTGCTTCAGGAAGTTGGGCCCAGACAAGTCTGTG
TGCCATGATGAGGCCAGCATCCTATGAAAGCCTCTGGGAAGTAATCCATTGGGGTATTATATTTAGGAACAGCCTGACTGAGAGCATCCCTGT

GGCATTGAAGGGATAACCTTGTGCTCAGTTGGCTTGTCCTTCATTTCTGTTTAAAAGAATCGGGAGGGATGCAGCGTACAAAATGCTTCTGAA
TGGAGCAAGTCCTCTAGGCTTGATGGCATCTCTTCAGAATGGGAACTGTGTAGGAAGTGCA

**Sox7+14 core**
Conserved to opossum
>ACTCAGGCGGACAGAGAGGAAGGCACAGAGCTCTGCTAATCTGGAGAAGAGATTTTCCTCCTAGGAGGCCCTTCCTTGGGCAGGGCTCTTTT
GGTGGCGGCCGAGGTGGCTGGCCTGAAGGTGGTGGCCAGGCTCTATAGGGAGAGGCTGCCTTATCAGACAGGAAACACTGGGCTCTGGTCACT
TCCTCCCACTCCTGGTCACAGCAGCTCTCAAGGCAGGAAAAGCTCGGCAGCTACTACTCA

**PAN-EC ENHANCERS IDENTIFIED IN OTHER PUBLICATIONS**

**Apln+28 core**
Conserved to opossum
>CTTTTGCTTTTGCAGCTGCGGTGGAAGTGAGGAAAGCGGGAGGGCATCCTGAGGCGGGAAATGCCTATCGGTGTCATCCCAGCTGCTTCCGC
CTTCCTCCTTCAAGGATTCCTCAGCCTCTCTGTATCCCTGCCAACCCCTCTCCCCTCCCCTGAGTTCTGGCCAGCAGCTGTGCATTCCCCCTT
TAGGGCCCACTTCACAGATCCTCTGAGAGGCTGTCAGTTCCCCATTGTTCTCGGGGCTTCTCCGGCATTCCTTAGGCTCATGAGGGGAAATGC
CAGAGGCTTGGCCCCCACCTCCTCAGAGATGCTTGGCCTACTAGTCACC

**Cdh5-1 core**
Conserved mouse-human only
>GAATACCCCAGGCAGGTCCAAGGCTTCCACGGGCCCAGGCTGACCAAGCTGAGGCCGCCCACCGTAGGGCTTGCCTATCTGCAGGCAGCTCA
CAAAGGAACAATAACAGGAAACCATCCCGAGGGGAAGTGGGCCAGGGCCAGTTGGAAAACCTGCCTCCCTCCCAGCCTGGGTGTGGCTCCCCT
CTCCCCTCCTGAGGCAATCAACTGTGCTCTCCACAAAGCTCGGCCCT

**Egfl7-2 core**
Conserved to chicken
>CAGGCTTCCTGCAGCCCCAGGCCCTGACGTAGTTCCCCTTCCTCTCAATGGTGCTTCCATTGTTCTCAGGGGCCTGATTCCCCATCACCCAC
CACACACACACTTCCTGTTTCCTGGCCCAGAGGAAACAGAGATTTCTGGCGGGCCACAGATGCTTGTTCCCCCCACCCCCAAGCTTTTAAAGG
ATATTAAAAATAACAAGAGTAGACAGCTTA

**Egfl7-9 core**
Conserved to opossum
>AGGAAGGCGGGCAAAGTAGAGCAGCATTTCCTCTTGGGCCAGGGGGAGCAAGTGCAGGCCCTGATAAGGCACACATGGCCCTCTGCACGTCC
GCCTTTTGTCCCAGCCCCACCAGCCTTTCCTGCTCATGTCTGCCAGGGCGGGAGGACTGGGGCCTGCCTGTTTCTCTCTTCCCTTTGCATCTC
TCAGCCTGTG

**Eng-8 core**
Conserved to opossum
>GGAAGCCCTGTGTTCACAGTCTGCACATGTTCAGAAGGGCTCCTGGCCCCCAGCCCTCCAAACACCCCCACCGTGGAGCCGCCAGGCCTTCC
TGTTTACTCGAGGCCCCTCTCGGGTCTGGGAGGCGACAGCCTTGGCCTCCTCTGCTCCCCACTTCCTTCCTCTCCCACTTCAGTGCCTCCGGC
CCAGGCACTCAGGGGGCCCTGAGGCCAAGGCAT

**Fli1+12 core**
Conserved to chicken
>GCTAAGTTATGTTTGCTCCCAACGATCCTGAGAGCTATGGTTTTCTTTGTTCTACCAAAGGGAAAAGGAAACATATCAGGAAGCTGGGCCAG
TCCACAGGAAGTCTCACTTGGGCTGATAAAAGGAACTGATGTGCAGTGGCTTGGGATGCAGTTATAAGTCAATCTGCCTTGAAATTACAAGAA
ATGCTGTTCAGGAGAAACTGTAGCCTGAAGTCAGCCTTGCTCGGGTCTTTCCTTTACCT

**Flk1+3 core**
Conserved to chicken
>AAATGTGCTGTCTTTAGAAGCCACTGCCTCAGCTTCTGCAGCTCAGATACCAAAGGAAGTCTGGTACACAGCATGATAAAAGACAATGGGAC
GGGGTCACAGTGGCTCCCGTCCCTTTCAGGGGTATGGAGACGAGCTGTAGAGAGATGTCTCCAGGGAGTTTTCATTAATCAGCAATTTAGTCA
GATCTGTGCATCCTATGCTTTACAAGAAATGTCAGTGGGCCTGAGATCATCAGATGGAGGTTCATCGGGTTTCAATGTCCCGTATCCTTTTGT
AAGACCTTGAAGTTGGCAACGCAGGAAAACAGGAACTCCACCCTGGTGCCGTGAATTGCAGAGCTGTTGTGTTGGTTTGTGACCATCTGCCCA
TTCTTCCTGTTATGACAGAGCTTGTGAACTTTAACTGGGACTGGGGCAAAGTCAATCCC

**Gata2+9 core**
Conserved to xenopus
>TTTATTATTTTTCCATGGAGTCACCTATACTGTGTATTTTCATTTGAGTGATTTTTTAAAAAAATGTCCTTTCGGATCTCCTGCCGGAGTTT
CCTATCCGGACATCTGCAGCCGGTAGATAAGGAAACTTCGTGTATCTGTTTCCGGACCGGCAAGTTTTCAGAGCCACCTCGACTCAGTCCTGC
CTCTTGCTGGGCTGTTTTGAAATTTCTAATACCCTCCACTCTGCAAATAATGTGTAAAATGCTAAGAATAATAAATATATTTTTTCAGGGCGA
AGTGATTTATGAGTTTAAATCGTTCACCCGCT

**Mef2cF7 core**
Conserved to opossum
>GTGCTGCCGAGCACACTCAGCCTGCTCTACTCAGAGAAGGAAGTGGAGAGTTTTGGTTCCCATTTTACACTTGTGGAGCAGTTTAGGGAAGG
AAGCACCTTTACACCTTTTATTGTTCGCGGTTGCTGACATCATATCCTTCCCTGGCCAGTCTGTGCGTGTTTCCTGCCATTCTGTCAAGATTC
TTCTGTTACTAGGAGGGATTTGCTGAAGGGAAGGAGGG

**Notch1+33 core**
Conserved to tenrec
>ATCCCAGTTAGCTTTCCTGTTTATGGCCATGGTTGGCTCCAGGAGCTAAGCAAGGAAACCTTATCTTCCTTGTCTCCCACCATGTCCCCAGC
CCACTGGTGAGACTCTCCAGGGCCTCTCTCTCTATCGGTTTCCACGGTGACCTGGGCAGACAGGAACTTTGACAGAGTTTCCGACAATTGTGC
AAAGGGAAGCAGGAAGCTGGTCAGCGCGGCTTGACTCCTCCC

**Pdgfrb+18 core**

Conserved to cow
>GGGCTAAGAACCTGGCTCCTCTCTCCTGGGCCTCCTTTGCAGCTCTGTGCTGAGTAGGGGGGTCAGCCCACTCCCACATTCCACAGTCCTGA
CCCCCGCCCCCTCCCCACTTCCCTTCCCACAGACATCCCATTTCCTGTTGTGCTTGTAGAGGGAGCCGTGGGAGCTCTTCAGGCTCTGGCTCT
CCCCCTCCTACTCCTCCAGGCCTTTCATCACACTTTCCCAAATATTTGTCCTGGCAGATAAGTGCCTGGTATTGGAACTGTCACCGGGAGGAC
GGGGCTGTCAGAGCTGGCTGGCCCCCAATCTAGCCCAATTCCCCC

**Tal1-4 core**

Conserved to cow
>TACCCGGGGCCAGACTCCAGACTCCCTGGTTCCTCACCTCCCCGCCCCCTCACCACCCCCACCGAGGCGCTCCGAATTTCCTGCCCGACCGA
GGCCCGGCTCGGGCGGGTGGAGGAGGGCTGGCATTTCCTGGCCGCCGCGTCACTGGCTCAGCGGTGCTCGGACAAAGCGCTGACCGACAGGCA
CCAGAAGCTATTTCAGGCGGCGCCCAGCTTAGCGCGCAGTTTCCGTTTTTCCACCGTCGGAAACAGGGAACAGGGAGCTTGCAGACGTCACAA
ACCCCCAGCCTCAGGCGTGGGTCCAGGGACCA

**Tie1-1 core**

Conserved to cow
>GAGTGTTCAGCCTGTCCAGCTCAGTGCACCTCGGTGCTGGGGAGGAAGAGGAAATGATGGTTAGGGTAGGAGGGGTGGCAGGGAGAGAGGGT
GGGAAGAGTTGTCGCCTGTAGTCCTCCCGTGTCCAGCCCCCACAAGCCCGGGATGGGTGTGGCCTGGAAGTCTCTGGAAGGGGGGGCATTAGA
GGTGGGAGGCAGGTTGTGACAAGGACAGATCTGGGGATGGTTGGGCTCTCTCTCCCATCCCTTCGGTCCCTTCCATCTGCATTGCTGGAGCAC
GGGAGACAGGAAAGGGAGGAAGCCAGTGGCTTCCGCCTATTGAGAAGGTTTGGAGGCAGGACATTGTTCTGGGGTCTCCCCTCCTCCCAGCAC
ACACACGCTG

**VEIN ENHANCERS IDENTIFIED IN OTHER PUBLICATIONS**

**CoupTFII-965 core**

Conserved to zebrafish
>GCTGAGACAAATGGAAAGCTGAAGATAAGGATCCTCTGAGGTGCGAACATACAGCTGTTGGGAATTGCCAGAGAATCGGACCAATAAAGGAA
GTCACTATTTTTCCAGGCCTGAAGTGAGTTATAGGGCGAGACGGGTGTTGTATATTTATGTAAGGCAACAGCAGGGAGTTTAAGCGGCTGGAT
ATTGCTGAAAGAGCATCATTCACATTCAGGCGGAGACAAAAGGTGGAAATGAAGCAACATCCTGGCCAAAGAAGGCCTCAAGACAGAATAATA
ACAGTTCAGAGAGGGGGCTGTGTGCACGGCCGAGGGTCGGCCTCAAAACCAGGAAATGATCGAGATGCCTTGTCAGATCTTC

**Ephb4-2 core**

Conserved to manatee
>CACTTCCCCGCCGCCTGCAGCCTTGACCTCCAAGGCGGGGGTAGGGACCGTTGTGGCTCTTTCCTGAGGCTGTTTCCTGTCTGGCTCCTGGG
GGCCCTCGGGATGGCTGGGAGGGCCCTTCCTCTCATTTGCTAGCACCCCCTCTCATCCATCAGTTTGAGGGGAGGGTCCAGGAAAGACGGCCT
CCTATCTACATCAGGGCACTGTGAGTGTGGGGCACGGGATGGTTGGATGAGAGAGGTGCTGTTCCCGAAGTCGGTCCTTTAAGGGCTGCGGTA
AGGAGACTTTAATTTAAGGTAATTAGTACAGGGTCTGGAAACTCTGAGGTAGGAGTCTGGGGCACCTGGGAGTCTGCCAAATACCCTAAGGGC
GCACACACACACCCCAGCGGGCGCGGGAGACCGGTGATGACCTCTTGTCCGCCTGCGCGCACACACACACCAGCGGGCGCGGGAGACCCGTGA
TGGCCTTTTGTCCCCGTGCACTTATCTTCCTGGCGCAAGTAGTGCTCCCCACCCCCTGCCCTTCCTCACAGCCCTGCCTGGGTCCCGCTCCGG
GGTGGGGGGTCAGCCAGGGCAGGAAACAGCCGGCTTGGCTGGAGCCAGGCTGACCGGCTAGATCTGGGAGTCCCCTCCTCCTTCCCCACGCAG
ACTCAGGCTCCCCTTCTCTTAT

**Mef2cF10 core**

Conserved to chicken
>ATTTAAATAACATTTCAACAAATGTAACTGTTTGGAAACTACAGCTGAGGCAGCCTCCTGCCTTCTGTCTTGCAAGGATTGATTTGGAAGGC
CCTGAGGGAGAAAATATTGGAGTTGGGAGCTAAAACTGAAGCCACTGTGGGAAGGAAGAAGTTACTCTCTTCCTGTTATGACAGGAAAGCGTA
GACAAATGTGCTTCCTGAGTTAGCTCAGCAGGGAATGTAGGATTCTTTGACCTTTAACTTAGGTATATTTTGACCATGTTGAACTAGTAAACA
TGCTTTGGACTTCCAGATA

**Figure S9.** Genomic regions around 13 previously described pan-EC enhancers (grey dashed box) and three previously described vein enhancers (dark blue dashed box) alongside tracks showing ChIP-seq/CUT&RUN signal for ERG [11], ETS1 [12], SOX7 and SOX17 (this paper), FOXO1 [13], RBPJ [14], MEF2C [15], SMAD1/5 [16], SMAD2 [17] and NR2F2 [11] in HUVECs, alongside FOXO1 [11] and MEF2A [18] in adult mouse hearts.

### Figure S9

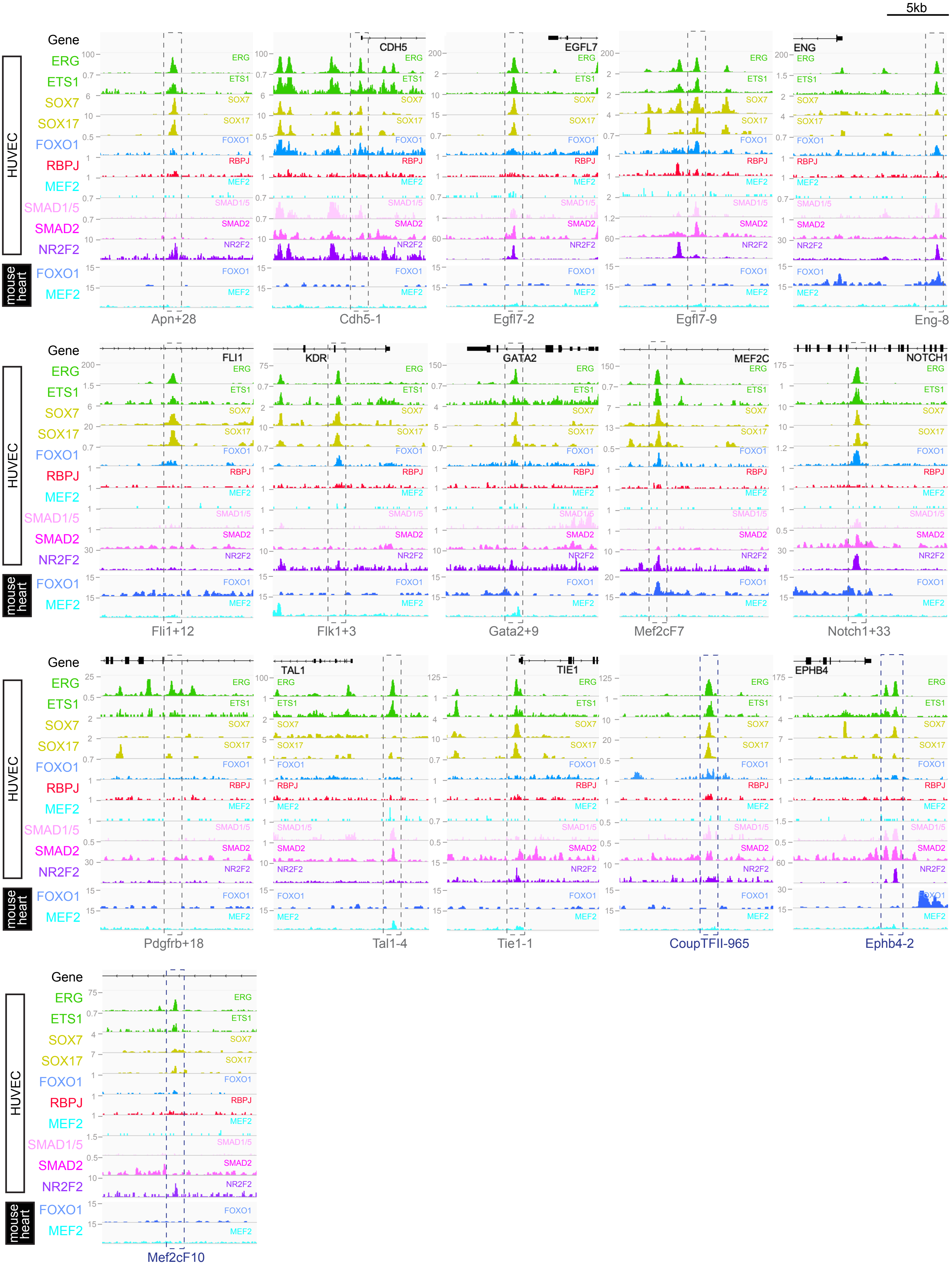
5kb
Gene
HUVEC
ERG
ETS1
SOX7
SOX17
FOXO1
RBPJ
MEF2
SMAD1/5
SMAD2
NR2F2
mouse heart
FOXO1
MEF2
CDH5
EGFL7
ENG
FLI1
KDR
GATA2
MEF2C
NOTCH1
TAL1
TIE1
EPHB4
Apn+28
Cdh5-1
Egfl7-2
Egfl7-9
Eng-8
Fli1+12
Flk1+3
Gata2+9
Mef2cF7
Notch1+33
Pdgfrb+18
Tal1-4
Tie1-1
CoupTFII-965
Ephb4-2
Mef2cF10

**Figure S10**. **A-B, D and F** Venn diagrams assessing overlap of genomic regions called as peaks of various combinations of ChIP-seq and CUT&RUN datasets. **A** Comparison of called binding peaks for ERG[11] and SOX7_mCherry[19] with marks for enhancers and TSS [11]. **B** Comparison of called binding peaks for ERG and SOX17 (new CUT&RUN) with marks for enhancers and TSS. **D** Comparison of called binding peaks for ERG and SOX7 (new CUT&RUN) with marks for enhancers and TSS. **F** Comparison of called binding peaks for the overlap of called peaks of our SOX7 and SOX17 CUT&RUN data with each another and previously published ERG ChIP-seq data.

**C**- **E** Top three motif families called by HOMER in the SOX17 (**C**) and SOX7 (**E**) CUT&RUN peaks.

### Figure S10

## A

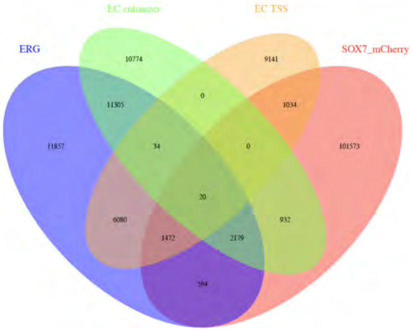

## B

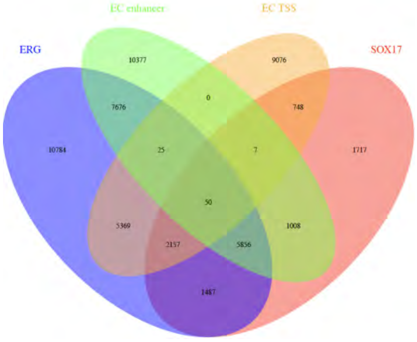

## D

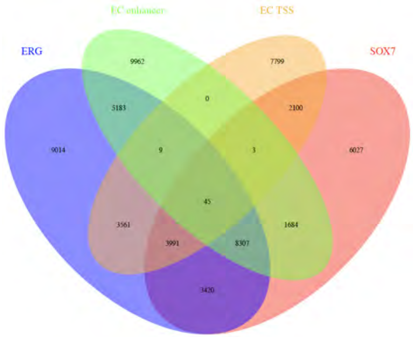

## C

| Rank | Motif | Best Match/Details |
|---|---|---|
| 1 | 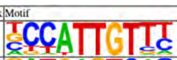 | Sox17(HMG)/Endoderm-Sox17-ChIP-Seq(GSE61475)/Homer(0.988)<br>More Information \| Similar Motifs Found |
| 2 | 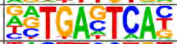 | Atf3(bZIP)/GBM-ATF3-ChIP-Seq(GSE33912)/Homer(0.987)<br>More Information \| Similar Motifs Found |
| 3 | 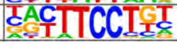 | ERG(ETS)/VCaP-ERG-ChIP-Seq(GSE14097)/Homer(0.967)<br>More Information \| Similar Motifs Found |

## E

| Rank | Motif | Best Match/Details |
|---|---|---|
| 1 | 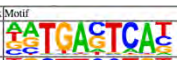 | Atf3(bZIP)/GBM-ATF3-ChIP-Seq(GSE33912)/Homer(0.994)<br>More Information \| Similar Motifs Found |
| 2 | 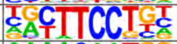 | Etv2(ETS)/ES-ER71-ChIP-Seq(GSE59402)/Homer(0.968)<br>More Information \| Similar Motifs Found |
| 3 | 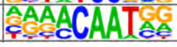 | Sox17(HMG)/Endoderm-Sox17-ChIP-Seq(GSE61475)/Homer(0.954)<br>More Information \| Similar Motifs Found |

## F

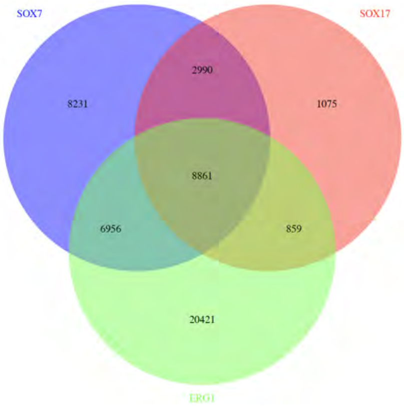

**Figure S11. A** UMAP visualisation of scRNA-seq data of CD31+ EC isolated from E12 *BmxCreERT2;Rosa*$^{tdTomato}$ lineage traced hearts with accompanying gene expression profiles. Raw data obtained from D'Amato et al (2022)[20]. **B** Chosen gene expression profiles from scRNA-seq data of ApjCreER lineage traced EC isolated from E14.5 hearts. Plots taken from publicly available Shinyapp visualisation of data from Su et al (2018)[21]. **C** UMAP plot and chosen gene expression profiles of CD31+ EC from *BmxCreERT2;Rosa*$^{tdTomato}$ lineage traced from E17.5 hearts. Raw data obtained from D'Amato et al (2022) [20] for reanalysis.

#### Figure S11

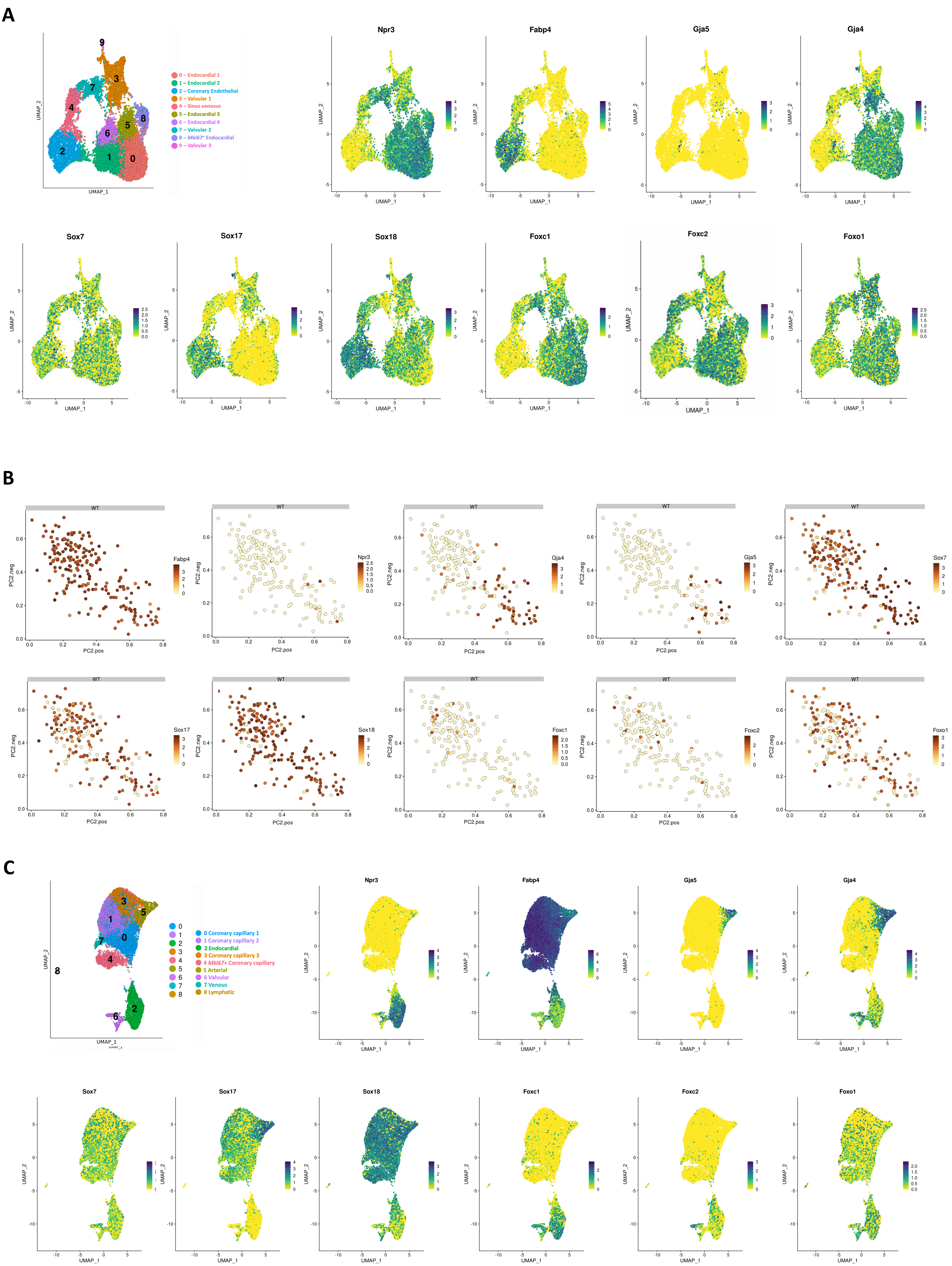

A
0 – Endocardial 1
1 – Endocardial 2
2 – Coronary Endothelial
3 – Valvular 1
4 – Sinus venosus
5 – Endocardial 3
6 – Endocardial 4
7 – Valvular 2
8 – Mki67+ Endocardial
9 – Valvular 3
Npr3
Fabp4
Gja5
Gja4
Sox7
Sox17
Sox18
Foxc1
Foxc2
Foxo1
UMAP_1
UMAP_2
B
WT
PC2.pos
PC2.neg
C
0 Coronary capillary 1
1 Coronary capillary 2
2 Endocardial
3 Coronary capillary 3
4 Mki67+ Coronary capillary
5 Arterial
6 Valvular
7 Venous
8 Lymphatic

**SUPPLEMENTAL METHODS**
**DNA sequences of the arterial lines, in the orientation used.**

**>Mouse Acvrl1+6**
GGTACCTACACAAACGTCACACAGTTGTCTCTGCTCCTGGTGGTCCCTTTAAACCCCCGTCCCGCTTCCAGCCTG
CCCTGAGCTCCCTGCTGTGCGTGACTTCAGCCTGTTCTATCCAGGCCTCAATCTAAACAATCTTGATTCCTGTTG
CCGGCCTGGCGGGACCCTGAATGGCAGGAAGTAAGGACAAGAGCCTGTTTATGTTTGAAGCAGCCAGGCTGGGGG
TGGGGAGTGGGGGCACTGGGAAACGGTGGGCAGGGGTGGAGGCTGGAGCGATGGGCAAACGGCTGAGGACAAGAG
ATGAGCTATGAGAGAGTCGTCCTTCCCACTGTAGCTCTGTCTGTCTGCAACCCTCCCGGCCTATTACCCTCTGAA
CAGTTGCAGTGGG

**>Mouse Acvrl1+16**
CTGCTGCCTTCCAGGCTAGTACCCCCCCGCAGTGGTCCAGTTGCAGGGGTCCAGAGAGAACCCTAGGGGGATGGG
AAGTAGAGCATCCCAAAGTACACACCACCAAATTTCCCTGGGACAGGCTTGCTCTAGAATGCCCCCATACCTAGA
ACCGTGCTGAGTGAGGTTCAAACTTCAGATCCCACCCGGGATAGACAGCAACCTGATCCATGACAGCCACGTCGC
CCTCTGCTGGCCAATGCGCTGCCCACGGCTGCCCTGCTCCCTTCCTACAGCGGGCTGGCTCCTCCCAGTAGGCCT
GGGCCCTCCTTCCCGTCCCTGCCGCAGCGTGCGTGCTTGCGTGCATGCGTTCAGCTGTTCTGGGCTGTAGGGACT
GTGCCTGGAAACCACACCTAGCAAAGGAGCTTTGTGCTTCCTTACTCAGAGGCGGCTCCGCCCCTCCCTGTTCAG
CTCCTTCTTAATTCCCTCCCTAGGTGCTCCCCACCCCTTGTCTGCAATTAGTCCACACCCCCGGAATGAAAAGCT
CCCCTTACCTGGAAATCTCAGAGAACACCTCATGTCCCCTTCCGCCCCACCCCAGAAACTATGAGCAAAGTGCCA
CTGGAG

**>Mouse Acvrl1+19**
CTTCTTCTCTGGGCTCCCTCCTGTAAACACTCAAGCCCGGGGCCCCTGCTGGAGTATACAGTACAGCTTTCGCTG
AATGCCAGAGCAATGCGTCTGGATTCTGGAAATCCCAAGACAGCTCAGTTAGGCTGATCAGCAGCAAGGCACTAC
CCAGAATTCCCAGTCTCTGGCATCCAATTCCCCTGGACCTGCCTCATCCCTTCAGGCAGGCGTTTCCTGTTGATA
AGAAGCTTAGTTACAAACACTGATAAAGAGATGAAAAAGGAAGTCGACAGGCTAGGGTATGGCGGGCATTAGGTC
TGCCTTTGCTACCCCTCTTCCTGCTTGATGCCACACACCCCTTCCCCCCACCCCCACCCCGCAACCATTTCCCAG
AACTGCGCAGAAAACAGGCCCCCACCTCCACTCCCTATTTTGGAGACTCCCGTTATTTGGCCCGAAGTCTTACAT
AGTTCTCAGAACCCTGGGAGAGGAGCTTTGGAAGCAGGAAGCACTGAGTGCCGGGTCCCCCCAACCCCAAGTCCA
GGGGCAAGGGCCGAGGCACAGAAAGTACTGCCCCACAGGTTGGGTTCCATCTGAGATCTTGTCTGCCCTTTTCTC
AGAGGCCTGGATCAAAGCTGTGCTTTAGTGGCTGCTGGCTGGTCTCTAATTAGTAGATTTTTTTCCCCATCATTT
TATTCTCGGGGTTACTGATGGGAAGTCTCGCTAATTTCCAGTTGACTAATTTGGAACCTGTGATCCTTCAGGACA
AATGCTTCAGGCAGATGCTTAGAATAAGAAGCAAAGCAAACAAGTAGGGTTTAGATTACAGAGGCTGCTTTCTAA
GCGCCCTGCCTGCAGTTACACCAGGTGGGTGAGGAGGCTCAGCCTGCTTCTGGGCTTCAAATTGTCCTTATTCAG
GGACAGAAGGTACCACGATTTGATGT

**>Human Cxcl12-184**
gaatttagtggggagaagcagagagaggcaatctaccccatcttCTAAGCAGAGCAGTAACAAGGTTAGACTGGT
GTTTTAAAAACGAGTGGTGATGGtctggagtgagtcaggtctgcattcaataccagcctttccacttactcaata
tgacctggcatattaccaaagcatccctaaatgtgtttcctatctgtaaactggggagaattggagggtcttcct
ggtaggatctttttgaggactgaatgacgtcacaggtctaaagcgctcattgctgtgtctgacCTGCCAGTTATT
GTTAACACATTTACAATTCTAGAACAATGTGGGAAGGACAGACTGAAGCAAGATAAGATTTAAAACAGAGAAATT
ATTAAGAAATTTGCAATTGCCCACGCAGAAATGAAGAATGCCTGAATTAGGCCAATGAAAACAGGAAATGTCCTC
CCTTAAATGGCATATGCTGAGAATACTGACTTGGCCAAGTCATATTTGCAGGAATCCTAGGCAATCAAGAATTCT
GAGCAGACATTCCTCCCCTGTGAAAAACACAGCACTGGGGACAAATGATCAGGAAATGCCTGGTCCCTGCTCACT
TGATGAGGAGCCACCCTGAGAGCACTGTGGGAGCGGCTGAGCTGGAATCCAGAACTAAGACTACGGAATAGCAAT
GTGGAGCTGCAGCAGCAGAGGTGGCCCTGCTTCAAGGCCCTGATTGTCCCCGACTCTGCAGCGGAGCTTGATGTT
GGAAGTCAGCCCGCCCTTCACTCCCAGGCTGGGCTAAGGCTCTTGAGGCTGCCAGGATTTCAGTAGAAAGCCTGG
GGCCAGATCCTTCGCAGAAGACTGAAGAAGACAGGACAGGGTGAATTTGCAGTGCCGAGTCTCCATTC

**>Mouse Cxcl12-2**
GGTGCCACAGTAGCTTCTGTTCTGGGGCCTCAGGGCGCCTGCGGGAGGGGCAGCGGCCAGGCCTGAATCCCTGGT
TCGCCCTCGACTCAGCCCACGCGGGCCTTTCAGGCTTCTGGGACAGATCCTAGGTCCAGCTGCCCACCTGATTTT
GTGGCAAAGAAAAAAAGAAGAAGAAGAAGCGAGAGGCGATGGCGTTTGCTTTGGCCAGATTTAAGGGCAAGCGAG
GCTGCGCGCGGCTCCCGCAGGGTCGGATCTCCGAGCTCCAGGGCGCCCCTCCACCCGGGTGTAGATTTCCCGCGG
ACCCCTTCGCCCTCCCGGGTTTCATCAGCTGCGCAGGAATGGAGCTGGCCAGAGCTCTGGGAGCGGGGAGGGAGG
CGCCGCCACCAGAGGGCGCCGGAGCCCCAGCGCTGCGGCCGGTGAGCGGCCAGGCTCCCCGGCCCAGCCCAGCAA
AGGCCTGGGGACGCCCGACGGCTGCCTTCTTCACTGGACCTCACTGCCTTCAGTTCTTTAAGAGCAGGGCCAAGT
CAGTGGGGTTCCCGGCTCCAAGCCCAGTGCCCAGGGTGGGTGGGTGGGTGGGTGGGTGGGTGGATGGATGGATGG

ATGGATGGATGGAGTGCCGGCCCACAGCCATCTAACGGCCAAAGTGGTTTTGGAAAAAAAATGCACAGAAGACAC
CTACTCCCACCAGCGGAGTTCCGGAGCCCTCGCAGCCTCCTGTTGACCGCTCCCGCCTAATGCAGCCGCTGACCG
CCCACTCCCCGACGGCCAGGACTCCCCAGGGACAGGGACGTGTCC

**>mouse Cxcl12+239**
TGCAATTCAGAGAAGCAGCAGACCTGTCCTCTCTAGTTTTTGAGCTGGCAAAGAAATAGTCATGGGGAGGGTTGT
CAAATGGTACTAGAGAACGCGAAGCAGGCGCCATGTTTTGAACAGCTGCAAATTCTCCTGCTTCCTGTATTCACC
AGCGAAAGGTAGGATTTCTTTTCCTACACTGTGGATCTTGGCTGGCCTAACACTTAAATCAACAATGCGGGGTAC
ACTTGGCAGGTGCCTAACTCTGTACCAGACCCTCTGCTCTCCACTGAAGGTTCCACTTTCATTTCTGGAGTCCTG
CAACCACTGTGTAGACAAGGCTAGGTCACCTGCTGGGTTACAGGAAGCCTGTGGCCCACTCTTCCCTGTTTATGC
CTCTGCCAAAATGAAAACAATCTCCAGACAAGTTAAAAGGGCCACCCAGACTGGAGGAACCCCCCTCTCAGCTCA
TCACAGACACAAGCCAGTCCAGCCTCCATCCACGCAGCCTGAACATCCATTGAACAACAGGCTTGTCCAGCTGCC
CACAGACAGCGAGCAAACACCCACAGTGTTTTATCA

**>Mouse Cxcl12+265**
AATCCTTCCTGGGACAGGATGATGGCCATGTGTGCAATGTGCCAGTAGCACTATCTGATGCATAGCCCTGGCCCC
TCTGCCCACCAGGACAGAGAAGAGAAATGAAGCCCTCCATGGCACACTGCAGATCACAAGGTTCACTCATGATTG
CTTCAAGCTGACCTGGGGCAGGATGGGCCTCGCTTAAGAAAAACAAACCAAGGGAGACTCTCCCAGCAATTGGGG
AGGAAGTTCAGTCAACGAGGGTTCTTGCACGACAGAGCATGGGCTGAACTGGACAGAGCGTCTGATTTAGAAAGT
ATTTTTATTCTATTACCTATTTTATTATAATGGTGACTTTACTCTCAGAGCCAGAAAAATAAACATCAGCTCTGC
AAGAGCCTTTGGTCTGAAAGAAGAGCATAAACATTCGTGACCTTCCTATTCCGAGTGGCGGGGACTGCTGCCTGA
ATAGGAGTCCCCTACAGAGCCTTCTGTGCTGGGAGGCTTCAAGAGTTGGCCTTTGTGTATTGGGAAGATGAGGTT
GGTGTGGAATATAAACACTCTTCTTCTATAGGTTTGAAAAGGTGAAAGGCTTGAGGCTTTTACAGCCACCTCCTC
ATATGGAGGTGTGTATTTCCTATTCTGTCGGTG

**>Mouse Cxcl12+269**
CAGGACATGTAATGACCTATTGCCTGTGTCTAGGGACGCATCACCCTTGGCCTTATGCAGTCAGTCAGCATTTCT
GAGGAAACCTGACCCTCGTTGTGGTCAGTCCCATCTCCTTCCCACACTGGAATTTTATTAGGCAAAGTTGCAGAG
TCTTTTTCTACAATTGTTTTCTTGATAAACACCCCGTGTACAGCATGTGGACTTTCTGTAGTTTCATTGCACCAG
AGTTTGCTCCTGAGAAAACACTGCTATAGGGGGAGCCCAAAAAACGAAGCTGGAAATAACTGGGGAGTGAATCAG
GAGCTGGTGTGACTCTTACTCCTTGAATTCCAGGAAAGTGGGGATGTGGCACAGCCCTGTGTGGAGCAGGCGCCT
GGCAGCCCGCCCAGCACACAATGGGCCTGTGATAGCATCATTGGCTCCCTTGGAGAGAGGACAATGGAAGCCAAC
AGCCACGCAGGCCCCAAGTCGGAGCCATCACAGTCAACAGGACAGGCGCAGGCACCCACTCCCTCACCCGGAACA
AGGATTCAAAATAGGACCTGGCCAGGAGGCCC

**>Mouse Cxcl12+298**
TGTGTGTGTGTGCATATACATGTGTGTATGCATGTACTTGTGTGTGCACTGATGGTGCATGTGTACTTGTGTGTG
TATGCACGTGCACACCTGTGTGTATATGCATGTGTATGTGTATACTGAGAGTGGAACCCAGCTGTCTGCACATAC
TAGGCAAGTACTGTGCCACTGAGCAATGTTCATCCCCAGAATATGTTTCTTTATACACATTAGTATCTTTTTTCA
TTCCAGGTATGATTTTATGAAAAACTATTTCAAATCTTGTATATATTATACTTCAAAATCTAGACAAGGAATATA
AAAGTCATGTCATCTCAGGCAATCCAGAACAACTTATGCCAGTATTTGAAAGCCCATTGGCTAGCAGAGCCTTCT
GTGGCTGCCTGCCTCTCTCACAGTATCCCCCTCCCTTCAGTATTCCCCCCTCTCATCCAGATATATACCTGTGTT
CACTAGGACAGGGAGGAGGTGAGCTGGCTAGCAGGCTGGACAGTGGCTTGGGTCATGGGTCTTCTCAGAGGCATA
GCATCTGACTTGGAATTCCCCTTCTTGAGGTCCTCCAGGGATGACATCATGCACATTCTGCATTGTGGTTCACTT
TCAGTGCACAAATGCCAACTGTATGCACAGGAAACCGAGGAAGGAAAGTGAACGTGAGAATACAATAGCTGTCTT
TAAAAATTACTCATATTCTAAACAAAAGTATAGCAGGCTGTTTCAAATTTCTGCTTATTAGGAAAGTGCCTTGAA
GCATCCTTGATAGACTTGTGACAGGGCTGGGATGCTGTTGGTGCCTGTGGGGCAGGGGACATTTGCTGAAATGAG
CCCCTTAAAGCACTGTCTACACAGGGAAGTGAAGAAGACCGCTGTGGTGGTTGAGGTGATGTCTCCTTGGATAAT
AATGTCAACCTAGGCTGTCACTGAACTTGGATGGGTGGGTTTAGAAAGTGGGAGAACCTTGCCTAGGTCCCATGT
CTTCTAAACACTGCAGCAAGGAGAGGGGAAAGACATGAGTAGGGGGCCTGGGTCAAAGTATTCAGTTGTCCCTAT
ATTGACCTCGCCAAGATGGGCTTCACGCAA

**>mouse Cxcl12+376**
GCCATGAAGTGATAGCAGAGCCAGTCATCACAAAGGACACCCCTTTCTGTGGGTAGGCGCTATAGGAAAGGCATA
GTTCCTGGCCACTGCTTCTCTGTGTGCTTCCATTGTACTCTGTTCTGAAACGTCAGCAGATGCCCCTAACTACCT
CTTCTTGCCTGGGTCTAATTTGCTTCAGATTTGTAGATTGATCTCAATGTTTTCTGGAAACCAAGTCCACCAACC
AAATAGCTCTGCACACAGAACTATCCTTCCCACGTGACTTTCCTCAGATTGTCAGGCAACGTGACAACCGTCACA
GGTGGCGCTTAATTCCAGGATCATCATACATCTGATTGGAAAGAGACACAAAGAACAGCAGTCACCAAGTAAATA
TTCCTTGAATAAACGAATATATACTTAGCAATTTTAGGTATTTCCAGGCAAAACTGACTCATTTGTGAAGAGCCT
TTGGCACCCAGTTTAGCAGCTCCAATAATTAGTCTTCCTGTCCAGGGCTCACATTTCTCTGAGGGGCTTAGATAG
GCAGCCATTTCCCTCTGAGTAACATCCAGGACCCAGGACAGACAAAGCTACTCTGACTCACAGCACACAAGAGGG

CAAATGATGTCCACATTCTTGTGTGATTACAGGGTGCCTGGCCATGGACCTAGGTGTCAACGTGCCAGGCCAAAC
CAGGATGCTGACTGACCCAGATAGGCATCGCTGAAATAGCAGCCTGGAATTTACCATATTTATCATGGCTTCTCA
GTTTAGGTTTGTGAGATAGGATCTTTCTGGATATCCACGACTCATGTCCAACACATGGTTCTCCTACCTCTAACT
CCCATGTGCAGCTGTGGTGAATAATTATTCTCTAAACTTGAACGCAGATGTATCTCTTGAGCACCCTGTCTTCAG
TTACTTTGGCCAGGCCTACTGTCAGAA

**>mouse Cxcl12+383**
ACCAGCAATCCCATGCTGTGTATGTGTGGTTAAATGTGTGAAGCCAGAGAAAGGCTTGAATTCCCTCACAAACCC
GGGCTCCTCCCATCCTTCTTTTCATGGCTAATTTAGTTAAGTGTTTAAACAATGTAAACCTGGCAGCTCCCGTCT
ATTGAAGCTGAGATGGAGCACAGCTGCCGCCCAGCCCCAGCCCTGCAGTGCAGCCTGGGCTGCAGCACCCATCAG
CTTTCCTGAAATAGCAAGGCTTGCTTCCTTCCAGGCTTCCCCTGGAATAACCCGGGGAGGAGGCTGCGTTTCCTC
TGCCCAGGCCACTCTATTCATTACCACTCAGGGAGTCCATATTTGGAGAAAAAAGTTTCACTGCTGTGTGGA

**>Mouse Cxcl12+439**
CACTCTCTTCCTTCCTCCTCCTCTTCATCTTCTTCCTCCTCCTTCTCCTCCTCTTCCTCATCATCCTTCTCCTTC
TCCACCACCTCCTCATCTTGCTCCTCATCACTCTCCTCTTTCCTCCTACCCTCCACCTTCCTCCTTCTCCTCCTA
CTACTCCTCCATTTTCTTCATTATCATCTTTCTCTTCTCATTTGTAAGTCTGAGCTCACAGACACATCAGCCGTA
GGTGGGCTTGGTGGGTGAGCATGCCCTCACCTGGTCTGATGGATCTGGTTGGGATTTGTTTGCTACAGGAAGGCC
ACTGTGTGCAGGGCCTTGTTTAACCCGACTTTCTCTCTTTAGAAAGAGAATTCTCATCAGATGAATCATTCCTGA
CATGCTCGCTGGAGCCAGAGACTTTTGGCTCCCAAGCTGTTTACTGCGGCACTGACTTTGTCCCATAAGTGTTCT
TAGGAAACAACAAGGGTCAGAATTGTCATTGTTTCGGCTTTAAAAAGCTGGAGACTTGGAGATAAGCCAGGGCTG
GGTAGAAGGGTGTTCCTACCCATGGGACTACTCAGTGTCACACAGATCAACACCTCCTAGACTCCTGAAGCAACT
TGCCTCTGTTGCCGAGGCCTGGCTGCCTATGTATCAGATAAGACACTTGCTCTCTGATTGTCTGAGAAGCCCTTG
TTACTGTCCAACAAGGCCAGGCTGCTCATTTCCACTCACAGGTGTGATGTAACCAGTATGTAGGG

**>Mouse Cxcl12+445**
GAACCATATGCAATCCTTATCATTTAGGGAAAATATTCCATAAAAAATACTAGATTTCATTACGGCCTGTTTGGT
GCGAGCGGGCTGCAGGTTTGGTTGCCACAGAAATCGCATCATTAGGATACTACAAGTCTGCTTTGAGTTACCCTA
GCTGCAAAGTCAGCCAGCCCCGTGATCCCATCCCCTGCATCTGTGTGCTTACCCGCTGTGATTCATTACAATAAC
AGGAATTTGCTGCTGTTATGATTGAAATAAATCAGCTTTGTGGTCCGGGCAGCCCGTTGTGATAACCATGGCTCC
ACCTCAAATTTGCTGCCTGGTCTTAAGTCATCTGCTGTACAGTCTAGCCAAAGACACAAACAAACAGATACTATA
TATGTTAGGAATTTATATATAAACTACCGAGAGCCAGGAG

**>Mouse Cxcr4-232**
TTGAAGAATCAGAGAGCAGAAATGGAAGGTGCATCTAGCCTGTCAGAAGACTCTGCCGAGGGAGCAGGTAGAATT
GGATCCTTTCTGGCAGGTTGTTCCACAGACATCATTAGCTACAGAGGTTGCCCGGCCACTAGATGGCAGTCCTCA
ATTGACGTTCGAGTAACTTGGGTCGTTGGCCTCTAAACCCACTGATATCCACAAGGACAATAAAAAGCAGATGAT
GAATTTGATGGACTAAAATAGGCTAGCTTCTCTCCCGGGAAACTTGGCATTAAACAAATTCATTTAGCAAATGCA
GAGCCTAGAACAATGACCCAAAACGGGGCTGCAAGCATTCCTTTTGGTCTCTAAAGGCAGGACCAAAGTTCCCAC
ACTTAAGTTAGGAAGCACGGAGCCAACTGGTCGTATGGGAAACAATGCAGCGGAAAAGTAAATCTGTGGACCCCC
AAAACATGAGGATAAGACATCTGGCTGTTATAATGCAAGTGATAGTAACCTGAAATGAGATATTATTAAAAGATG
TATTGAAGTTTTCGAGCC

**>Mouse Cxcr4-194**
GGTGGGGAATGGCAGAGAGAAGAAAATCTTGGGAAAGGCAAAAGAGCATTGCTTATCTTGGTAACTTTGGCAAAG
TGTCTTTGATAGTCAAAACCTTGAGGCAGGCACAGTCTTACATCATCATATGACATCATTTCATGTAGCCATTTG
TCCATACACTCTACTGATTCCTTCCCTTCACCAACTCATAGAATGTGTGCCAAAGGAACCTGAGAGGTGCTTTTT
GGTTTAGAAGTGTTTACTATTGTGTCCATGTATAGACAAAACTAAGACCTGGCTTTCCTCAAGGGTTGTACAGGG
TGAAAGCCTTTTACCTCCCCCCCCCACACACACAGCTGGAAGAGCTACATTCAATGAACTTATAAATAGCCTTTG
ACACCACTGGCACATAGAACTTGTCTTTTCCTGCCCCTCCTTCAAGTCTTTCACTGAACCCAGCAATGCTCTACT
AAAGATCAGGAAAAGGAAACATATTTCAAGCCTCCTACCATTGTGCTATTTGTTGACAGTAGAAAATTTAGTAAA
AAGGTTAAAGGGCTGGCTACAATGGCTTTAAGAGTCAAGTGTAAATTTTCTCTTGAGCAGGGACCATCAAGGAGC
TAGGGGGTATAGTTCAGTTAACAAGGTACTTGTCTGGCATGAACAAAACCCTGGATTCCATCTCTACTATACTGG

**>Mouse Cxcr4-130**
ATCAGGGCCCCTGGCCAGCCTTGTTGGAGGACCCAGGGAAAATAGAGGAAGCTGAGCTTGGAACCTGCTCTCTGG
TGTTCCTCTTTGGAAAGGCTGGCATGTAAGGGGACTTGCCAGGATAAGCCTGCAGAGAGGAGGAAGCCAGACTCT
CAGGGTAAAATTCTGTTGAAGTACGGTAAACAGCCCAGGACTGTTGCTTTCCCAGACCTTACTTCCCTAAACGAG
GGCATGTGCTACAAAGGCCTGGTGATAGAACCCTGGGATCTGTAAGGCCAGGCTAACAGAAAGGCAAAATCTGAG
CCAGTGACTCATGGGGACCTTCCAAAGTTCTCTTTCTCCTAGGACTTTTGAAAAGCCTCCTAGAAGCAGATATCA
CATGTCAGACTCCACTCTTGTCTCTGTGCCAAAACA

**>Mouse Cxcr4-117**
ACTGAACATGTGGGCGGTCTATGAAGAGAGATCATAGATGCTGCCATTATTATATTTCCTCATACACTTACTATG
TGCTCAGGGGTCTCAAACTCAAGGTCTCATTTTGTCAATACTGTCATCTTAAGAGACGGTTGGTCATTCCTAATG
TAAAGGATGAGAAGATAGGCTCAGAGAGGCGAAGCTAGATGTCTGTAGTCACACAGAGAGTAAACACAGAAAAAC
AGATTTGAGAGCTCCATGGAGGAACCTTGCCCCTTAGCTTACCCTGGGGGTCCCTCCCTTCCATTTCAGCTTGTT
ATTTCCTAAGTTTGTTCTCTGTCAGCAAAGACATCCAGAGGCTTGCTGTTTTAGCCTTGGTGACCTCTGTTTATT
AACGTTTGTCTTTCCCACCAGGTTTCAGCATCCTCAAGTCCCCAGCACCTTGCCCCAGAAAAACCGTGCTCACTC
AGCATGTATAAACTTAGTACCCAATGAATGGAAGGCGGTGACTATGGGGATTTCAGGGTTCAGGGCCATCCCACA
TAGCCAGCCTGTCTTTTCTGAGGTACTTTGCAGAGACAGGCTGTCTGAGACCCCTGAGATGTCTGTGGCTGGTTA
TTTCCACTTACCTTGAGTCATCAACACTCCCCTGGTAGGATGACCTGGGGGCCCAGAGGCAAGCATGGAGTCCCA
CATCCTGCCCCCTTCCTGCATATGCATCACTTGGTACTAACATGCAGAGAGAGTCTACACCACATCAGCCTCTTG
TTCTTCTGTCCATACCACAGAGCAGAGCCAACACATGAGTTCTTAATCATGTATTCAAACATGCTCATCCATCCA
TTTGTCCATCTGTCCATCCATTC

**>Human Cxcr4-117**
CTAAGGACAGACAGTTCCTTCAGGTTGTGTTATACATATGAGAAGTGTATGAAAAGGGCTCAGTGAGTGGTTGTT
GTTGTTCCATTTTATTGAGTACGTATTGTGTGCTCATGTTCTCACACCTGATTTCTTGTGTTGTCAACACAATAG
TCTTGAGAGGTAGGAATTATTATCATTCTTATTCTATGGGATAGGAAACTGCGGCTCTGGGAGGTCAAGTCTGAG
GTCACATAGCTAGGAAGTGGTTGAAACTGCACTGAAATCTCAGTGTGGGAATCCTTCCACATCACTCTCCAACTG
AACCCTATACATTTTTACCCCAGCCCTAGGGTTCCTACCTTCTCTTTAGCTTTCGGCATTTCCACGATTCACTCT
CCTCTGGGTGTGCCACCCTGTCCTCAGCCAGGATGCCCAGAGGCTTCCTGTCCTGGCCTCTGCCATCTCTGTCTC
TCTCCATGCTTGTCTCCCCCACCAGACTTCGGCATCCTTGAGGCCTGTGGCTAGGCCAGTCCCTACTTACCCAGC
GCTGGTGGCCAGCCTGGCCCACAGCAGACCTCTCCCCAGTGTGTGTAAAATTAGAAACTGACAATGCAAGGTAGT
GCCTGGGAGTTTCAGGGTGCAGGGCCATCCCACACTGTCTGTCCACCTCCATCTCTGGGTGACCTGGCACAGATG
GGCTGTGGGACGTTCTGAGATGTTCCGAGATGGCTGTGGACTGTGAACTCCACTTTCCTCTGAGTCATCAGCACT
CCACATACACTGAGCTCATCTGACTCTACTGGCAAGATGGCCCAGGCCTGCTGGCTTCGGGACCCACATCCTGCC
CCCATCCTGTGTACCCATCGTGTGGTATTAGTAAGCCCAGCAACTCTGGGCCACACTGCTTTACCAGTCCTCTAA
GCCTCTACTGCTGACCACAGCCACATGCAAGCTTCCAAACATGCCTCAAACATGTATTTGTCTGTTCATTCATTC
ATTCATTCA

**>Mouse Cxcr4-113**
CCATGGAGGAACCTTGCCCCTTAGCTTACCCTGGGGGTCCCTCCCTTCCATTTCAGCTTGTTATTTCCTAAGTTT
GTTCTCTGTCAGCAAAGACATCCAGAGGCTTGCTGTTTTAGCCTTGGTGACCTCTGTTTATTAACGTTTGTCTTT
CCCACCAGGTTTCAGCATCCTCAAGTCCCCAGCACCTTGCCCCAGAAAAACCGTGCTCACTCAGCATGTATAAAC
TTAGTACCCAATGAATGGAAGGCGGTGACTATGGGGATTTCAGGGTTCAGGGCCATCCCACATAGCCAGCCTGTC
TTTTCTGAGGTACTTTGCAGAGACAGGCTGTCTGAGACCCCTGAGATGTCTGTGGCTGGTTATTTCCACTTACCT
TGAGTCATCAACACTCCCCTGGTAGGATGACCTGGGGGCCCAGAGGCAAGCATGGAGTCCCACATCCTGCCCCCT
TCCTGCATATGCATCACTTGGTACTAACATGCAGAGAGAGTCTACACCACATCAGCCTCTTGTTCTTCTGTCCAT
ACCACAGAGCAGAGCCAACACATGAGTTCTTAATCATGTATTCAAACATGCTCATCCATCCATTTGTCCATCTGT
CCATCCATTCACTCATC

**>Mouse Cxcr4-109**
AGAGTGTTGGTAATTCTTTGGTATGCTGTAGTTTAAATTATACTTAACTAACCCCTCCACCCTCTACCCCCAACT
AGGAGAAATGCATTACCGCTGGCCTTCTTGGTGGAAAATTGTATTCCATGACAAAAGACAACATCCTCTGACTTG
GCTTCCACCTGAAGTGAGTGCCTGCAGGATGCATGTGACTCCTTTTAATTGATAACAATGGCAAACCGAGCAGGC
CCTCTGGGGACATGACTCATGGTGAGTCAGGTGCACATGAAGGCTCGGCCCAGCTGTGGGTTTTTTATTTGGTTG
TTGTTGTTGTTTTGTCAAGGGGTCTCTACTGTGCTGGAGCTTAAATGTTGGTAGGATAGAACTCAACCAAGTGGA
AAGAGAGACACATTCATTTAGTGACTGCACCCCTGTTGAGTACCAGTGATGCCAC

**>Mouse Cxcr4+1**
TTGTGCCGGGTCGCCGGAGGGATGAACAGGTGCAAGCTCGGAGACTGTTTGACTGAGCCCCAGTGTTTAACCGTC
GGTGGTTTCCTGAGCCTTTTGAAACCCCCTCGCTTAGGGAGGGTTTTGTTCTTTGGTGATTTTTTTTTTTTTTTC
CGTTTCCTCCGGCGGCGCGCAATTCAAAGGCGCTCGCGCTGGAGCAGCCCAATCTGTCGCTCGCACCCGGAGGAA

**>Mouse Cxcr4+135**
GTTTTCCCTTCTGGCCACCGCAGCATTAACACAATTGTTAACCCTGGCATGCCTCGTGGTCCCGGGAGCCCGCCG
CCTCCCCCTTCCTCGGGACTCTGGGAGCTTCCTCTGCTCGCTGTGTGTGTTCTAAGAGTTTGCCTGCACTCTGCC
TGTTTATAGCACAGGAGCAGGAAGAAACAAAGCAATTGTGAGGCTTGTTGAGAAGCTGTTCGTATGATATTAAAG
GTGACCTTGCCACTTATGTTTCTATTTATTCTACTTCCTTCCAAGCCGGGGCAAATCCTTCCCTTCCGAGTTGGA

GCGGCGCTCTTTGGATAGCAGTCTGTGGGAACAGAGAGGCGTTCAGGCCCTCAGAGGTTATTCATTACAGTATCT
GGAGTATAGCGTTTCCTATTCACTCCCGGTCCGGTCTGTTAAGTGACTGTTGC

**>Mouse Cxcr4+151**
TAGAAGGCAGCATGGGGGAGGGTGGTGTTGAATAAAACTGGAAGCACCTCCGCACAGCCCTCCTCCATACCACAG
GATAGATCCTTGTTTGTTATCATAAGCACGTGTTTCCCCAAAATTTCCTGTTTTTGAGATGATTTGTGAATTGAA
AATTCTGTCCGTGGAAGGAAAGCCGTGTTATTGAAGCCTTACACAATGGAATGACTCCGGCTTTAATCCTTTGAA
TAAGGCTTTGTTTCCCATTGACTGAAAACTTCCTCCGGGAAGGAAAGCTGGAAATACTCAGGAAGGATTTGTATT
TGATTTCACAACAGCTTTGTGAGGAGGGGGAAAGTCATGGAAGCTTTAGTCTCCCATTCCTGGTGGACCGGCATC
AACAACACGCAGTTCACCGCTGTGGATGGCGGCGGCTCCC

**>Mouse Efnb2-333**
TATTTTAAATTCGAGACTTTATGTTCATTTTATTATACACAGTCTTAATTTGCGCCTCAAAAGAGGGGGCTCCAG
GGCGGACATAAAATTAACTTCTCTTTCAGTGTGCTAACCTTTTGGTTTTAATTTATTCACTTGTCAATCTTATGA
ATGCCTGGTGAACTCAGGCTATTGATTTCCGTGGCCAATGCTATTTTTCCTGTTTATTATAATATATTGTCCCAT
TGTTACCAGCGCTTGCACTCAGCTAGATTTCCCACACAGTGGACAGGAAACCGTTCCATGGCATCAGACTTCCTT
AGATTGATTTTCCAGAGGCTGGGAGGGACCTGCCCACTCCTAGCTCTTTGTGCTGGGACAGCCCAAGGGCCCCAG
CAGGCCTCCTTGGACACTTCCTCCCAGGCCCCAGAGTGGGGGTGGCCTGTCGCCCCTCCCGAGACAGTCCCCGCA
GCCCTGGGAAACGCTCCTTATCAGTGCCAAGGTTATGTCATTTCTTTTTACATACGATGTTTATTGTTTTTAGAG
CCAGCACTGTGGGAATGTTCTTTTCAAGCGCGACAATAGGGCAGCCACAGAGGCAAACTGGTGAATCCCCAGCCC
GGAGTTGCGGGCCGTCCATGCAGCTGGGTAGGGTGGCTCTGCGCAGGCTCCTGATTAATTGATTCAAAGATTTAG
CACCAGTCTTTAACCGGGATCTTGCCTGTGGGTTGCTTAAAACACCAAGAACGCTTTTAA

**>mouse Efnb2-141**
AAGGTTAGGTGCTTGGGTACAGCCCGACAGCCTTACTGACACATATCAAATGCAGGGTGAGGTGCCCAGGCACAT
CCTCAGGGGGTTTCTGTACCCATGGTGCAGGTGTAAGGACAGAGAGCTAGAGCAGGACTCAGGAACTTCACAGGA
CTCTCTCCCCCATCCCCCACATCTTTCCTCCATCTCATTCAGTGAGTGGTCACATTGTTAAGATGGCCTGAGAGG
CAGCCAGGAAGAGGCAGGAAGCAGAGACTGGCTAACAGGCTGCCTTTTTTGGTGCACTGGGAGCCCAGTGCCAGA
AAGTCTCATGAGAACTTCTAGAAAGGAAACACACTCTGTCTGTCACCAACAGAGGGAATAATGTCACCGGCCCCG
TAAGCACTGACTGGAACAATGAGAGCTTCCTGGTTCCAGGAGTCAGAACAAATGTGTGAAATGGGATGGAGGGGC
ATTGCCACCCGAAGCTCTCACAGGAAATTGCTCAGTCAATGGAGGGCTCCGTGTCAATATTTGGGTAAGGAAGCA
TGTCTCAGTCGGATTCCAAGGTGCCCCACAAGCAAGAATTCCTAAGAGACATTATGCTGTACTTGGGTTCTGGGC
TGCCCATTCTCATACCAATGTCATGTGTGTATCTACAGTGGTCTCTATCTGTAAGCTCAGGGGTTACATTTCTCT
AGATCTTATCTGAAGGAAGCTGGCCGCTTGGCCCATACCATGCGGTACAGCTGCAAAGTTCTCAACT

**>Mouse Efnb2-112**
TACATTTATTTATGCCCCGTAAAGGTACAGGGAAAGCCATACAGTGGTCTTTTTAAAAAGGTGATAACACGTTCT
TTACACTAAACCGTAACACCTACGCACACAGTGATTGAGTTATAGTCAGAAAGGCACCATAAATGACATTTCCTC
CCACTAATTTCACATTGCCTGTGCAGCTCTGGGGAGTAACATTACTCTTCAACAGAAAAATGTCCTATCCCACCC
TGGCAACGCCTCACAGCACTTGTATTCCCGTCAGTTTTTTTTTTTTATCAGCTTGACTTGCCCTGTTTCTTCAGG
GGTAACTTGTTCTCTGATGCTAGTCTGCATTTTATGGAGTGTGGAGTCATAGCAGTTTCCTTGTCCTGGTCCCAT
TAGCTTCCTGGTCCTATTGTTTGGAGTCTCCAGATCCTGATGTGTAATCCAACTGTTTCCCCACTATGTGGGAAG
CGGAAAGCTGCCGGCCTATGTCTTGAACAACCCAAGCTAGAGCTTTCCTTTCCCAGGAAGATGCTGGCAAGACTT
TCTGGGATGCAGCGCACACAGCAGCTGTGCACAGGGAGTGTGTCTGGTGGGTCCCGTCACTGGGTTGGAATCTGC
TGGCAGCATGATTTCTGAGTTAAGTCTAAAACTCTGAAAGGAAGATCTCAACAGTATTTGGGTCTGTTTCTTGGT
GGGTTTCAACTTTTTGCACCGGGTTATGTTTTTTATTCTGCTGTTTTCAAGTTCTTGATTCTACACC

**>Mouse Efnb2+3**
CAGCAACATAATTGACCAAACACGTTTCCCTTCGAAAGGGCAAAAAAATCTATAGTTGTAAATGACTCAAAACAT
TTTTTTTTTTTTTGCCTGCTGCCAGGAGGCCCTTGTTTTAAACTTACTGCGGCCCCCCTCTAACGTAGTCTTGAT
CTGTTAAAGAATTTTTTCGTTTTGTTTTGTTTTTTAAAGCAAGGTGTTGCATTTGCTCTCCTGGCGTGGTATCTT
TGCATCCACACGGGATCTACAGGCACCTGGTGGTGCCCGAAGGCGCCTGGCTGGGATCTGAGCGCTGCAGCCTTT
GCCATTCGGACTGGGGCCCTGGTGGTAGCTGCTGCTGACGCTCGCGGTGCCTACAGAGGTCACCCTTCCCTGCCG
AATGCTAGCTATGAGCGCGCCCATCCCCCCATCCCTTCCCTGGGCTGCCTCCCTCTCTCCACTTTGCACTTTCAC
TCACAGCCTCTTTTGGGGAGGCCGAGTCTCCCCCCCCCCCCCCCACTGCCTAGATTTAATATTTATATTTAAGTCT
AATTTAAAGAAATTCCCATTAAACCCCTTCTGAGTAACATAACCATCTTTTCTGTTCATAGTGACCACTATACTT
GGGTGTGCAGACGGAAGGATTAGAAATACTGATTTGTTTAAACAGCACTGCCACCAAGTGCACGTAGCACTTGGA
GAATCATGTCTTTGATTTGAAACCTGTTATGTTAGGGAGGGATTTGATCTTCAACAGGATAGTTTTCCGAGCTGT
GGCCACAGATGTATAGAGAGCTGCTAGGGCCCTGGGTTTTGG

**>Mouse Efnb2+37**

GCCCAGATGGGAAAGTGAGTGTGCTAACTTCTGTCCAGCTAGTGTTTCCTCCAGCAGAAAGCCTTGCTGAAAAGG
CTCATGGGGGTTGCTGATAACACGTGCTTGCTGGCATTTACAGAGTGAGAAGGCAGACTCAGAAGACCAGAGTTG
TAGGATGCTCCTTCAGTACATTCCAGGCTGGAGAAAAGAGCAAGGGCGTATGAGAAGGCTTCCCTGCCAGTGTTT
GCAGAACTTGGACGATTGACAAGCAGAGCATCAAGTTAATTGTTCTTGAAGCAGGAAGTCTGGAGTGGAGGAAGC
AGGTGTTGGGAGATGATTGCCGAGTTTTGAAATTACCGGGTTAGCTCCTTTGTGCTTTCTGCTTTTGGACCCCTC
TCAGTCCGTAGGCATTTGGAGGATGCTGTCCCTTTTACCTGTCGAGGAATCCTAAATCGGATGTGCATGGGAAAG
GGAGGGTTTAAGCCCATTTAGAAGCAGGTAGCATACGTGCAGCAATGTTTA

**>Mouse Efnb2+172**
GCCCATAGATGCACAAGCACCCAGCATCCAGAGGGTAAGGTAGTGTGTTCCTTGATGAGCGCTCTGGGAACAAGC
CCAGCCTGGAAAGGGTTGAGAGCATAGGGCTGCCAGGCCAGCTCACACTCCCGAGCCAGTGAGCCTGTCCAACTA
ATCGGCCGGGTTTGTTTGCACAGCTTTGCTGGAGCGTTTGCAGGGATGCACCCCCAGCTCCCCGCAGAGCCTGAG
AGAGAGTGGCCAGCCAAGATACCAGCGGGACCAGGATTCACCTGCATGCCCAGATACGGGCAGAAAAAACATAGC
ACAGTTTGTCCAAACAGCCCTGGGCGCTGCAGGATGGCTGTCCCCGAGGTCTTCCCCCAGCTTTGCTGCCAGGGG
ACACCAGCCAGGTTTATCTAAGGGCTGAGATACACTCTTTGAAATTCGCAGTCAGGCAGGCCTGCTGTGACCCAG
GGAGGGACTTCTAGAGCTCCCCCGGGGAGCCAAGCTCTACCTGGAAACGCTTGGCAGACACATTCAATGAGGTTC
CCACAGCCCACAGCCTTAGTCAGCACCTGGTCGGTGGAGTGACTCAGAAATTGGGAGCGAGGAGGGGCTTCAGGT
TTTCTGAACCCACTGTCACGTACGTCACGTGATTAACCAAAGCATGAGCCTGCTCAGAACCACAGGCAGGAGCCA
TCTGCCTTTGTCTTGCCTTTGTTATCTTACAGAACCAGCACGGGGAGGAACAGCCTTTGACTTTCCAGATTCCCC
CACCACCATCACCAACCCCCACCTCCAGCAGCAGGCTGGGTGATAACGTTTTTGATTAAGCTCTTGCACACATCA
GCAAAGCCTAATTATGAAAGATCTAATAATATATCGCCGTGTTTCTCTGACAACCACGGGCGTCTCCTCTCAGCA
TCCTCCGAGCTATTGGAAAGGGGAGCAGACACTGACACCTCGCACTAACTGCACAGGGAGAGGG

**>Mouse Efnb2+209**
CCTAAATACCATGATTTACTATAGCGCTTATGATTTCCATGCTGTAGGATTTCCCATCATGGCTGGTTTCAGGCT
AATGAGGGTTTAACCTGCTTGTGAAATAGTTTGCTAGTTGGCTGACTCCGTGTGCTGGTCTGAGTCAGCTATAGC
GAGTGCCGCCCCCACAGTGCCCTGCTGCCTGACCCCACGATACAGAAGCATGAAGAGCTTTGATTTTCATCACTT
CCTGTCCCACAACACGAAACAAACAGATTCAGCATCTTCGTAAGCCCCGCCTCTCACACTGCAAGAGGAAGGAAG
TTCAGGCTCTAGGAAGGCAGAGCAGCCAGAGGCTCCCCTGTAGCTGATGCAAGAACCGAGGAACCAGCATAGGAC
CAAGACAGGGACTCTCCCTGGGTTTAGTAGCCGGAAAGAGAGGAGTGAACGGGACAACAAGCTTGTAGCAAAATT
GGGATGGAAACCGAGCAAATGGTGGGTAGGTGTAGGAAGGCAGACCGATCTTTCCAGTGTGTGTCTCCAATCTTC
AA

**>Mouse Gja4+24**
GACACGGACCGGTAGGTGACTGCTGGGTTCCCAACCCTGTGCAGATTGGAGCCCTACAGTACCAGGTCCTCTCTA
GTCCCCTCCCCCAGCCCTGCTGCTACCCAGGGCTGCAGAACCCCAGACCAGACTTGCTTCCTGAGGTTGCTGTTT
TCCTTGGCCCCTGAGATGGGAGACCGTGTGCCCGACGTGGTGGAAAACAAGACATGGGATGGGGGAGGCAAGGGG
AGAGCAGTCCCTGCTCTCACCCCTACAACCTGTCTCTAGACGGTGGAAACAGAGAGCAGAGGGGCTATGCTTCCA
TCCAACCACCAGCCCATCCAAGCCATTGTCCCCTTGTTGTGACAGAGGGTTTGTCTGAGGCCCGTGGACAGGACT
AAGTGTTTCCTCTCAGCTCTCCCTGCTTGGACAGCTGGAGGAAACATGCAGCCAGGCAGCCAGCACTCCATTCCA
ACAATAACAGGCTGCCTGCCCCACTGGTAGGCCCACTGTTGGGGGGGTGTTGGGGGGGGAGCCTGGAGCCAGAGA
GGCCACCCCACTCCACCCTGTCACTGACAAAACATGTTTTCTTGTTTTCTGAGGGGACAGGGGTGGCTGGTACCC
ATCCTGATCAAATATGGTGCTGATAAGCGCAAACGGGCCAGGCCGCCAGGCCAGCAGCTAGGGCTACATTCCTGG
ACTGAGTCCGGTTGACCCTGGGGGTCACTCAGACCAGCCTGCCAGGTCTGGGAGTCTGGGCAGTGTAACAGAGAG
TTCAAACCCAAGTTCTCTTTTCACCGAAAGGAAGACC

**>Mouse Gja4+50**
GAGTCCTGAGACGCCTTTCCCAGCAGTAAAAGCCCCGAGATCCTGGAAAGACGGGGTGTGGCTGGTGAAACTGAG
GAAGGCTGAGGTTGTGAGAATCTTTAAAAATAGGTCGTTTTGCTCTGACTGGACTTGGGCGGGGGTTGGCTGAAG
GCAGAGGAGCTGACTGTCAGAGGCCAGGAGGAATTCACTCTCCAATCACAAGGCAAGTGAGACTGGTGTTGTCCT
CAGAATACTGTAGGAGCCCGGAAGGGGTAATGGTGAGCCTAGCCTCTGCCCCTGTCCCCACCCCCACCCCGGGTT
GCCTGACTGTATTTTCGGAGACAGAGCAGGAAGTGGGCCTTCACAATGTCAGGATACAGGAAGCTATCGGACTAT
TGAAGGCCACTTCCCGTGAGCTGGTCATTCACATACATTTGTCACCGGCTATCAGGCCTTTCGGGCCCTACCATT
GTCTACGGGGGCTGCTCAAGCCTGTGGGCCGCCAGCAGAGGCCCTTGGACACAGAATACACTATTGTGCTAACTG
GCAGCACCACACAGGGTGGGAAGGAAAGTGGTGGGTAAGAAAGGGGACGCCAGCTTTGCATTGTCATCTGTCGCA
CAATGGGGCAGGGGGCTGGGGTCCTTTCTTGTTCGTGGGAGAGTGAACTTCAGGAAGGAGGGATGGCCGCCCAGG
AGTCTGACTTCAAGTTCAGTACCAGATGGCTGTAAAGGTGCC

**>Mouse Gja4+57**
CCACAGTCTACCCTTGTTGCTGAGAATCCTCCTGCTTTCCTAGCCTCACCTGTCAGCCTGCCCCACCTCTGGGGA
GACTCAGCTAGAGGGAAGAAGGTGCTAAGGAGAGAATGAAAGAAGAATAGAGGCATTCCAGGCCAGTTCCTGTAG

GCTCTTCCAAAGAACATCCCTCAGGGGAGGACTTCCTTAGTCCAGAGGCTGCTGGGACCTGCTGAGTCTGTGAGC
ACCACCTGGCAAGTACATTAAGTGGTTCCCTGGGCTGCAGCCCAAGTCCTTGCTGGGTTCCTTGGAGTTTCCCTG
TCCACCCCGCTAAGCTGCTGCGGGGCTTTGATCACCACTACACGAAGGGTTTCCTGAATGCTTTCCTGAGGACAT
TGTTCCCCACTGCAGAGTCATCAGGGGTCAAATTCATCATTGGCTGGGCCGGGCTGGTCCAGGTCCTACCCGGTG
GGGAAACACAACCCTCTGCCCAAGCCCTGCTTCCCTTGGAAAATTAAGCCAGCAGTTTGTGTTGGGGAAGACGAA
GCTGCCGCAAGCAGGACACTGCTTTTCCTGACCTGGTGCCACACCACTGGCCCCTGAACACACAAACCATTGGCC
CTGCTCCTTAGCATTGCCTAGCACTGCATTCAGGCCAATTCCTCTCCCTCCCAGAATCACTCAAGGACCCCTATT
TCAGCAT

**>Mouse Gja5-7**
GTCACAAAGGTCCTTTCCATTCACAGAGTCTTGCCTGCTCAACAAACTTTCAGGTGCCTTTAAAGTTCTTAAGAA
AGGGGCCTGACTGAGGGCATTGTGCTCTGGAGCTGGCCAGCTTCGCCCTGCCCTGGCCCCAGCACCCCAACTTGT
AAAGAAGGAATTGCGAGGTTATCTAAAAGGAAGCCAAATTGTTTTCGGAAAACAATAGCCAAGTAGCTACAAGTC
ATTAGTTTCCTCCCAGGGGCCTAAAATCAAGCTTGTCTGGATAACTGAAAGGGCTGTTAGCACTGTTTACTTAAA
GACCTCTGGACAGAACCCATAAATTAATTCCTGTGCCTGGCTTTCTGGAAGAGGGAGGGGGTGAGTCATGGAAGT
GGGATCATGCCCAAGATTCAAGTGATTCCTGTTTCTCTAGCTGAGTCCCTGCGGCTGGACCTGCCTCTTCTGGTT
TTCTCCATGGAAGTTTTTCTTTTAGAAGAGTGTGGGG

**>Mouse Gja5-21**
AGTAAGGGGAGGAGAGGAGGGAGGAAGAGGAGAAATCCCAGGAGGATATTGCTTCAGGGGTGTGGAGGAAGGGAG
TGGGAGTCTCACCTGAATCTATGAGGGGTGAATGAACAGGCAGTTCCCCAAAACCCTTGTCTCCCCCAGAGCTAT
GGCAACTGAGTTATGCTGAGCTGTGACTCTTTGTTCTCAAGAATGTGATTCAGAGCAAAGGAGACGAAACGCCAC
CGAGGAGGAAATGTGCCCAGGTCCCTCGCCTGCCCGGTCGGTTGGATGTGCTATCAGCCCAGAGCCCACTGCAAC
TTCCGCTGCTATTGTTTAGCATTTACCCTCTCACATAACTGGGTTCTGGGAGCCACTGCCCAGAGGACCTAGGCC
ACCACGGGCAGGAGTGTGCTCTGTGAGGGTCTGTGGTCTGAGGTTACCCCAGTCTGCATCAAGTTC

**>Mouse Gja5-28**
GCACACTCCATCGATATTTAGCTCATTATGTTTAGCATTTGTCTCTCACCACTAGATTGCAGGCTCATGTGACTG
AAAATTAAGTAGTCTTGGTTTACTGCTAGATCTCCAGCCCTAAGAGCAGCGGCTGGTAAATGCTAAGAACTCAGA
GATTGGCTAAATGAATTAATCATCTTCCTCTTGGCCACCACCTTTCCAGTCCTCTCAGCTCTGTTGCTTTTGGAA
GCTCTTTTTTCTTCTGAATCACACTTTGATTTGCCTAAGATGTTGAACCACCCTGTTCCTCCATCAGCACCACTT
CCCCTGGAGTAAGATGTGAACGCTGATTATGCTTGCCCTGGAGTCAGACAGTACCTTACCTTCAACCCCATTATC
AGTACCTTTTCGTGAGCTGGTTGATAAGATTTCCATAAGGATTGGCGAGTACCTATCACAGTTCTTCACTCTACA
GGAAAGAGGGTGAAGCTCAAAAGCACTAAAGGAACATGCCAAGGTCTCACAGGCAGAAAGCTCAGGAGTCAGGAT
TCAGACGCTGAACCATCTAAGCTCAGAGCCTACAGTCAGCTGCCAAATAGTGTAGCTGTTCTCAACGGGTGTTGC
CCCTGAAGAGTTACTGCGATTGCTTCACAGTAGAAATTCACAGCCAACATTCGCCAAGCAGGTT

**>Mouse Gja5-78**
GGAGGGGTGCAAGAGTGAAGCTCTCCACGTCCACATTCCAAGATTCCCAAGGGAAACTGCATAAACATAGTAGTT
AGAAGGGACTGAGCCATCGCTGCCTAGCTCGGATGGGAGCCTAAGGGTTCCCATGGTAGCTATATCTGGGCCCCA
GGGCCGACTGGCAGTGTCTGTACAATGTCCCTGCTTACAGCAGAAAGAGGACGTAAGGTCCAGGGTTCCCAAGTT
CAATGGTCTTTTCTTTTGTTGCTGTAGAAAGATTGCTGAGGCATGAGCTACTTCCGTCCTCCCTTGGCATCTTTG
CACTGTGCCCCAAGAAACAATCATATCACATTCAGGAAGGCTGGGCCAGCAATGGCAGGGAAACAGGAACTGTCC
CCACGGATGTCCTCCCCAGGGGAGACAGCCAGGGACTGGGAACAAATACCGTGTGAAAAGGAAAGCCAGACTCCA
GGTTGAGGAGCCAGCCCTTCCCAGCATTCAGCACGACCCTCTGCCGTGAAGTGCAGGAAAGGAACGTTATTCTTA
ATGCACGGACTTTTCCTTCCCTACCCCCACCCTCTTTGCGCACTCCCTGATTCCTGTGAGAAACTGGGCTATAGA
ACACATTCCAATGCCTTTGCCCAGCCAGTCCATTTCCCTCTAATAAATTCCACAAGCCGAGCTA

**>Mouse Gja5-93**
AGGGGATTCAGTGCTATCTTCTGGCCTCCATAAGCCTCGGGTATGCCCCCAGTGCCCTGACATGCAGGTGAAGCA
CTCACACACATAACATAAAAATAAATCTATCTTAAAAAAAAAGATCCAACAGGTCACATTGGAGCATTGTAGGTA
GTTATATATCTTATCTGTGGGTAAAACGGGCTGAAAAGGTAGCGGGAGTGAGGCGGTGCCATTTACAGAAGCCAT
TGAACAGCCAGAAATCTAAGGAACTCATTTCAGCATGCCATTACCTCCTGCCTGGGTAATCGAGCAAAGAATCAG
CCTGGACCATTGGTGAGCCTTGACCACGAGGGGGTGCTGCTGTTCCGGCTTTAGAGCCTGAGAGCATCAGGGGAA
CTCTGTCACACCAGAATAATGGGCGTGGAAATGAAGGGACAGGGAACGGCAGGCAGGAAGATGGGGTGTGCGTGT
GTAACTTTTTATTCGGTGGCCACAATTATTCTGGTCTTCTAATGTCCTTCTCACTATCCTGGTGGGGTTAGGTGA
CCCTCAGGGATGGTTGTTGCTGTTGTTGGTAGTGGTGGTGGTGGTTGTCGTCGTCGTCATCTTCCTCCTCCTCTC
CCTCCTCCTTTTCCCTCCTCCTCCTCCCTCAAAAGCCCCTCTCATAACCGTGATCTTTGTCCTACCATCTGTTGG
TCTCAGTCCCAAATGACCAAATCC

**>Mouse Nrp1+76**

GTAACCATCACACCACTACCCAAGGCCCTGTGCTTGGTTTCATGGAGTCTTATGTGAAAATTTTGCTTATATGCC
AATCAATTGGTGGAAGAGTAGGCAGAGCCTGTCTTTCATTTATTGCCTGATATATAAAAGTGACTAACTCCCAGG
GAGAAGAGTCTCCATGCAAGAAGAAAGTGAGATGGTTTATCCTCTCTTCATGTCTGTTGAAGTTTCAGTCATTTG
AGCACAAATATAGTATTCTGTTTTACTTTGCAAAGATCAATATTAGTTTAACATACACTTAAGTGAAAGAGCCCA
TTACAGGAAAGTGAACTCATGCTAACCTCTGAATGATAACACTGAGCCATGTGGAGAGGGTGGAAAAGATGAGG
TCTCACTCGGGATGGCTGAAGCCTGTGCTAGATGAAGCCAGAGGTGAAGTTAATTTGCTCACGGTAGCTTGTCTT
TCCATGTCTCCAAATCCAAAGCAGCTCCCTTTGAGTGACAGACACAGCTCTGAAGGAAGCTTGCATTTTTGCAGT
GCTAAGCATGTCTGCATGGTAATGCACTTCTATTCTGGGTAACATAGACATTACTCAATAACAGAGACCCTGTGG
TCCTGCAGGGGTTCTGAAACTGCCATAGTCGCACTTCTTCTGCGTGAAACAGTTCTTCATTGGCAGAAATACAGA
GACAAAATGGGCATTCCCAAAGCACTTATCAATGGGATGGGATCTTAAGAATGCTCTCCAAACCTTTCTCTTGTG
GGATAGTGGAGCCAGGAGAAACATGGAAATAACACTCCCCAGGATTGTTTCTATTTATATTCTTCCACCACTGGA
CTGCCTGAGTGCAGGGGGTCAGACCTTTACATACCGGGCTGAACAAGGGGTGGCTGTTATCTGAAAGGCAGAAGA

**>Mouse Nrp1+78**

TGGGGGAACAGTGTTATAGTGTGCTCAACTGTGTTCACTGACAAGAGTTTTTTTTTTGTGAAGTGGCCTTGAGCA
GGGTGTGGCCATTGGAGAACCTTCCAGTCGGGACTTCCCTACATGTTCCGTCTTACCAGGCGGCTCCTTCTCTTG
TTTTGAAGCTTCCTGCAGACAACAGATCTCACAGGAAGTGCCAGTGTGGACGTTCATATCCATGGCTAACGATGC
ATTGTTGTTTTTTTTTTTAAATGTTACTTGGTAATTTCCTAGAGAAAACAATGGGAACCGAAGGCTGTTCCTGTC
TTTAAGGAAAGCCAGTTTCCAATGTTGCTTTATTAACTTAACACATGCAAGTATTTGTTCTAGTTGTTGCTACAG
CAACGTGTTTTGTGGGGGCATATTTCATGCAAAATGGGATAAAGCTGAAAATCAAGACAAATGTCATCTGCATAA
TCATGTGCATAAAAATAGAAGTGTACGTGTGTACTGCTGTACGTGCTATGCTGCGTGATTAGCAGCGCCTGCCT

**>Mouse Nrp1+91**

AAAGGCCCACATTTTACTGAGGATCACTGGAGGACGAAGCAGCATTTCTCAGCATTGTATAGTAAATCACACAGC
TCGTAGCCTTTGAGCCATGAAATGCTTGCCGCTGCTTGCCATGACTGCGCACAACCGGCCATTGGTAATGTAATG
TGGAAAACCTCATTTCCAAAATGAAAATCTGCCTTGCTCAATAAAATACGGCCCCGTGAGTGAGCATTGGCAAGG
GAGCATTTATCAGCTTTTAATGCTCCCCTGGCAGGCCAGCACAACAGCCTGAGAAAATAAAGGACAAAGAACTGA
GGTCTGATTAAAAACCGGAAAGTCTGCTTGTGAGGGGAAGACATGGCAGGTTTTCTTCAGAACATCTCTGATGAC
TCAAGGCTGTTTAATTCATCAGTTGATACTAAAGCTAAAGAAAATATTTTTAAGTAATTGTCTCATTTCCTTAGA
TAAAACACATGTTTTCTGACCTAGCCAGGCATGTAGACCTCAGCTATTCTAGGCTGAGTGGGCTGATTCACAGCA
TCTGACCATTGTCTGCTTGAAACTCAGGATCTCTGGCTTTTTATAGATGCTAATCAGAACACAACCACCTAAGGT

**>Mouse Nrp1+129**

CAGGAACACAGCATTTGGGGGATTTCCTTTGCCTAACGTCTGAGGGGTTTGGGATGCGGGAGCAAGCGTTTTGTC
AGCTGAACTCCTGAAAACTGGGGGAAAGTTCCAAAGAAGTTTGTCTTACAGTTTAATTAGGTCAATGGTAACAGT
GGCCTTTAAGGAAGCATGCTAATTAGGGAAGTCCTTTCTAGTCCTCTGTGAGGCAGTCAGTGTGGAGGAAACACC
GATACACACCACACGGTCCATTAATAAATGACTGCAGAGGCCAAGGCTGAGTGAACTTCAATAAAGGAAATGGAA
GAATCTGGGAAGCTGACCCTCTCACCTTCCTCTAATGATTAACTCTAAAGTATTTAAAAAAAAAAgaggtAAAAA
ACCTCTGGTTGCTTATACAACTGCTTCCCTTCATGTGAGTCTGTGGGGAAGAATGTCACAGAATACTAACAGAAA
ACAAAAGAGGGCAGCCTTTGATAACCCGCACGCACGGTTGGGCGGACAGAGTCCTTGGGTTTTCCTCTGAGCTAA
AGGATAAGGGGGATGCTCCCAGGGTGTTTCTCAGCCCGGGCACAGTGTAAACCTTGGTTTCCTGAGCCAGTCTTG
TGGTAAGTGCTGTTGTAAAACTGCCAACTCTGTGGCAGGCACAACAGACAAACTTTGCGCTTCTTGATTTTGCAG
AATGGTGATGAAAATAAATAGTTAGGCTGTGGCTTGTGAGATTTCTTTCCTTCTTCTAGTTTGCAAACACAGGCC
ACATCAGAGCT

**>Mouse Unc5b-57**

AGCTGTCCCGGACCACCCACAGGAACTTCTAGAGGCCCTACCCTCCAATGGCGCCCCCCAAACCTTTATTAGGTC
CCCGATCCCCCTGAATAACCAAATGCAACTTCAAAGTCAGCCATACCATGCGTGTCTTCCATCCTGGGCTTCCCT
TCCTGGGATGGAGAGGCCAAGCCATATGACACAATCAGGTTGATGGGATCCTCGCAGCCTTCAAAAGTCACTTGC
CAGAGAGTCTTCTGAATGACACACACTTAGGTCCCTGGTCTCTGCCTGTCTATTGTTTCTCCATGCCAGCTTCGG
GTTTCTAGAAACTTCCACTGCACACATTGTCCTGAGCGATCCATCAGCTCAGTTTTGTTTACTAGGCTCACTTCC
GACCCTGCACTCAGTCTGACCCCGGAACACACCACCTTCAGCCACTTCCCACCACCGGGTCACTTCCTGGCCCAC
TCAGCAGAAGAGCGCCAGTCCTTCTTCCACTCTTCCCACCCTGGGGGCACCTGCTTCCTGCTCTTCCTCTGCCTC
TGTGCCCTAAGCTCAGGGTCCACTCGCAGCTCAGGGGTCACTTCCTTGTGACCTCACCCCATCTGTTTGTCAGGA
GACAAAGCAGGGGTTTCAGGATGGCTTCCTGAATCTGGTGCCTCAGGCTTGGCTGAACTTGATTTTTTTCCACAC
AGAATAAAGATACTCTCAGGTGGTCTCTGCATATTATATCTATTATTTTAGCAACTTTTGCCATCCCCAAATCCA
CATAAGTAGTCCTTGTGTGTTAAGTGACTGGAAGAGGTCACATTCAGGTGGGACAAGAGAGAACCCTTCCCCCTT
CCATCAACAGCCCCTGCAGGTCACATAGCTTTGGATATCAGATTTTTGCTGCCTAGACCTGGGTCAGCCCAGTGT
A

**>Mouse Unc5b+14**
CTCCCTCACCAAGAGTCTCTTCCTAACAGACTGAAGACGTGTAAATGACCCAGTGCATTGCTGGGAGTGCCAACA
GGCGGAAGTGGTGGTGACATCATGCGCCCATCCGGCTCCGAGCCTTCCAGCAGCCTGAGGTCAGCTCACAGAGCC
CTGGCACCTCCCCCAGCAGCCCCACACCGAGCCAAGAGGAGCCTGGCACTGCCAGCGCTGGGGCTGGGCTGACAG
CTTGGAGTCCCTCCTTCCTCACGTCCCCTCTCTGTGATGCTTCCCCCCCCACACACACACACTCCAACCCCCACC
ACCCCAATCGCATCCCTGAACAGCATTCACCAGCAAACGTGAGATTGCTGCTTCTCATATAACACATGAGCTGTG
GTGTGAACTTCTGGTAAATGGAGATCTTATTTAAACTGCGTGCTATCAGGGGAGAAGAGACAGGTTCCAACACAG
CTCAGCTCTGTTGCCAGAAAAAAGGAGATGCCGGCACTGCTGGAGCCCAGAGCCATTGGTGTCTCTCAGGCACCA

**>Mouse Unc5b+23**
AGGCAGAGCTGAGCTCTGAACGTGAAGGGAGGTAAGACCGTCTGGGAGCTTCTGGCTTGAGTCTGGGTAGCACTG
GGAATCCACACTGTGAAAAAGCTTACAGGTGACAAGGACAGAGATGCTCTAAGGAGGTCCCAGCTGAATTCCAGC
GTCCCTCCGCTTTCCCAAGCACCAGCTTCCCAGAGAGCTAGAGTCCTGGACCGCAAGCACATACACACCATGTGG
TATAAAAAGAACTCTTTTCAAAAGCAGTCCCCAAGGGTGGGGGGAAGGGAGGTCGATGAGAAAGGAGCCAGGGGG
ACAATAGTGAATAGAGTTAGGATTATTTCCTGAACTCTGACTCCTCGTGGGAACGAGGTTATGAAAAGCTTTTCT
TAAAATGGTTATTATTTAGTGGGCTACCCAAAATAAACACACACATCCATCGCACGGAGCCAAGCAGGAAGCGAG
GAACATCTGGGAGACATCTGGTTGCTGGAGAAGAGGGAAGTGGTGGGAAGCTGGCCCGGGGAGGAGCTGGCAGGA
GCCGGACGTCTGCATGCAGTGGTCACCCAGAGTTACACACACTGCCACGCCAGCACGCCAAGTCTGTGGCCCAGC
TTGGAACACGAATCACACGCACGGAC

**>Mouse Unc5b+30**
GGGAGGGAAGGCGGCCGCGCTTATCTGAACAATGTAAACAAACAAGCAAATAAACCTTTTCCTGGACTCCCAGAT
ACTTGTGAGGGCAGCAGAGCCTGGAGCTGTTTCAAAGGACCCAGGAATGCTGGGAAGTGAGCTTCCCCAGTGAGT
GGCTCGCTGGGCAGCCTGTGGGAGGGAGCTCAGGGGCTACAGCTAACCATCAGGGGCTGCACCTGCTCCCACAGG
AAGAGTACCTCTGGGGACCCTCATTGTTCTCTGGCTAGGAGGAAAGTGAGGGGCGAGGAACAGAGCCCACTCCGA
AGCCAGGAAATAAATCTTCCCTTCATTCAGGTCAGCTCATTTAATGGGGACCCGGGGGTCTGAGGACTCGTGAGA
AAAATCTCCCCAAGCTGGGCCCACGCCATTGAGTAGACTCCAGGGGCCCATCCCAGAGGTTTCCTTTGGCTATTT
CCTGGCATTGTTGACCCCATAATCCCAGCAGCAGCAGGACATGCTACCTAGCATCAGCAGACTGTTGGGACCCAG
GTCTTCCCAGACCTTGTGAGATCTGATAATCTGTGAAAGAGGAAACCCCCACCCCACCCCCTCTTCTCCATCAGC
TCCACTTGGACACCTTACTCCCCAGGTGCTGGTTAGTAAGAAGCCATT

**>Mouse Unc5b+39**
AATGTCCTTAACTTCCTCCCAGTTTAACACCTGCCCCTCATTAAGCCCTGGGCCCAGGGGGTGGGGGGGTGGGGG
AGAGTGTCTCAAAGACAGGGACTAGAGGCTGTGGCCTTGGGTAGCCCAGCAGATTCCCTCTCAGGCTGTCCACTC
TCCGGTCCTTTTATTGGACATTTCTGAGGTTCAAGAGTGGCTCATATCATCCGGGAGAGGGACAGCCACCAACTT
CCTCAAGTCCCCTGGGATCCAAAGTTCAAAACAATAGCTGTGTCGGTGGCGGCGGGTTGCCCTTGATGCAGAGGT
TCGGTGGGTGGCTAATTTTAGCTGAGGACTACAGCTTTTCCCAGGCACGTCACTGGCCACTGTCTCAGCCCTGGC
CCTGGACAAAAAGGTCTCGCTTTTCCTCCCCAGCTGGGAGCTGAGGACTGCCTGGCGCTCAGGACAATTCCCTTT
GTGCCTGCTTCCGGGTGGCTCTTAAAGCTCAGAGAATGGTCAGCCCTGAGCTCTCCGCCCCAGCTCTCTGGCCCC
TCTCAGTCTCTATTGCTTCAAGGTCCCCTTATTGTTATTCCTGAGAGCTGGCATTGGTGGGGAAAGAGTAGCAAG
TTCGGGGTGAGGGACTGATGATGTCTAGCCGCAGTTCAGAGAGGGTGTGAGTGGCCTGAGGTCACAGAGCACT

**>Mouse Unc5b+43**
CTTCCCAGAGGAATCAGCCTAAGTGCACTATTCTCTTTCCTTCCTTCAAAAGTCTCTCTGTAGACCAGGCTGCCC
TCAGATCTGATCCTCCTGCCCCAGCCTCCTGAGCGCTTGAGTCACAGGCCCATGTCACCAGCCCCACTCTTGTCA
CTTTATGAAGAGAGAGTAGCTTTGCCTCCCCTGAGGTGACCTTTGACCCCAGGCTGGGGCTGAGCAGGAGTGGGC
AGGTTGTCCAGAGCTCTCTAGCCTGGTTCAACTCCTCAGTCTGAATTAGATCTGAGAAAGTGGTTTGCCTCTTGC
CTGCTAACTTGAGGGTGTTGCCATGAGGGTTGTCACCCAAGGAACCGGATGACAGTGTGCCCTGAGAGTCCTGCA
TGTTACCAGTCTCAGAAATGGGGAGCAACTGACAACATCACACAACACACTAACCCAGCCGGATCTGAGTTCTGT
TTCTGGCTCAGGGCTCAAGAGAAAGGTGCTTGGCTGTGGCGGGTACACACCCAGTGTCCCACATCGGGCTATGAA
CGCTTACCATTCCTTTCTGAGGCCGGGCTTTGCTCCCAGGGTAGAGGTCCCAGGACAAAGCTGCAGCTTGGCAGG
GAGGGTCTAGGCATATCCATCACCTGCCTTTATCCCTGAGGCCGAGGGATCCTGGCACGGTTCTAGGAATCTCTC
GACAGCAGAGCAGGACCATGTGTCCAGGGCTGGCTGGCACAGGACAGGGAGGCGAGGAGGGGGTAAGGGAAGTGT
GCGCAGAGTTGTCCCACGCTGGGCACATTTATCATCAGCAAGCACAGTGGCTCTGGACACCGGCTAATTCTCCCA
GCCCTGACTCCCACTGCATGGAAAATCTCTCCCATTCACTGCGGGCCGCTGTGCTCGGTGGGGAGCGGGCAGAGA
TGGAGGAGATGAGCCAACTTATTATTATGTTTTAAAAACACAAACCCCACAGGCTGGGCCCCTCCTTGGACACAG
AACACTGCCTCCATCTGGCCCTAGATCCTGTGGGGGTTCAGGCTGTGGTCCCCCCTCAGAGGCTGTTTCAATGCT
CGTTACTGATGACAAATGGGGGGCTGGCCAGCTTGGAGGACACCAAGTTGCTCCTAACAAAGGAAGCCTCTCACC
CAGCCCTGTTCCCACACCCTCCACAGACCCATCCGTGCCCATGATGTCATTTGCTGTGATGGCCACAGGAAGTAG
CCTTGTCAAGAGGGCCAAACCTAAGGAGGGTCCAGATGCCCACATCCCTCCAAGGCCCAGACATCAGGCTAGAGT
AGCCCATGCTACTTACAAGAGTGGGAATGGACACACAGTCCTTGGGTCACTGGGTCACTGGGTGACCCG
